# On the intra-laboratory replicability of results in animal cognition research, obtained in animals that were not exposed to, or unaffected by, experimental manipulations — Exemplified by spatial learning in pigs *(Sus Scrofa Domesticus)* across 12 holeboard studies

**DOI:** 10.64898/2026.04.19.718108

**Authors:** F. Josef van der Staay, Alexandra Antonides, Alexandra K. Dwulit, Lisa Fijn, Elise T. Gieling, Charlotte G. E. Grimberg-Henrici, Ellen Meijer, Sanne Roelofs, J.C.M. (Hans) Vernooij, Vivian L. Witjes, Saskia S. Arndt

## Abstract

The replicability of experimental results is considered a cornerstone of empirical research. However, the reproducibility of results from groups that have not undergone experimental manipulation — the standard against which comparisons in an experiment are made — has been almost entirely neglected in animal research. Our aim is to address this gap by exemplarily determining within-laboratory replicability using research in pigs, an increasingly popular large animal model species.

Drawing on data from twelve independent porcine holeboard studies conducted in our laboratory, we examine the replicability of groups that were not subjected to experimental manipulation (typically the control group), eventually including groups on which the experimental treatments had no effect. These analyses were also performed on all eight studies involving the Terra x Finnish Landrace x Duroc pig breed, with all other breeds excluded to increase genetic uniformity.

The holeboard is a complex spatial discrimination task in which an animal must learn to find food at four of sixteen possible locations (holes) arranged in a 4 x 4 matrix. The main variables measured are spatial working memory (WM), reference memory (RM) and the inter-visit interval (IVI), which serve as an index of motivation. All studies showed predominantly linear improvements in WM and learning rates across successive trialblocks. IVI showed greater variation across trialblocks, but this did not affect WM and RM learning, which are robust and replicable indices of spatial learning in pigs.

Assessing replicability provides relevant information, such as whether behavioural and physiological traits in a model species are stably expressed and robust across studies. Including replicability research and publishing its results can stimulate the development and use of more replicable methods and workflows, thereby increasing scientific rigor. Provided the data are available and accessible, the next step should be to expand replicability studies to include those conducted in different laboratories.

## Introduction

Awareness is growing that research areas such as psychology and medicine, as well as more “exact” sciences such as biology, physics, chemistry, engineering and earth sciences, are confronted with the problem of non-replicable research results, albeit to varying degrees (Baker 2016). Replication studies are important for scientific advancement, as they form one of the pillars of science (Kelly 2006; Macleod and Mohan 2020; Park 2004). According to a questionnaire by Farrar et al. (2021), in which researchers specializing in animal cognition were asked about the prevalence and importance of replication studies, 70.0% disagreed that enough replication studies had been conducted in their field and 74.1% disagreed that sufficient replication studies had been performed in animal cognition research in general. However, 79.8% agreed that it is important to perform replication studies in this field (percentages from Fig. 15 in Farrar *et al*. 2021). Experimental results should be considered as preliminary if they have not been corroborated (van der Staay *et al*. 2009, 2010).

Although concerns about the replicability of study results have been discussed for decades (e.g., Smith 1970; Ioannidis 2005), the discussion about replicability has increased since the narrative of ‘science in crisis’ gained growing attention, and since this phenomenon was labelled a ‘replication crisis’ (Maxwell *et al*. 2015), or ‘reproducibility crisis’ (Fanelli, Daniele 2018). Despite growing awareness of the reproducibility problem – which threatens the credibility of scientific results and public trust in science (Anvari and Lakens 2018; Wingen *et al*. 2020) – and various initiatives to promote replication efforts (see e.g., Derksen *et al*. 2024; Mullane and Williams 2017; Open Science Collaboration and Nosek 2012), the number of replication studies remains limited (Cobey *et al*. 2023). The terms ‘replicability’ and ‘reproducibility’ are not clearly distinguished and are sometimes used interchangeably. Following definitions proposed by Bollen et al, “reproducibility refers to the ability of a researcher to duplicate the results of a prior study using the same materials and procedures as were used by the original investigator” (Bollen *et al*. 2015, p. 3), whereas “replicability refers to the ability of a researcher to duplicate the results of a prior study if the same procedures are followed but new data are collected” (Bollen *et al*. 2015, p. 4). A slightly different definition states that “replicability is obtaining consistent results across studies aimed at answering the same scientific question, each of which has obtained its own data“ (Committee on science, engineering, medicine, and public policy 2019, p. 46). In the following, we use the terms ‘replicability’ and ‘replicability study’.

Currently, there is no consensus on how to classify replication studies. Various classifications and taxonomies coexist (e.g., Baldassarre *et al*. 2014; Bettis *et al*. 2016; Greulich and Brendel 2019; Hüffmeier *et al*. 2016; Matarese 2022; Schmidt 2009). We found the following division useful: original study, exact replication and extended replication. The latter is subdivided into systematic and differential replication, conceptual replication, and quasi-replication (van der Staay *et al*. 2010; see S1 – Supporting Information – Classification of Replication Studies for more details).

Most replicability studies are designed as **inter-laboratory studies** that aim to establish the robustness of results from experimental interventions. Robust scientific findings are characterized as reproducible, replicable, and generalizable (Bollen *et al*. 2015). While studies on inter-laboratory replicability of experimental results have been published extensively, studies on **intra-laboratory replicability** are lacking. Rather than focusing on the robustness of the effects of experimental interventions, the focus could be on replicating data from groups that have not been subjected to such interventions. Groups that have been subjected to such interventions but appeared unaffected, and that have been tested in highly similar conditions using the same equipment, could also be included. To our knowledge, this issue has not yet been explicitly addressed. Eventually, replicability studies could be expanded to include groups tested under control conditions at the start of the study, before any manipulations are applied. Additionally, groups that are unaffected by the manipulations, as indicated by a failure to detect treatment effects through appropriate statistical analyses, could be included to increase the sample sizes of a replicability study. This approach can be advantageous if group sizes within studies are small, and the number of studies available for analysis is restricted. Since control groups provide the baseline (Schickore 2025), or reference point (Torday and Baluška 2019), against which the effects of experimental manipulations are estimated, this type of analysis could offer valuable insights into the animal models, as well as the materials and methods used in the experiments.

For example, for less frequently used animal models, such as large animal models (Dwulit *et al*. 2025; van der Staay *et al*. 2017), or when assessing treatment effects using a newly identified phenotype or genetically modified strain of laboratory animals, relevant questions may include: How replicable are the expressed phenotypes (e.g., in learning and memory tasks)? And how robust are the materials and methods used to test them across experiments? Answering these questions and confirming replicability is also relevant for laboratories that use the same control condition in experiments with different scientific questions. Assessing the replicability of effects and estimating effect sizes in this case would not make sense.

We want to draw attention to the replicability of intra-laboratory controls and groups that are unaffected by experimental manipulations, and address the replicability of baseline (control) measurements. To this end, we analysed the data of 12 independent holeboard (HB) studies with pigs as subjects, all performed in our laboratory. Compared to rodents, pigs are a far less commonly used animal model species, though they are growing in popularity as large animal models.

We have performed a series of experiments with pigs as subjects, assessing their spatial orientation learning in a holeboard task and the effects of different experimental manipulations (see S2-Supporting Information, Table S2.1). Holeboard-type tasks have been used in many previous studies, most of which have used rodents as subjects. A wide range of testing arenas and procedures have been developed (van der Staay *et al*. 2012). We used a holeboard version and testing procedure close to that originally developed by Oades and Isaacson (1978) for testing rodents, adapted for assessing spatial cognition in pigs.

The holeboard is one of the tasks used in translational biobehavioural and neurotrauma research involving pigs (Netzley and Pelled 2023). To date, the number of holeboard studies involving pigs remains low. While different laboratories have conducted holeboard studies in pigs, the largest number of these studies originate from our laboratory.

### The spatial holeboard discrimination task

The first studies on the holeboard task were done in rodents in the late 1970s to early 1980s to assess spatial discrimination learning (Oades 1979, 1981; Oades and Isaacson 1978). Since then, a large number of variants of the holeboard task have been developed and tested on a wide range of species (van der Staay *et al*. 2012). Over eight years, we have conducted 12 studies assessing spatial holeboard discrimination learning in pigs (see S2-Supporting Information, Table S2.1). The holeboard task has also been used by other research groups to assess the effects of experimental manipulations on spatial learning and memory in pigs (Arts *et al*. 2009; Bolhuis *et al*. 2013; Clouard *et al*. 2016, 2021; Galvagnon *et al*. 2026; Gautier *et al*. 2018; Haagensen *et al*. 2013a, 2013b; Val-Laillet *et al*. 2017). In addition to studying the effects of different experimental conditions on cognitive functions, performance in the holeboard task has been used as a behavioural proxy for the effects of these conditions on pig welfare (Ede and Parsons 2023).

Holeboard tasks enable a large number of variables to be assessed simultaneously (van der Staay *et al*. 2012). The most important of these are spatial working memory (WM), spatial reference memory (RM) (Olton *et al*. 1979; Olton and Samuelson 1976), and the inter-visit interval (IVI) (van der Staay *et al*. 2012).

In reference to the HB task, pigs use **working memory** to store the list of the holes visited during a trial. This list is trial-dependent, as it is erased and reset once the trial has finished. In contrast, pigs use **reference memory** to store information about the solution to the spatial holeboard discrimination task that remains relevant across many trials of an experiment and is therefore trial-independent. This includes general task rules, such as the location of the food and the necessary actions to obtain the hidden bait (e.g. pushing up a ball to uncover the food trough containing the food reward) (for more detailed definitions, see van der Staay *et al*. 2012). The time the pig spends per hole in the holeboard task – the **inter-visit interval** – can be used as an index of motivation to perform the food-search behaviour (Witjes *et al*. 2025). For instance, pigs with lower food motivation (due to milder food deprivation) took longer to run down a runway to a food source than pigs with an increased level of food deprivation (Hsia 2004).

### Aim of the study

The present study aims to demonstrate the feasibility of replicability analyses within a laboratory or research facility for control groups and groups unaffected by experimental manipulations. This is demonstrated through a worked example of 12 independent studies performed in our laboratory, in which pigs performed a spatial holeboard task. All holeboard (H) studies whose results were used in the present analyses have been previously published in peer-reviewed scientific journals [H01–H12: H01, (Gieling *et al*. 2012); H02, (Gieling *et al*. 2013); H03, (Gieling *et al*. 2014); H04, (Antonides *et al*. 2015b); H05, (Antonides *et al*. 2015a); H06, (Antonides *et al*. 2016); H07, (Fijn *et al*. 2016); H08, (Grimberg-Henrici *et al*. 2016); H09, (van der Staay *et al*. 2016); H10, (Roelofs *et al*. 2017b); H11, (Roelofs *et al*. 2018); H12, (Witjes *et al*. 2025)].

We hypothesized that the learning curves of pigs in groups not exposed to or affected by the experimental manipulations would be similar, as indicated by comparable initial performance levels and rates of learning. This was expected because identical or very similar procedures and learning equipment were used in all twelve studies. Finally, we will discuss the limitations and opportunities of this approach to replicability studies.

## Materials and Methods

### Ethics statement

All of the studies were conducted in strict accordance with the recommendations of the National Institutes of Health’s Guide for the Care and Use of Laboratory Animals. Each of these studies was approved by the local ethics committee (DEC, DierExperimenten Commissie) or by the local Animal Welfare Body at Utrecht University, The Netherlands. The corresponding ethics statements can be found in the original publications.

### Inclusion and exclusion criteria

It was not straightforward to determine which groups of pigs should be included in the replicability analysis. We ran our analysis using two sets of different exclusion/inclusion criteria. The first set included untreated pigs categorized according to criteria such as birth weight (low versus normal), sex (male versus female), breed (e.g. conventional versus minipig), housing conditions (barren versus enriched), litter size (small versus large), and birth order (first versus last) (see S2 – Supporting Information, Table 2.3 for details). Some of these studies involved pigs that had undergone experimental manipulations (e.g. prenatal allopurinol treatment or diet-induced iron deficiency). Data from the experimental groups were only excluded from the analyses if they had an impact on HB learning in the original study. In study H04, for example, the experimental group developed anaemia after being fed an iron-deficient diet; in this case, only the data from the control group were used for the replicability analyses. Where there was no difference in learning the holeboard task between these subgroups in the original study, their data were pooled.

The second set of exclusion/inclusion criteria was identical to the first, but also excluded pigs of any breed other than Terra X Finnish Landrace X Duroc (see Supporting Information, Table S2.1). This left eight homogeneous studies with respect to the genetic background of the tested pigs.

Although we performed replicability analyses based on two different sets of exclusion/inclusion criteria, other choices of groups of pigs are also conceivable for this analysis. To exclude, for example, putative effects of experimental manipulations on the pigs selected for the replicability study, we performed an additional analysis on a selection of pigs, using a third set of strict exclusion and inclusion criteria. Details of this additional analysis and its results can be found in ‘S3-Supporting Information-Alternative exclusion and inclusion criteria for selecting pigs per study for the replicability analyses’. We will take the results of this additional analysis into account in the discussion.

Additional information about all studies can be found in S2-Supporting Information, Tables S2.2 – S2.3, which provide details such as breed, post-weaning housing conditions, training and testing methods, and composition of the groups of pigs selected for the present analysis. Full details of materials and methods can be found in the original publications.

### Holeboard apparatus

Two different apparatuses were used: one holeboard with a large arena, measuring 5.3m X 5.3m (Fig. 1A left), and a holeboard with a smaller arena, measuring 3.6m X 3.6m (Fig. 1A right) surrounded by a corridor. A guillotine door could be opened in the centre of each of the four sides of the arena, allowing the pig to enter. Food bowls serving as the ‘holes’ of the holeboard were distributed within the arenas in a 4 X 4 matrix (Fig. 1C,D). The guillotine doors and holes were identical in size and construction for both holeboards, as was the width of the surrounding corridor. However, the distance between holes was shorter in the smaller holeboard.

All holes had a false bottom underneath in which rewards could be placed to control for odour cues. To render identification of the baited holes by visual cues impossible, a large and hard plastic ball (24cm diameter, weight 1400g) covered each food bowl. A pig could easily raise the ball off the food bowl using its snout to gain access to rewards underneath (Fig. 1B). Guide rails ensured that the ball could not be knocked off the bowl and that it returned to cover the bowl once the pig had retracted its snout. Rewards used were chocolate M&M’s^®^ (Mars Nederland b.v., Veghel, The Netherlands). More detailed illustrations and descriptions of the construction of “holes” can be found in Antonides et al (2015b) and Roelofs et al (2017b).

Lifting the ball interrupted the connection between a magnet in the ball and a sensor in the food bowl. This interruption was registered by an interface (LabJack) and sent to a PC. In all studies except H07, hole visits were automatically recorded using custom written software (SeaState5, Delft, The Netherlands). In H07, hole visits were scored manually as the automatic system failed to register the hole visits. Melendez et al (2013) used J-Watcher version 1.0 (Blumstein and Daniel 2007) to assess the intra-rater reliability of manually recording hole visits from a monitor compared to automated registrations; WM and RM scores were calculated based on manual and automated recordings of hole visits. The correlations between the scores obtained by the two methods were very high (product-moment correlation coefficients ranging from 0.928 to 0.995), and the kappa coefficient of 0.8 indicated high intra-rater reliability.

Although different patterns of baited holes were used in our studies, these patterns were selected according to the rule outlined in van der Staay et al (2012). This ensured that the number of holes in the corners, periphery, and centre of the arena was equal across studies. The location of the reward is a relevant aspect of this task because it has been shown that mice initially prefer corners and the periphery to holes in the centre. This makes learning the baited holes in the centre more difficult .

### Habituation and training in the holeboard

During pre-training and holeboard training, pigs received one-third of their daily food allowance before behavioural testing in the early morning and two-thirds after testing in the late afternoon. All pigs were habituated to the holeboard before the acquisition phase started. During the habituation phase, each of the 16 holes contained a food reward (Fig. 1C), encouraging the pigs to search the entire holeboard arena. During the acquisition phase, four of the holes were baited with food (Fig. 1D).

The exact habituation methods are described in detail in the Materials and Methods sections of the respective original publications (S2-Supporting Information, Table S2.2), whereas an overview of post-weaning housing conditions and training and testing procedures for each HB study is given in S2-Supporting Information, Table S2.2. After thorough familiarization with the experimenter(s), test site, holeboard apparatus, and reward, formal training began, in which four of the 16 holes were baited with a food reward. On each trial, a pig was released into the holeboard and voluntarily walked around the perimeter of the arena until it found the open entrance to the arena, the location of which was randomly selected on a trial-by-trial basis. Pigs could then search the arena for the baited holes.

Trials lasted until all four rewards were found or until a preset maximum trial duration had elapsed, whichever event occurred first. For each trial, WM and RM were calculated as ratio measures, which reduces the bias due to incomplete trials (van der Staay *et al*. 2012).

#### WM was operationalized as

[Number of rewarded visits divided by the number of visits and revisits to the baited set of holes]. A score of 1 implies that a pig had not revisited any of the holes of the baited set in a trial.

#### RM was operationalized as

[Number of visits and revisits to the baited set of holes divided by the total number of visits to all holes]. A score of 1 implies that a pig had only made visits to baited holes in a trial.

In addition, the inter-visit interval (IVI) was determined as a proxy for the pigs’ motivation to search for food rewards.

#### IVI was operationalized as

[Time (s) between first and last hole visit divided by (number of hole visits-1)].

**Figure 1.**
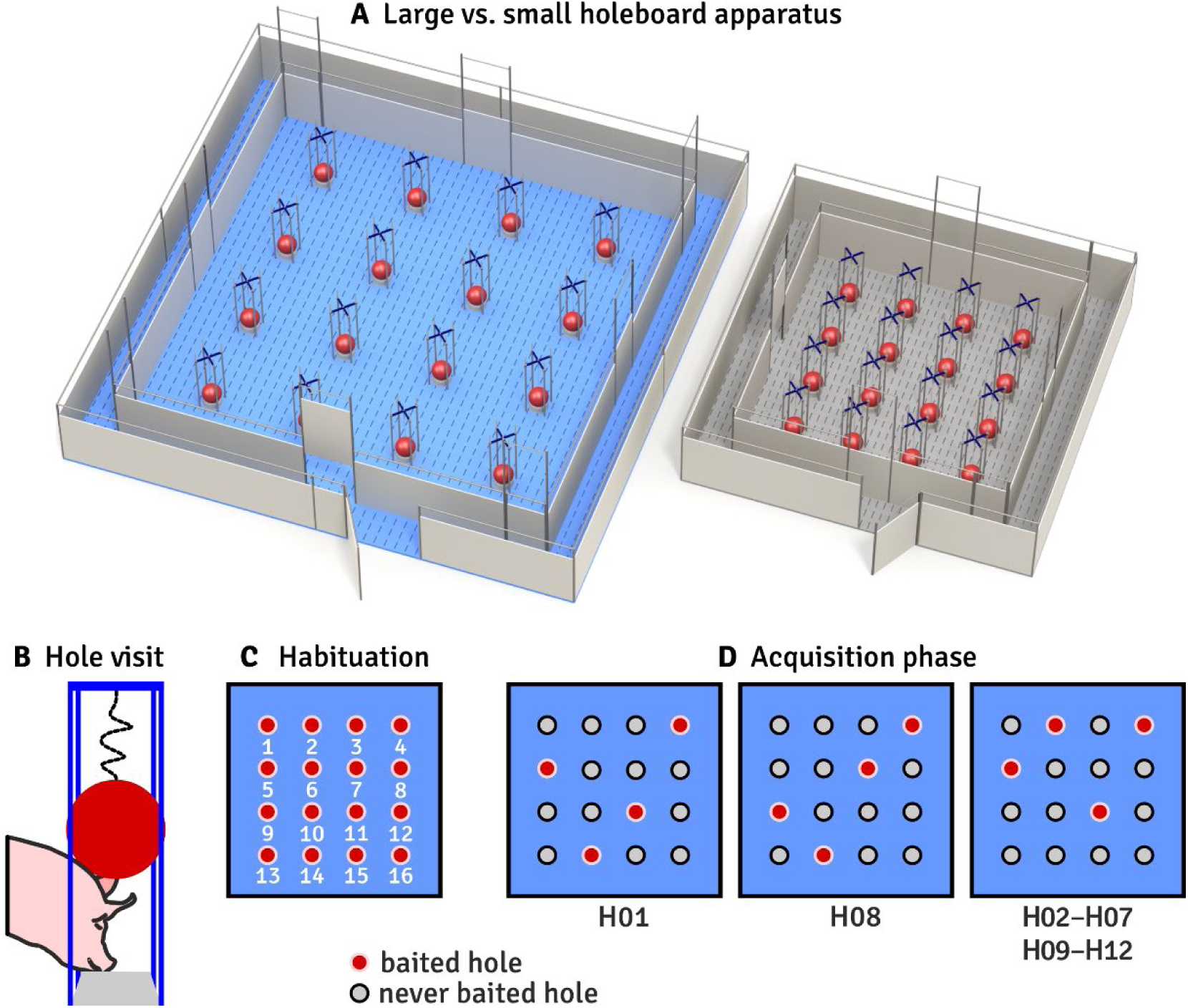
A side-by-side comparison of the large (panel A, left) and small holeboard apparatus (panel A, right), a pig pushing up the ball covering the food trough (panel B), configurations of baited holes during the habituation (panel C) and acquisition phases (panel D). During habituation, all 16 holes were baited with a food reward. Three different configurations of baited holes were used during the acquisition phase of the studies analysed, where H01 to H12 refer to the publications listed in S2-Supporting Information, Table S2.1. The configuration was also used rotated 90, 180, and 270 degrees in all studies. Panel B shows a pig discovering food in a trough hidden under a ball. The pig manages to push up the ball and retrieve the bait from the food trough. Hole visits were registered by a detector positioned in the centre under the feed trough (panel 1A drawn by Yorrit van der Staay).

### Statistical analyses

After about 40 trials (10 trialblocks), the pigs typically approached ceiling performance on the holeboard task, i.e., performance in later trials (overtraining trials) did not reveal any additional information about task acquisition (see e.g. Fig. 1 in Gieling *et al*. 2013, which shows the learning curves of pigs overtrained to a total of 104 acquisition trials). In the first study (H01), the pigs received only 26 acquisition trials. Therefore, for H01, we used the means of the first 6 trialblocks-each representing the mean of 4 trials-in the statistical analyses (S2-Supporting Information, Table S2.2), while 10 trialblocks were used in all other studies.

Replicability was analysed using data from all individuals or from subgroups of the studies (see S2-Supporting Information, Table S2.3 for details). The selection was based on the results of statistical tests reported in the original publications.

Replicability was also analysed in a subset of 8 studies which all used (Terra X Finnish Landrace) X Duroc pigs (S2-Supporting Information, Table S2.1).

Normality of the data was checked in the original publications and showed that WM and RM could be analysed untransformed, whereas IVI had to be log_10_(x+1)-transformed to meet the normal distribution requirements. Within all studies except H04, we pooled the data of the different groups. Provided that the intra-subject variance exceeds the between-subject variance, pooling is not expected to bias results (Leger and Didrichsons 1994).

### Estimating intercept and slope of the WM and RM learning curves

A substantial proportion of the variation in the increase in WM and RM performance in the holeboard task was expected to be accounted for by the linear trend component (see Table 2B in Roelofs *et al*. 2017a). The learning process may be adequately described by a linear regression function of the form: y = b₀ + b₁x. In this function, ŷ is the predicted value of the response variable, b₀ is the y-intercept, b₁ is the regression coefficient (slope), and x is the value of the predictor variable.

To determine the percentage of variation explained by the linear, quadratic, and cubic trend components across trialblocks, a repeated measures analysis was performed for each study with successive trialblocks (6 trialblocks of 4 trials each for H01 and 10 trialblocks for H02-H12) as the within-subjects factor, supplemented by polynomial contrasts (SAS PROC GLM; SAS 14.3; SAS Institute Inc., Cary, NC, USA) for each of the 12 studies separately. These orthogonal trend components represent independent dimensions that help characterize differences in the shape of learning curves (Winer 1971, pp. 577-594).The aim of such an approach is to estimate the predicted value of a response variable Y (WM, RM) in terms of the value of the predictor variable X (WM or RM performance in successive trialblocks). We determined the percentage of variation explained by the linear, quadratic and cubic trend components of the learning curves (Cotton 1998; Winer 1971) in the holeboard task over successive trialblocks (Spowart-Manning and van der Staay 2005; van Luijtelaar *et al*. 1989). The percentage of variation in the learning curves explained by the linear, quadratic and cubic trend components was calculated as the percentage of the sum of squares of the linear component of the total within-subject sum of squares from the repeated measures analysis (Roelofs *et al*. 2017a).

SAS PROC REG was then used to estimate the intercepts and slopes of WM, RM, and IVI per individual pig. The slopes, which represent the linear change across trialblocks, were used in subsequent analyses. The faster the acquisition of the WM and RM components of the holeboard task, the steeper the slope.

### Comparing the replicability of learning curves between studies

We used performance in the first trialblock as a measure of initial performance level of WM and RM (alternatively, one also could have used the individual estimates of the intercept), and its slope as a measure of learning progress across trialblocks. These data were subjected to a one-way ANOVA with the factor Study (levels: H01-H12), supplemented by Sidak post hoc pairwise comparisons between studies (SAS PROC GLM). In addition, the same analysis was performed using only the studies with (Terra x Finnish Landrace) x Duroc pigs as subjects, i.e. studies homogeneous with respect to the pig breed.

In the analysis of differences in IVI between studies, unlike for WM and RM, a two-way ANOVA with the factor Study (levels: H04-H12) and the repeated measures factor Trialblocks (levels: trialblock 1-10) (SAS PROC GLM) was used to assess replicability. Data from nine studies were used, as one study (H01) had only six trialblocks in its acquisition phase and two studies (H02, H03) did not have IVI data available. In addition, the same analysis was performed using only studies that examined holeboard learning in the crossbreed (Terra x Finnish Landrace) x Duroc. Breed as a factor has been found to affect the results obtained by different studies (see Supplementary Information, Table S2.3: studies H02 and H07).

### Correlations between WM and RM

Performance in the first trialblock and the slope of the learning curves of WM and RM were taken as measures that together describe the learning curves of WM and RM. These measures were subjected to Pearson correlation analysis to investigate the independence of these spatial memory components (SAS PROC CORR).

### Replicability, assessed by determining the intraclass correlations as a percentage of the total random variation due to the experiments

The assessment approaches described above examine whether studies differ from one another, using analyses of variance and graphical comparison of WM and RM learning curves during the first 40 acquisition trials. The graphical comparison gives only a subjective impression of replicability. Unfortunately, no consensus has yet been reached on the exact definitions of reproducibility and reliability, or on the metrics used to assess them. However, Heyard et al. (2025) proposed different methods of assessing reproducibility and reliability between studies. As a quantitative complement, we estimated the proportion of variation in learning performance attributable to differences between studies.

Determining the replicability of learning curves presents its own challenges. To make learning curves comparable across studies, we reduced each pig’s learning curve to two variables: initial performance, represented by performance in the first trialblock, and the linear trend component (slope), which represents the speed of learning over successive trialblocks. These variables were derived for both WM and RM. The IVI was excluded from these analyses because the linear trend component varied greatly between studies and, in most cases, only provided a weak explanation of development across successive trialblocks (see Fig. 2).

We fitted a linear mixed-effects model to the data from all 12 studies and a subset of studies with (Terra x Finnish Landrace) x Duroc pigs as subjects for each of the four outcomes (analyses of WM and RM performance in the first trialblock and slopes of WM and RM). We analysed the data using the lmer function of the lme4 package (Bates *et al*. 2015) with studyID as a random factor for the first trialblock of WM and RM, as well as their slopes, in R (R Core Team 2023). We added studyID as a random effect to assess the correlation between individuals within a study (two-level model). For each model, we calculated the intraclass correlation coefficient (ICC) as the studyID variance divided by the total random variance (studyID variance + residual variance). This can be interpreted as the correlation between observations within the same study.

As calculated in the present analysis, an ICC = 0.3, for example, means that the degree of dependence (correlation) within a studyID is 0.3, or in other words, 70 percent of the random variation is caused by the differences between individuals within a study i.e. differences between study levels are relatively small.

## Results

The speed of learning of the WM and the RM, i.e. their increase across trialblocks, was predominantly covered by the linear trend components. This component explained between 45.2 and 95.4 percent of the increase of in WM performance and between 97.2 and 98.3 percent of the increase in RM performance (see Figure 2 and Table 1).

IVI learning curves showed the greatest variability across studies. As a result, the proportion of variation explained by the linear trend component varied widely—from 8.7% to 80.1%. In some studies, higher-order trend components accounted for a substantial portion of the variance (Table 1), indicating that, unlike WM and RM, changes in IVI over trials were not consistently linear.

**Figure 2.**
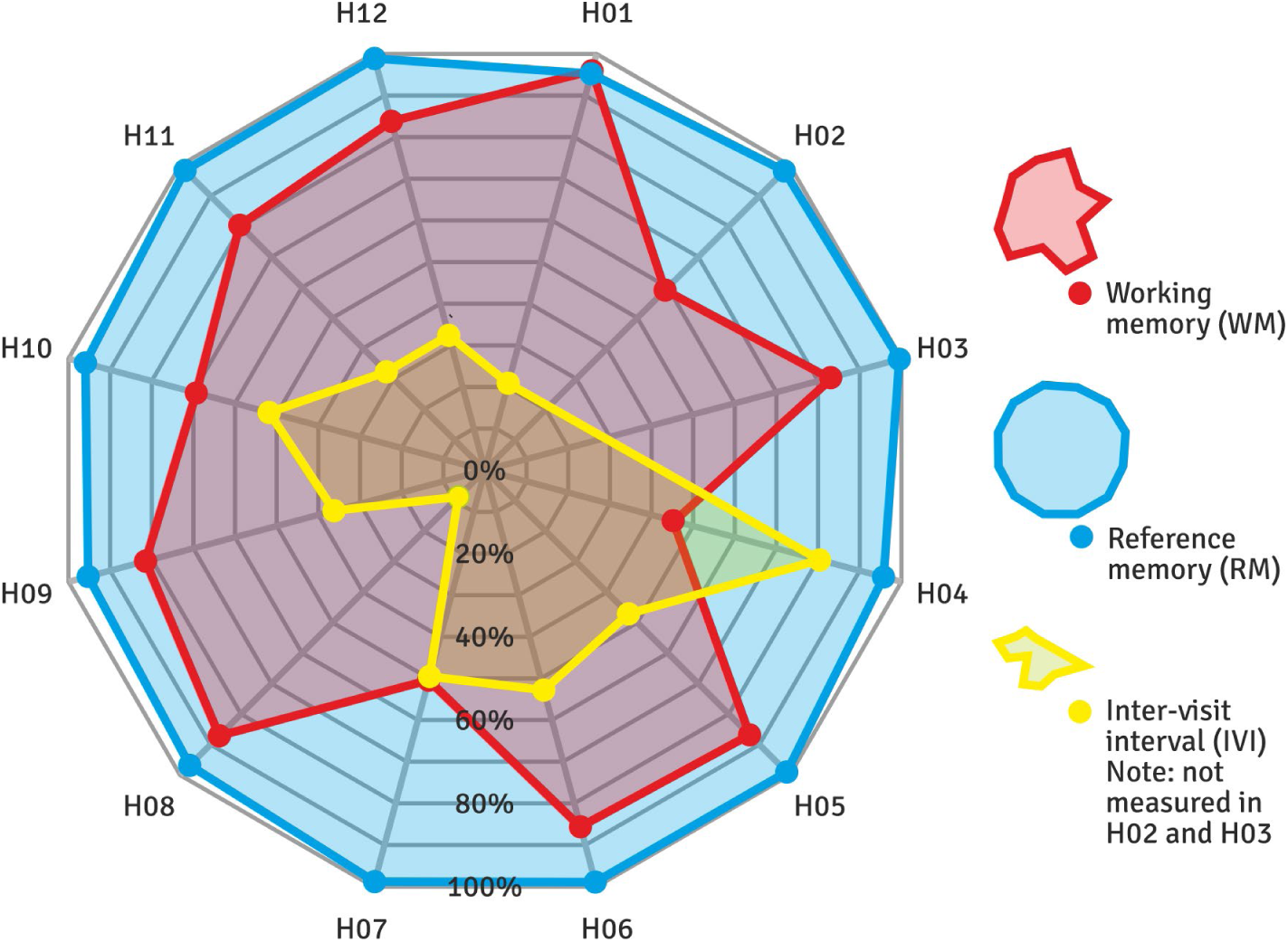
Spider plot showing the proportion of variation accounted for by the linear trend component in increases in working memory (WM) and reference memory (RM), as well as changes in the speed at which the holes were visited (i.e. the inter-visit interval, IVI), across all pig holeboard studies (H01–H12). Note that IVI was not registered in H02 and H03). The linear trend component accounted for a large proportion of progress in acquiring RM in all studies and WM in most studies, whereas the IVI showed much more erratic progression. See also Table 1, which lists the percentages of variation accounted for by the first to third order trend components.

**Table 1:** Percentage variation of the speed of learning for WM, RM, and IVI, accounted for by the linear (1^st^ order), quadratic (2^nd^ order) and cubic (3^rd^ order) components of the acquisition curves. The percentage variation accounted for by the sum of all trend components equal to or larger than the third-order trend is also given for the inter-visit interval. Abbreviations: comp., component; n.d., not determined

| Study | N | Percent variation explained by the linear, quadratic, and cubic trend component |  |  |  |  |  |  |  |  |  |
|---|---|---|---|---|---|---|---|---|---|---|---|
|  |  | WM |  |  | RM |  |  | Inter-visit interval |  |  |  |
|  |  | Linear (1<sup>s</sup> order) trend comp. | Quadratic (2<sup>nd</sup> order) trend comp. | Cubic (3<sup>rd</sup> order) trend comp. | Linear (1<sup>s</sup> order) trend comp. | Quadratic (2<sup>nd</sup> order) trend comp. | Cubic (3<sup>rd</sup> order) trend comp. | Linear (1<sup>s</sup> order) trend comp. | Quadratic (2<sup>nd</sup> order) trend comp. | Cubic (3<sup>rd</sup> order) trend comp. | Sum of trend comp. ≥ 4<sup>th</sup> order trend |
| **H01** | 18 | **95.4** | 1.6 | 1.1 | **94.7** | 3.6 | 1.0 | **20.9** | 52.4 | 26.5 | 0.3 |
| **H02** | 16 | **59.1** | 3.7 | 1.2 | **98.3** | 0.4 | 0.9 | n.d. | n.d. | n.d. | n.d. |
| **H03** | 37 | **83.0** | 4.5 | 0.2 | **99.0** | 0.1 | 0.1 | n.d. | n.d. | n.d. | n.d. |
| **H04** | 10 | **45.2** | 16.7 | 0.0 | **97.4** | 0.1 | 0.2 | **80.1** | 9.4 | 2.3 | 8.3 |
| **H05** | 19 | **86.3** | 1.5 | 6.8 | **98.0** | 0.8 | 0.2 | **48.0** | 37.9 | 1.9 | 12.2 |
| **H06** | 22 | **83.1** | 1.4 | 7.4 | **97.8** | 0.5 | 0.1 | **52.9** | 7.4 | 0.5 | 39.2 |
| **H07** | 20 | **51.0** | 11.1 | 0.2 | **97.6** | 1.8 | 0.5 | **49.6** | 30.3 | 5.0 | 15.0 |
| **H08** | 20 | **87.1** | 6.2 | 0.4 | **97.2** | 0.3 | 1.4 | **8.7** | 59.6 | 10.4 | 21.2 |
| **H09** | 19 | **80.2** | 1.9 | 4.0 | **95.7** | 0.0 | 0.4 | **36.2** | 0.4 | 44.7 | 18.6 |
| **H10** | 20 | **69.3** | 1.0 | 15.2 | **95.4** | 2.1 | 1.2 | **51.8** | 11.4 | 16.9 | 19.9 |
| **H11** | 38 | **80.8** | 2.3 | 1 | **98.7** | 0.2 | 0.8 | **32.3** | 38.6 | 24.6 | 4.4 |
| **H12** | 22 | **83.6** | 0.0 | 1.2 | **97.8** | 0.0 | 0.2 | **32.4** | 0.1 | 9.4 | 58.0 |

**Table 2:** The means and the lower and upper bounds of the 95% confidence intervals of the initial performance level (first trialblock) and the estimate of the slopes (linear change) across trialblocks of WM and RM are tabulated.

| **Study** | **N** | **Initial performance level (mean of first trialblock; lower and upper bonds of 95% confidence interval)** |  | **Speed of learning the HB task (mean slope; lower and upper bonds of 95% confidence interval)** |  |
|---|---|---|---|---|---|
|  |  | **WM** | **RM** | WM | RM |
| **H01** | 18 | **0.73** (0.67–0.80) | **0.31** (0.29–0.33) | **0.042** (0.031–0.054) | **0.052** (0.035–0.068) |
| **H02** | 16 | **0.78** (0.75–0.82) | **0.31** (0.29–0.33) | **0.009** (0.001–0.019) | **0.044** (0.035–0.053) |
| **H03** | 37 | **0.75** (0.71–0.79) | **0.28** (0.26–0.30) | **0.018** (0.013–0.022) | **0.049** (0.044–0.055) |
| **H04** | 10 | **0.76** (0.63–0.88) | **0.30** (0.27–0.33) | **0.010** (-0.003–0.022) | **0.040** (0.028–0.053) |
| **H05** | 19 | **0.75** (0.69–0.81) | **0.29** (0.27–0.31) | **0.017** (0.010–0.023) | **0.029** (0.023–0.035) |
| **H06** | 22 | **0.80** (0.74–0.86) | **0.29** (0.27–0.31) | **0.017** (0.012–0.022) | **0.050** (0.044–0.056) |
| **H07** | 20 | **0.86** (0.81–0.91) | **0.29** (0.27–0.31) | **0.010** (0.004–0.016) | **0.052** (0.045–0.059) |
| **H08** | 20 | **0.81** (0.74–0.88) | **0.30** (0.28–0.32) | **0.013** (0.007–0.019) | **0.056** (0.049–0.063) |
| **H09** | 19 | **0.78** (0.73–0.82) | **0.36** (0.34–0.39) | **0.018** (0.013–0.023) | **0.049** (0.044–0.053) |
| **H10** | 20 | **0.80** (0.77–0.84) | **0.29** (0.27–0.30) | **0.016** (0.011–0.020) | **0.051** (0.043–0.059) |
| **H11** | 38 | **0.79** (0.76–0.83) | **0.31** (0.30–0.32) | **0.018** (0.014–0.022) | **0.058** (0.053–0.063) |
| **H12** | 22 | **0.88** (0.84–0.92) | **0.31** (0.29–0.34) | **0.010** (0.006–0.014) | **0.065** (0.061–0.068) |

The means of WM and RM and the lower and upper limits of their 95% confidence intervals are tabulated per study for initial performance level (performance in the first trialblock) and acquisition speed (slope across 10 consecutive trialblocks; except for H01, where slopes were estimated based on 6 consecutive trialblocks) in Table 2.

### Replicability of learning curves across studies: repeated measures analysis

#### Working memory

##### Comparison of WM across all 12 studies (Fig. 3, and Fig. 4A,B)

###### Initial WM performance (trialblock 1)

The initial WM performance level differed across the 12 studies (F_11,249_=3.12, p<0.0006). Sidak post hoc pairwise comparisons revealed that this study effect was due to H12 having a higher initial performance level than H01, H03 and H05. The other studies did not differ in initial WM performance.

###### Speed of WM learning (slope)

The rate of WM learning differed between studies (F_11,249_=7.67, p<0.0001) and was higher in H01 than in all other studies, which did not differ from each other, as confirmed by the Sidak post hoc comparisons between studies.

##### Comparison of WM across studies in (Terra x Finnish Landrace) x Duroc pigs

###### Initial WM performance (trialblock 1)

Performance in the first trialblock differed between studies (F_7,160_=3.47, p<0.0017). Sidak post hoc comparisons showed that pigs in study H12 had a higher initial performance level than pigs in studies H01, H03 and H05. Initial WM performance in the other studies did not differ.

###### Speed of WM learning (slope)

The speed of learning differed between studies (F_7,160_=10.23, p<0.0001). Pigs in H01 showed a steeper WM learning curve than pigs in the other studies, which did not differ, as confirmed by Sidak post hoc comparisons.

#### Reference memory

##### Comparison of RM across all 12 studies (Fig. 3, and Fig. 4C, D)

###### Initial RM performance (trialblock 1)

The initial RM performance level differed between studies (F_11,249_=4.64, p<0.0001). Sidak post hoc comparisons revealed that this effect was due to a higher initial performance level in H09, which exceeded that of all other studies except H01.

###### Speed of RM learning (slope)

The studies differed in the speed of RM learning (F_11,249_=5.89 p<0.0001). This effect was mainly caused by a lower learning rate in H05 than in most other studies except H02 and H04, as confirmed by Sidak’s post hoc comparison.

##### Comparison of RM across studies in (Terra x Finnish Landrace) x Duroc pigs

###### Initial RM performance (trialblock 1)

In all studies, pigs started at a similar RM performance level (F_7,160_=1.46, p<0.1845).

###### Speed of RM learning (slope)

The speed of acquisition of RM differed between studies (F_7,160_=6.75, p<0.0001). Sidak post hoc comparisons showed that the pigs in H05 improved their RM performance more slowly than those in the other studies, which did not differ from each other.

A similar pattern of differences between studies was found when pooling all 12 studies and when restricting analyses to only the (Terra x Finnish Landrace) x Duroc pig studies.

**Figure 3.**
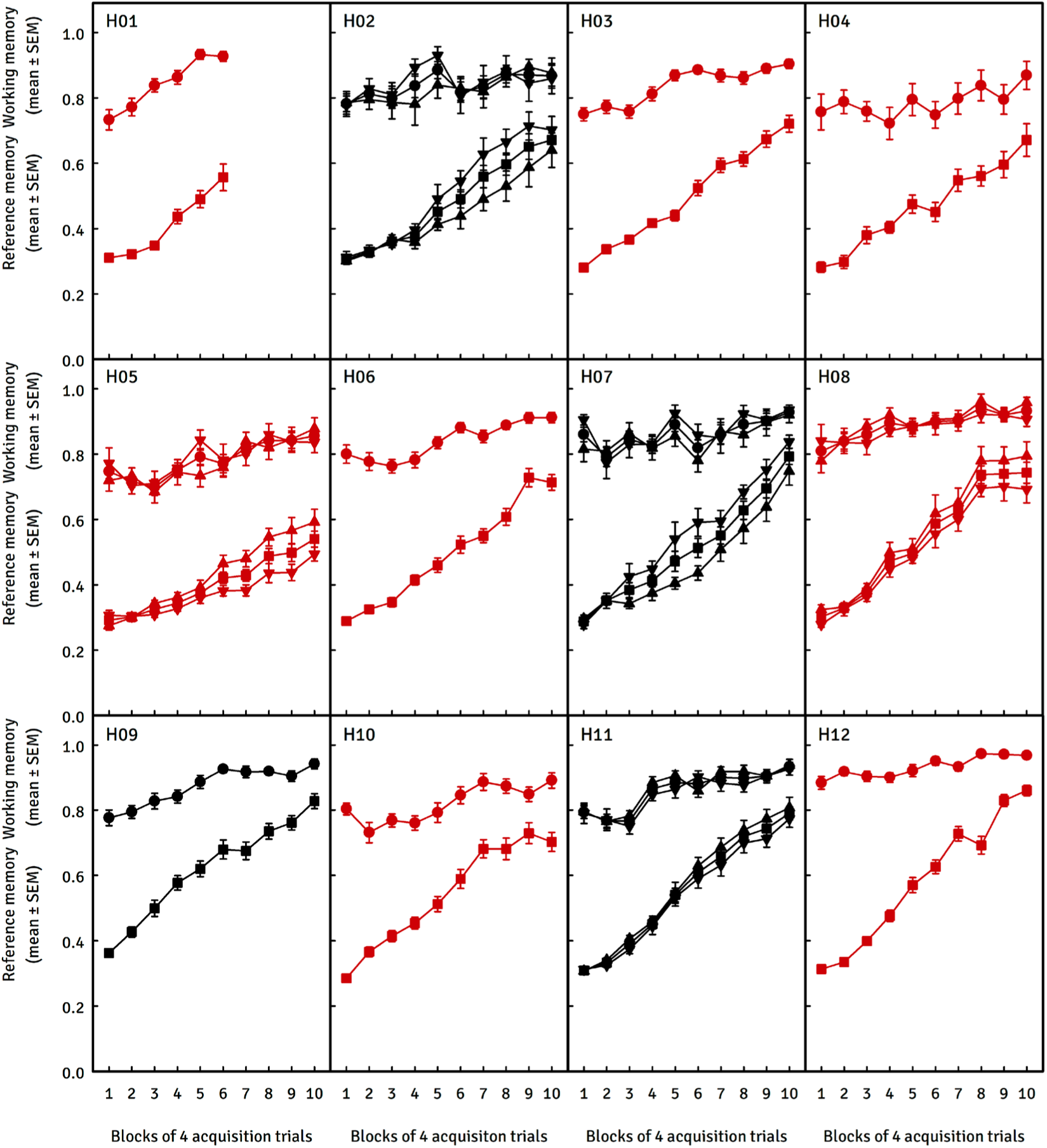
Acquisition of working and reference memory in pigs from the 12 original studies. Note that in studies H02, H05, H07, H08, and H11, data are presented per group because differences between groups were found in the original study (S2-Supporting Information, Table S2.3, and a brief description of these differences below). In these plots, the curve with the symbol λ represents the mean WM, while the curve with the symbol ν represents the mean RM performance of both groups. Shown in red are the eight studies where learning curves were obtained with (Terra x Finnish Landrace) x Duroc pigs. **H02:** Conventional pigs ▾ showed a faster RM acquisition than Göttingen minipigs ▴ **H05:** Slower RM acquisition in NBW pigs ▾ than in vLBW pigs ▴ **H07:** Large White x PIC426 pigs ▾ showed faster RM acquisition than T40 x Pietrain pigs ▴ **H08:** Faster RM acquisition in conventional pigs from an enriched environment ▴ than from a barren environment ▾ **H11:** Normal Birth Weight pigs ▴ showed a slightly faster RM acquisition than Low Birth Weight pigs ▾

**Figure 4.**
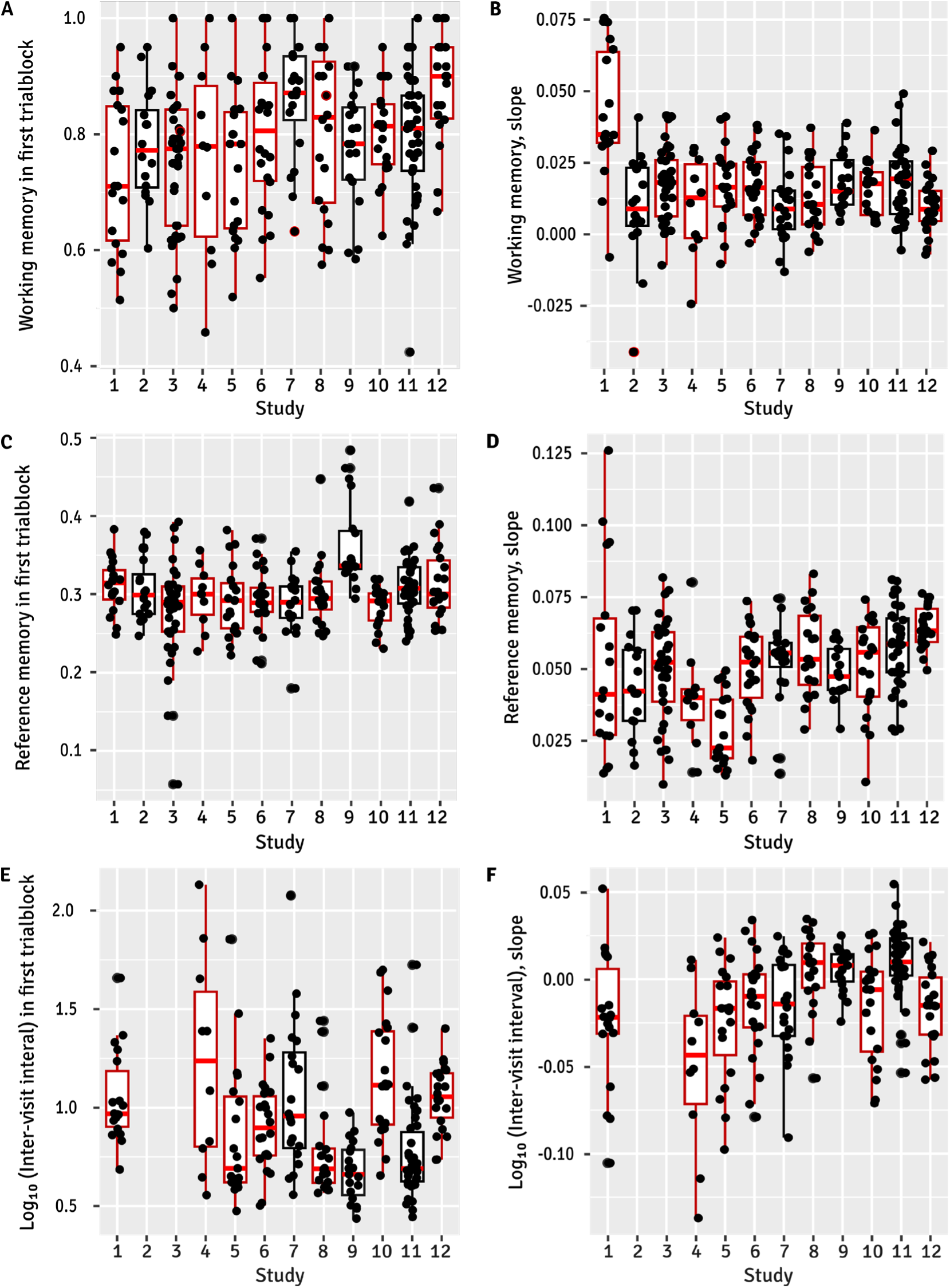
Performance in the first trialblock (panels A, C, E) and slopes (linear trend component; panels B, D, F) as an index of learning rate in pigs from 12 different holeboard studies (1-12). All box plots also include the individual value per pig λ. The eight studies in which learning curves were obtained with (Terra x Finnish Landrace) x Duroc pigs are shown as dark red boxes (Boxplots by ggplot2, Wickham 2016). The median is represented by a horizontal red lines; the box indicates the upper and lower quartiles; the lines extending from the box (whiskers) show the range of values within Q3 + 1.5 × Interquartile Range (IQR) to Q1-1.5 × IQR, where the IQR is the difference between Q1 and Q3; potential outliers are shown as dots above and below the ends of the whiskers.

##### Inter-visit interval (*Fig. 4E, F* and *Fig. 5*)

In some experiments, the slope alone was a poor indicator of changes in IVI across trialblocks, as higher order trend components accounted for a large proportion of these changes (Table 2). Therefore, only the results of the repeated measures analyses are reported here.

**Figure 5.**
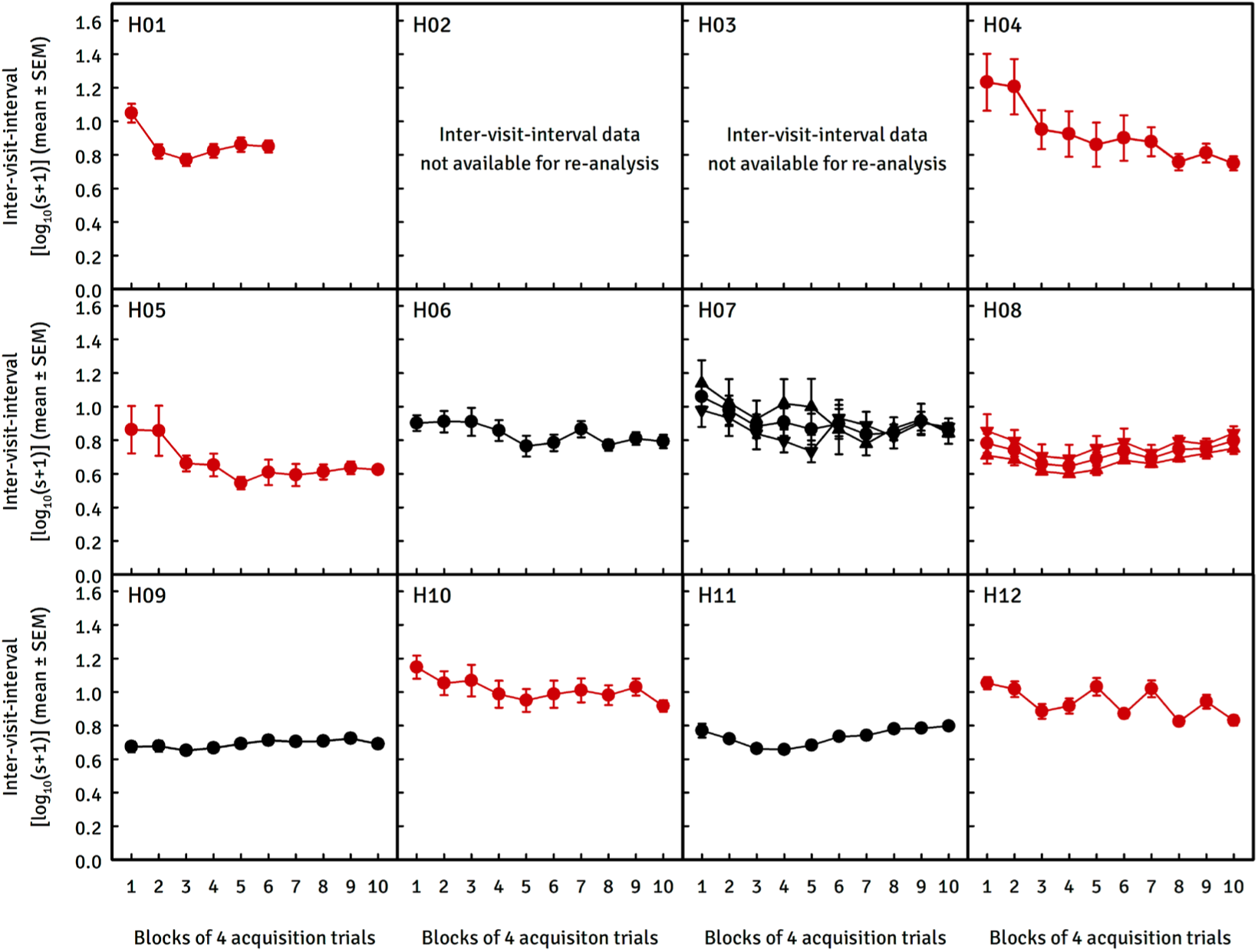
The inter-visit interval, i.e. the average speed of visiting the holes, in 10 of the 12 holeboard studies with pigs conducted in our laboratory. This measure may reflect the motivation of the pig to navigate the holeboard and find the food reward. Note that for studies H07 and H08, the curve with the symbol λ represents the mean IVI of both groups involved in the study. Shown in red are the studies where curves were obtained with (Terra x Finnish Landrace) x Duroc pigs. H07: ▴ T40 x Pietrain pigs, ▾Large White x PIC426 pigs H08: ▴ conventional pigs from a barren environment, ▾conventional pigs from an enriched environment

##### Comparison of the IVI across all 12 studies

Repeated measures analysis showed that the studies differed in mean inter-visit interval (F_8,181_=10.74, p<0.0001). The IVI fluctuated between trialblocks (F_9,1629_=20.66, p<0.0001) and these fluctuations differed between studies (Trialblock by Studies interaction, F_72,1629_=3.52, p<0.0001).

##### Comparison of the IVI across studies in (Terra x Finnish Landrace) x Duroc pigs

A similar pattern of differences between studies was found whether all 12 studies were compared or the analyses were restricted to the (Terra x Finnish Landrace) x Duroc pig studies. In this analysis, based on the data from H04, H05, H06, H08, H10 and H12 with a total of 113 (Terra x Finnish Landrace) x Duroc pigs, the mean IVI level across trialblocks differed between studies (F_5,107_=10.40, p<0.0001) and varied across trialblocks differently between studies (F_9,963_=17.73, p<0.0001) (trialblock by study interaction, F_45,963_=2.70, p<0.0001).

##### Correlation analysis

Across all 12 studies (Table 3, above diagonal), WM and RM performance in the first trialblock were uncorrelated. However, WM and RM slopes showed a very low correlation (r_PM_= 0.134, P≤0.030, N=261) when all 12 studies were considered. This correlation was not observed when the analysis was restricted to the eight studies using (Terra x Finnish Landrace) x Duroc pigs (r_PM_=0.099, P≤0.202, N=168; Table 3, below diagonal).

The initial performance of working memory (WM), i.e. performance in the first trialblock, was found to be correlated with the slope of the WM learning curves; the better the initial WM performance, the shallower the increase in WM learning curves across trialblocks.

**Table 3:** Pearson product-moment correlation coefficients (rPM) between WM and RM performance in the first trialblock (as index of the initial performance level) and slope (as an index of learning). Above diagonal: rPM across all 12 studies (N=261). Below diagonal: rPM across all studies with (Terra x Finnish Landrace) x Duroc pigs, (N=168).

|  |  | WM in 1<sup>st</sup> trialblock | WM slope | RM in 1<sup>st</sup> trialblock | RM slope |
|---|---|---|---|---|---|
| **WM in 1<sup>st</sup> trialblock** (initial performance level) | *r<sub>PM</sub>* | 1 | **-0.583** | -0.002 | 0.098 |
|  | *P* |  | ***<0.001*** | *0.978* | *0.113* |
| **WM slope** (working memory slope; linear increase) | *r<sub>PM</sub>* | **-0.588** | 1 | 0.040 | **0.134** |
|  | *P* | ***<0.001*** |  | *0.520* | ***0.030*** |
| **RM in 1<sup>st</sup> trialblock** (initial performance level) | *r<sub>PM</sub>* | 0.059 | 0.080 | 1 | -0.035 |
|  | *P* | *0.445* | *0.305* |  | *0.573* |
| **RM slope** (reference memory slope; linear increase) | *r<sub>PM</sub>* | 0.107 | 0.099 | 0.041 | 1 |
|  | *P* | *0.167* | *0.202* | *0.600* |  |

### Replicability: the ICC as percentage of total random variation

The variance estimate for the latent variable and the proportion of variance explained by the studyID were used as indicators of similarity or dissimilarity between studies. The higher ICC, the less replicable the study is, as the indices characterising the learning curves of WM and RM are more strongly influenced by the study itself. A low ICC indicates high individual variation within a study, meaning that the type of individuals used (breed, sex, etc.) makes little difference.

If the ICC is high, the type of pig may be important for the reproducibility of the study. Table 4 shows the percentage of total random variation that can be attributed to the study effect, alongside the 95% confidence intervals (CI), for three sets of data composed using different selection criteria. However, the values in Table 4 for selection 2, which includes only the Terra x Finnish Landrace x Duroc pig breed, do not differ greatly from those of the other selections.

**Table 4.** The intra-correlation coefficient indicates the percent of total random variation that can be attributed to initial performance levels in the first trialblock, as well as the rate at which spatial working memory (WM) and reference memory (RM) improve in the holeboard test, as indicated by the linear increase, or slope. Abbreviation: CI, confidence interval

| **Selection \*** | **Pigs (N)<br>In analysis** | **Studies (N) in<br>analysis** |  | **WM 1<sup>st</sup><br>trialblock** | **WM slope** | **RM 1<sup>st</sup><br>trialblock** | **RM slope** |
|---|---|---|---|---|---|---|---|
| 1 | 261 | 12 | % | 9% | 27% | 15% | 20% |
|  |  |  | CI | 0.05–30 | 0.6–50 | 0.3–40 | 2.5–44 |
| 2 | 168 | 8 | % | 11% | 34% | 3% | 23% |
|  |  |  | CI | 0.05–30 | 0.05–70 | 0.007–20 | 1–50 |
| 3 | 177 | 12 | % | 7% | 33% | 16% | 35% |
|  |  |  | CI | 0.1–26 | 4–64 | 0.4–44 | 4–63 |

\* *The first selection used the broadest set of criteria to include as many pigs as possible from all twelve studies.*
*The second selection applied stricter criteria. Only the eight studies involving the Terra x Finnish Landrace x Duroc pig breed were included to increase genetic uniformity.*
*The third selection applied the strictest set of criteria to all twelve studies, incorporating all breeds tested in the holeboard. Even if appropriate statistical analyses revealed the treatment(s) to be ineffective, treatment groups within studies were excluded. For further results of analyses of the third selection of data, see 'S3 - Supporting Information - Alternative exclusion and inclusion criteria for selecting pigs per study for the replicability analyses'.*

The intraclass correlation coefficient (ICC) in the analysed studies is ≤ 0.35, i.e. the variation between individuals is quite large compared to the differences between group means. Consequently, holeboard studies with pigs as subjects appear to be reasonably to highly reproducible. The slope of the studies is primarily determined by individuals (with values greater than 0.65) rather than by the studies themselves. However, it should be noted that the 95% confidence intervals of these measures were quite large. The ICC is based on variances (the square of the standard deviation), which means that variation can quickly increase dramatically, resulting in a wide confidence interval. In this case, the 95% CI is based on the model results, after which simulations are performed to determine the CI.

In all three selections, the random factor study of the initial performance level of the pigs (i.e. the first trialblock) was very low, ranging from 3 to 16%. For the rate of learning, as indicated by the linear increase across trialblocks (slope), the random factor studyID accounted for a larger proportion, ranging from 20 to 35 per cent. It is noteworthy that the confidence intervals (CIs) for the linear increase across trialblocks for both WM and RM were relatively large.

The percentages of total random variation in Table 4 for Selection 2, which includes only the Terra x Finnish Landrace x Duroc pig breed, are very similar to those of the other two selections. This suggests that the pig breeds had no effect on the replicability analyses.

As previously mentioned, the above analyses did not consider the inter-visit interval because the values of the first trialblock and the linear trend component (slope) of this measure are likely to be directly affected by environmental factors, such as ambient temperature. Such conditions would primarily impact the speed at which the pigs navigated the maze rather than their cognitive performance, as represented by WM and RM.

## Discussion

### Replicability of the holeboard task

In general, when two studies are designed to address the same scientific question, the consistency of their results reflects the degree of replicability. However, there are factors that can limit replicability: in particular, measurement precision and methodological differences between studies (Committee on science, engineering, medicine, and public policy 2019). Unfortunately, there are no formal criteria for when a replicability study can be considered successful; that is, we lack a rigorous theory for thoroughly understanding real-world non-exact replications (see, e.g. Buzbas and Devezer 2023). Some authors have addressed the question of which statistical procedures might be useful for analysing replicability (e.g., Errington *et al*. 2021), but no consensus has yet been reached on the most appropriate approach. As far as we know, no initiatives have been launched to reach this consensus. There are also no standardised approaches to determining replicability (e.g., Heyard *et al*. 2025), which may differ depending on the purpose. We chose to assess replicability by determining the intraclass correlation as a percentage of the total random variation due to the study.

It should be noted that the 12 holeboard studies used in the present analyses were not pre-planned replication studies for which recommendations for study design and statistical analyses were proposed (Frommlet and Heinze 2020). Many aspects of our replications are imprecise: the populations from which the samples were drawn, unequal sample sizes, differences in housing, age at testing, habituation and testing procedures, etc. These and other as yet unidentified factors may have an unpredictable effect on the results of an individual study. The three measures considered, WM, RM and IVI, varied between studies, i.e. showed a “study effect”. However, consistent patterns across heterogeneous study designs reflect an additional degree of replicability.

We characterized the acquisition of the WM and RM components of spatial memory in the HB task using two measures: initial performance level (mean of the first trialblock) and learning rate (slope of the learning curves for WM and RM across successive trialblocks of the acquisition phase).

#### Working memory

Firstly, we found that the initial working memory (WM) performance level was similar in most studies, but higher in study H12. Secondly, the learning rate was also similar in most studies, except in study H01, where the increase in WM performance was steeper. In retrospect, the detection system in the holes may not have worked optimally in study H01. The system may have failed to detect visits, and this may have (slightly) overestimated the performance of the pigs. The system underwent improvements in the course of the reported series of studies, and its reliability to detect hole visits was confirmed by Melendez et al (2013). The major improvement was to hang the balls on a stainless-steel wire, which ensured that when the pig retracted its head and the ball dropped back onto the food trough, the magnet in the ball was positioned exactly over the sensor in the food trough (Figure 1B).

The WM performance was reproducibly high from the start of the acquisition phase. Poor initial working memory could be an indication of adverse factors that have a negative effect on performance and have gone unnoticed. In all studies, a slow but steady improvement in WM performance was observed during the acquisition phase. However, the scope for further improvement through additional holeboard training is very limited. This may be explained by a natural, innate, win-shift foraging strategy (Gustafsson *et al*. 1999), which is an ethologically relevant characteristic of pig foraging behaviour (Allen *et al*. 2023). After finding and consuming the bait, the corresponding hole will not be replenished in the same trial. Research has shown that pigs acquire and perform win-shift/lose-shift tasks more accurately and faster than win-stay tasks (Laughlin and Mendl 2000). This may explain the rapid attainment of a ceiling effect in the working memory learning curves for pigs. Allen et al (2023), using a T-maze alternation task designed to test spatial working memory in pigs, found that pigs alternated with high accuracy from the start of training. Because the hole visits were self-paced, the time between hole visits varied. One might speculate that the time between visits might affect working memory performance. Allen et al. (2023) addressed this question in their T-maze working memory study with pigs and found no evidence for effects of the length of time between choices when they were self-paced.

Because of the high level of working memory performance in pigs from the start of training and the limited room for improvement to maximise performance, this component of spatial memory may be better suited to studying the effects of treatments that are expected to impair memory performance, such as ‘cognition impairers’, rather than the effects of treatments that are expected to improve memory performance, such as ‘cognition enhancers’ (for definitions of ‘cognition enhancer’ and ‘cognition impairer’ see van der Staay *et al*. 2011, p. 216). One study (Bolhuis *et al*. 2013) observed that the WM performance of pigs reared in an enriched environment exceeded the WM performance of pigs reared in a barren environment.

#### Reference memory

We found that the initial level of performance was very similar between studies (except for H09 where the pigs started at a higher level), i.e. they started at a level that would be expected for animals that did not know the location of the baited holes and thus found the rewards by chance. Progress in RM was almost entirely characterised as a linear increase across blocks of trials. The strong linear improvement in the performance of the RM over the course of training on the holeboard task has also been found previously in studies with rats (van der Staay *et al*. 1990a, 1990b) and can be considered a highly reproducible characteristic of this measure, at least for rats and pigs. The rate of learning of the RM was slowest in H05. In retrospect, it is not possible to identify the reasons for the small differences in RM between studies.

Inter-visit interval: The stalls and the test site were housed in a building whose internal temperature closely mirrored the outdoor weather conditions. The significant differences in ambient temperature between studies and the sometimes drastic temperature fluctuations during the studies very likely affected the pigs’ motivational state, as measured by the inter-visit interval.

Consequently, the development of IVI was quite variable between studies (see Fig. 4, panels E,F, and Fig. 5). This measure may be sensitive to the environmental conditions in which the pigs were tested and may reflect the motivation of the pig to seek the bait. For example, a high ambient temperature may be one of the factors that slows down the speed of navigating the holeboard (S2-Supporting Information, Table S2.4.). However, the fluctuations in motivation were not reflected in fluctuations in the effectiveness of working memory (WM) and reference memory (RM) learning. This observation was recently supported by a holeboard study in pigs by Galvagnon et al. (2026), which specifically addressed pig motivation using multiple measures. They concluded that the scores for reference and working memory in their study were unaffected by the level of motivation, i.e. there was no evidence of an association between motivation and competence.

#### Correlations between WM and RM

Few studies in rodents have addressed the question of whether measures of spatial WM and RM in the holeboard are independent, i.e. whether they are likely to reflect the action of different neural substrates (see section 6.5 in van der Staay *et al*. 2012). Note that the correlation found between WM and RM slopes (r_PM_= 0.134) can be classified as weak, poor or negligible depending on the rating system used (see Table 1 in Akoglu 2018). This correlation was not confirmed in analyses restricted to the eight studies using (Terra x Finnish Landrace) x Duroc pigs, where r_PM_ was as low as 0.099. Furthermore, the correlation was not supported in an analysis based on the selection of pigs using strict exclusion/inclusion criteria (see Table S3.2 in S3-Supporting Information) where rPM was 0.088. Thus, the results of all correlation analyses (Tables 3 and S3.2) suggest that RM and WM are independent of each other.

#### The ICC as percentage of total random variation due to study as index of replicability

We used the ICC, expressed as the percentage of total random variation attributable to differences between studies, as an index of replicability: the lower the ICC, the more similar the results across studies. However, it should be noted that this is a relative measure. If the total variation is large, there could be large differences between studies, as well as large individual differences. Even in case of little total random variation, the ICC can still be high if the differences between studies are relatively large compared to the smaller individual variation (residual variance).

Regardless of the selection criteria used to include or exclude groups of pigs, the WM and RM performance in the first trialblock was very similar between studies. This is because, in the first trialblock, RM performance is at chance level and WM performance is mainly driven by the pigs’ spontaneous win-shift search strategy rather than learning. The low between-study variation in the first block therefore does not indicate replicability in a meaningful sense; the ICC is informative only for the slopes, which reflect speed of learning. We chose this simple index because the dataset we had at our disposal did not meet the underlying requirements for single laboratory replicability recently described for large phenotyping projects in rodents (von Kortzfleisch *et al*. 2020). These approaches assume that experimental conditions were systematically varied across studies. It should be emphasised that differences between our 12 studies were not the result of systematic or controlled heterogenizations of their experimental conditions. For example, the choice of pig breeds was primarily driven by availability in the breeding unit at the time.

Our findings should be interpreted with caution, particularly given the large 95% confidence intervals (CIs). Increasing the number of studies and the number of individuals within the studies included in the replicability analysis, as well as including studies performed with the same breed/strain of animal under highly similar conditions, may improve replicability. It should be noted that environmental and phenotypic heterogeneity can be incorporated into the experimental setup and standardised (see, for example van der Staay *et al*. 2010), enabling analysis of replicability under less homogeneous conditions.

It should also be kept in mind that “a successful replication does not definitively confirm an original finding or its theoretical interpretation. Equally, a failure to replicate does not disconfirm a finding, but it does suggest that additional investigation is needed to establish its reliability” (Errington *et al*. 2021, p. 1).

### The holeboard for assessing cognition in the pigs

A wide range of species, from fish to humans, have been tested in the holeboard task (van der Staay *et al*. 2012). Pigs are becoming increasingly relevant as a model species for biomedical research (Ayuso *et al*. 2021; Li *et al*. 2022; Lunney *et al*. 2021; Netzley and Pelled 2023) as they can bridge the gap between rodent studies and human studies (Gün and Kues 2014; Lind *et al*. 2007; Lunney *et al*. 2021; van der Staay *et al*. 2017). However, reference material on learning and memory in pigs remains relatively scarce (Gieling *et al*. 2011). Increasing the number of validated behavioural and cognitive tests could make pigs a more useful large animal model in neuroscience research.

Cognitive holeboards are a valuable family of tasks for measuring spatial memory performance in a variety of species, including pigs (van der Staay *et al*. 2012). In the current version of the holeboard, pigs access the hidden food reward by pushing up a ball with their snouts. Uprooting is a component of a pig’s natural rooting behaviour, which increases with greater food motivation (Beattie and O’Connellt 2002). This behaviour could be unambiguously classified as a “hole visit,” both through direct observation and via an automatic detection system.

In addition to the three variables analysed here to assess spatial learning — WM, RM and IVI — a variety of other parameters can be derived from this task. These include trial-based measures and measures calculated across trials, such as indices of the food search strategy employed (van der Staay *et al*. 2012). Information on spatial memory, foraging strategy, motivation and anxiety can be collected in parallel in the holeboard task and can contribute to a detailed analysis of factors that may influence memory performance. The holeboard data can be subjected to very fine-grained behavioural analyses (e.g. in addition to registering hole visits, behavioural analyses of the hole visits themselves can be performed). For example, analogous to Casarrubea and colleagues (2009), a detailed ethogram of the pigs’ search behaviour in the holeboard could be made to analyse their anxiety-related and exploratory behaviour, both of which can interfere with learning processes.

### Putative publication bias

Unfortunately, not all scientific knowledge is published with the same probability. This means that some information is more likely to be shared than other information, due to a phenomenon known as the ‘file drawer problem’ or ‘publication bias’. This phenomenon hinders scientific progress and wastes resources. Publication bias has been identified as one of the causes of poor replicability (Nosek *et al*. 2022).

Studies reporting ‘non-significant or unexciting’ results are particularly likely to remain unpublished. The same may apply to studies using animal models other than rodents. In our experience, editors and reviewers may reject a paper on the grounds that a ‘non-standard’ or ‘exotic’ animal model species has been used. Scientists who predominantly use rodent model species may consider studies using non-rodent species, such as large animals, to be less relevant because they believe that results obtained with pigs, for example, cannot be compared with or generalised to those obtained in rodent studies (Curry *et al*. 2025). This view may result in authors deciding not to report the results of their study, or if they do submit them, they may face harsher peer reviews and a higher likelihood of lower citation rates (Curry *et al*. 2025; Thornton and Lee 2000).

Replicability analyses, much like systematic reviews, can be hindered by publication bias. This type of analysis should therefore consider whether the studies are sufficiently similar in design and conduct, whether they are free from serious risk of bias, and whether any observed statistical heterogeneity has been taken into account and addressed (Lensen 2023). We have restricted our analyses to in-house performed experiments. All HB experiments ever conducted on pigs by our research group, regardless of whether they found effects of the experimental manipulations or the division into groups with different characteristics, have been published and are included in this study. All relevant information and data were therefore available and used to analyse the replicability of acquiring the HB task. As the above criteria have been met, we can conclude that the present replicability study is free of publication bias (Berinsky *et al*. 2021; Curtis and Abernethy 2015).

### What has been replicated?

Goodman et al. (Goodman *et al*. 2016) distinguished three aspects of replicability or reproducibility: methods reproducibility, results reproducibility and inferential reproducibility.

“Methods reproducibility refers to the provision of enough detail about study procedures and data so the same procedures could, in theory or in actuality, be exactly repeated” (Goodman *et al*. 2016, p. 1). All of our HB studies using pigs as subjects have been published with their methodological details. Also, the equipment used was the same across studies, except for the smaller dimension of the holeboard in studies H04 and H05. We optimized the equipment used in the course of the period in which HB studies were performed. This includes an improved detection method (i.e. hanging the ball on a guide wire ensured that the magnet in the ball always fell back precisely above the sensor in the food trough; see Fig. 1B) and improved software for detecting and registering hole visits. These improvements may have helped to increase the reliability of data obtained, but simultaneously may have contributed to differences between studies.

“Results reproducibility (previously described as replicability) refers to obtaining the same results from the conduct of an independent study whose procedures are as closely matched to the original experiment as possible” (Goodman *et al*. 2016, p. 2-3). In all studies, all pigs were able to learn the HB task, irrespective of differences between studies, such as the time of year, testing environment, breed, sex, differences between the holeboard apparatuses used, etc. The suitability to assess pig learning and memory has been confirmed in other laboratories (Arts *et al*. 2009; Bolhuis *et al*. 2013; Clouard *et al*. 2016, 2021; Gautier *et al*. 2018; Haagensen *et al*. 2013a, 2013b; Val-Laillet *et al*. 2017).

Inferential reproducibility “refers to the drawing of qualitatively similar conclusions from either an independent replication of a study or a reanalysis of the original study. Inferential reproducibility is not identical to results reproducibility or to methods reproducibility, because scientists might draw the same conclusions from different sets of studies and data or could draw different conclusions from the same original data, sometimes even if they agree on the analytical results” (Goodman *et al*. 2016, p. 4).This type of reproducibility may be of special relevance. As Dixon and Glover stated, “From the perspective of the field and for the advancement of scientific knowledge, it is unimportant whether a replication produces precisely the same result as the original study. Rather, what matters is whether the replication evidence supports the same interpretation as the original, namely that evidence exists for a theoretically interesting effect.” (2020, p. 2). From this perspective, our studies addressing related questions, namely the effects of birth weight on HB learning and memory, have yielded mixed results. We might conclude that interferential reproducibility has not yet been established for this research question. Additional studies with larger numbers of animals (i.e., greater statistical power) may be needed to resolve questions about the effects of birth weight on spatial learning. It should be noted that a single successful replication is not sufficient to conclude that the original study produced replicable results (Berndtson 1991; Hedges and Schauer 2019).

Additionally, it should be noted that holeboard studies with rats (van der Staay *et al*. 1990a, 1990b) also found that the slope (linear trend component) accounts for a substantial percentage of the variation in the learning curves. Thus, the pig studies could be considered quasi-replications of the earlier rat holeboard studies (see S1 – Supporting information-Classification of replication studies). By demonstrating intra-lab replicability of the holeboard task in pigs, we provide cross-species evidence that the learning curves of this paradigm are not species-restricted. This strengthens confidence in the task itself and lends additional credibility to earlier findings from rat studies.

#### Limitations of performing replication studies in large animal model species

The number of published studies on large farm animals remains low. Only a handful of laboratories have conducted pig holeboard studies, and none have conducted as many as we did (Arts *et al*. 2009; Bolhuis *et al*. 2013; Clouard *et al*. 2016, 2021; Gautier *et al*. 2018; Haagensen *et al*. 2013a, 2013b; Val-Laillet *et al*. 2017). Studies in large model species, such as pigs, require more resources and are often more logistically challenging and time-consuming than studies in rodents (van der Staay *et al*. 2017). Efforts to assess the replicability of long-term studies, which can take weeks or months to complete, are less common than efforts to assess the replicability of short-term studies, which can be finished within a few days or weeks (Locey 2020).

Studies in which the number of subjects that can be tested in one day is limited, and in which testing is spread over a long period, are usually conducted with the minimum number of subjects required for acceptable statistical power. In this context, estimating replicability across a larger number of studies helps determine the validity of a particular experimental approach and the suitability of the model species used. Our study comparing the performance of pigs in twelve different holeboard studies contributes to the expansion of the repertoire of cognitive tests that can be validly used with pigs.

#### Options for future research

Using pigs may open new research opportunities and pigs may gain a larger role in neuroscience research (Dwulit *et al*. 2025). Generalizability across species can be determined in quasi-replications (see e.g., S1-Supporting information, and van der Staay *et al*. 2010). The majority of neuroscience studies use rodents as model species. This is also true for studies using the holeboard. However, the holeboard testing procedures and equipment are similar for different species – the most obvious difference being the size of the holeboard arena, which is adapted to the size of the species tested. Other differences between studies in different species are related to the habituation procedures and the exact behaviour required to inspect holes and retrieve the bait. In rodent (e.g., van der Staay *et al*. 1990b) and pig studies conducted by other laboratories (Arts *et al*. 2009; Bolhuis *et al*. 2013; Clouard *et al*. 2016, 2021; Gautier *et al*. 2018; Haagensen *et al*. 2013a, 2013b; Val-Laillet *et al*. 2017), subjects are typically required to dip their heads into holes to access them. In all pig studies included in this replicability analysis, the subjects were required to push a ball covering a hole with their snouts to access the food reward. This behaviour resembles pigs’ natural rooting behaviour when searching for food (Studnitz *et al*. 2007). As the operationalisation of variables such as WM, RM and IVI and the testing procedures are identical, results of holeboard studies can be compared between species.

#### Benefits of determining the intra-laboratory replicability of baseline measurements

While most publications on the replicability of results from animal behaviour studies refer to the effects of experimental manipulations (Ellis 2022), assessing the replicability of the behaviour of untreated and control animals has not received much attention. However, determining the latter’s replicability may have its own advantages. One area where the replicability of untreated animals may be relevant is the behavioural phenotyping of mutants (Kafkafi *et al*. 2018; Voikar 2020) and the use of these mutants as animal models of human diseases, for example in pharmacological studies. The selection of the most appropriate model species and tests to address a particular research question can be improved by phenotyping species commonly used in neurobehavioral studies such as mice and rats, farm animals such as chickens, pigs, sheep, and pets such as rabbits, cats, and dogs. Reanalysis of datasets from experiments conducted under similar conditions in the same laboratory can be used to assess their replicability, but can also help to detect non-random changes over time, such as shifts in the performance of untreated control groups across studies (e.g. presumably due to genetic drift, van der Staay 1997).

– A high level of replicability of controls may allow the direction of changes due to treatments to be compared across experiments, e.g. whether a compound acts as a cognition enhancer or a cognition impairer, where the direction of the effect is less likely to be an artefact of a deviant control group.
– Reanalysis of a set of data from previously conducted (and published) studies may provide new information and insights without the need to conduct new studies. This approach fulfils a principle of the 3Rs (Russell and Burch 1959), namely reduction “(…) in the numbers of animals used to obtain information of a given amount and precision” (Tannenbaum and Bennett 2015, p. 128).

Research into the replicability of scientific results provides additional benefits and opportunities:

– Proven replicability helps to increase the level of confidence in the proficiency of testing within and between laboratories (Crofton *et al*. 2008);
– Replicability of strains/breeds or genetic mutants, control conditions and control treatments (e.g. treatment with a control agent in neurotoxicology research, Crofton *et al*. 2008) facilitates direct comparison between experiments;
– It is important that behavioural and physiological traits in an animal, whether due to experimental manipulation or spontaneous changes in the genome, are stably expressed across studies (Kafkafi *et al*. 2018);
– Demonstrated replicability is of paramount importance when a full dose-response curve cannot be determined in a single experiment but must be determined across studies. The behaviour of animals in the control condition of different studies should be similar in order to provide an adequate and stable baseline against which the effects of experimental manipulations and their dose dependency can be measured (e.g. Luyten and Beckers 2017);
– The robustness of results across controls in non-exact replications may indicate that the variables under consideration are robust (van der Staay *et al*. 2017) despite methodological differences (Zwaan *et al*. 2018). At the same time, they may be less sensitive to systematic experimental manipulation. It should be noted that the relationship between robustness and sensitivity has not yet been the subject of scientific research. Further studies are therefore needed to address this question;
– Although less likely, any treatment effects found (e.g. of a drug thought to impair or enhance cognition) could be caused by a randomly deviating control group. Knowledge of the robustness of control results could help to identify such anomalous results and should encourage further replication studies;
– Random variation between studies can be detected because untreated controls should behave similarly across studies, provided that housing and test conditions are sufficiently similar and potential and already identified confounding variables are well controlled (see, e.g., Jaric *et al*. 2023);
– Historical (control) data can help increase the statistical power of animal studies (see Bonapersona *et al*. 2021). However, our data show that the results of studies can vary, even when animals of the same breed are tested under highly similar conditions. Relying on control group data from a previous study, i.e. conducting a study without a control group, can easily lead to incorrect conclusions;
– Although they were unrecognized and consequently neglected at the time of a particular study, putative confounds such as seasonal effects or occurrence of genetic drift may eventually be identified retrospectively through comparisons with previous studies. An example of possible genetic drift has been described in (van der Staay 1997), albeit for a different species (rats) and a different spatial memory task (Morris water escape task). The exact role of these putative confounds ought to be systematically investigated in further hypothesis testing studies;
– Embracing replicability research and publishing its results encourages the development and use of more reproducible methods and workflows and supports the development of best standards and practices, i.e. this approach helps to increase scientific rigor and replicability (Wilson *et al*. 2023);
– Determining intra-laboratory replicability can be an important component of quality control measures. Most laboratories will have the necessary data to determine this type of replicability, even if they are examining the effects of different experimental treatments in successive studies. In other words, this is useful when studying the replicability of treatment effects and effect sizes is not an option.

## Conclusions and recommendations

Research groups that have amassed a substantial body of results from studies employing the same animal model and test procedures should make use of these data to assess replicability. Information on intra-laboratory replicability could be incorporated into a laboratory’s quality control measures by identifying factors that reduce replicability, with the aim of improving the materials and methods to increase replicability.

As an example, we compared the acquisition of a spatial holeboard task in pigs from 12 independent studies conducted in our laboratory. Three different sets of exclusion/inclusion criteria were used to select the groups of pigs per study that were used in the replicability analyses. The first selection included pigs that were not exposed to or affected by experimental manipulations. The second selection applied all the inclusion/exclusion criteria of the first, but only included eight studies on pigs of the Terra x Finnish Landrace x Duroc breed.

The results of an additional replicability analysis based on a third set of criteria are reported in ‘S3-Supporting Information’. The similarity of the results of the replicability analyses, which used three different sets of exclusions/inclusions, supports the assumption that the broadest set — namely, pigs that were not exposed, or that were exposed but unaffected, by the experimental manipulations — is valid for assessing replicability. This selection also has the advantage of providing the largest possible sample size for analysis.

We found that the learning curves of WM in most studies and the learning curves of RM in all 12 studies were predominantly characterised by a linear increase. However, the slope of the increase differed between studies. In other words, the characteristic shape — a linear increase — was replicable, but not the learning rate (slope). This was true for analyses based on each of the selection of groups of pigs using three different sets of exclusion/inclusion criteria.

The curves of IVI across trialblocks between studies did not have a characteristic shape but were heterogeneous. This likely reflected the animals’ acute motivational state to search for hidden bait, which could be greatly affected by environmental influences, such as ambient temperature (e.g., high temperatures reduce activity and foraging behaviour). However, these presumed motivational fluctuations did not affect WM and RM performance.

Similar original data from other research groups should be included in this type of replicability study, so that both within- and between-laboratory replicability can be determined. Clear criteria for determining replicability would be beneficial for discussions of the replicability of experimental results within and across studies, especially when considering complex processes such as learning progress over a series of learning sessions.

## Supporting information

S4 - supporting informatio

## Acknowledgments

We would like to thank the MSc and BA students who assisted with the original studies. We would also like to thank the biotechnicians and animal caretakers of the Educational Research Farm ‘De Tolakker’, as well as the clinics of the Department of Farm Animal Health of the Faculty of Veterinary Medicine at Utrecht University, for their dedicated and expert assistance in caring for the animals and helping to conduct the studies.

## Author contribution

Shortened form of author names: F. Josef van der Staay (FJvdS), Alexandra Antonides (AA), Alexandra K. Dwulit (AKD), Lisa Fijn (LF), Elise T. Gieling (ETG), Charlotte G.E. Grimberg-Henrici (CGEG-H), Ellen Meijer (EM), Sanne Roelofs (SR), Hans Vernooij (HV), Vivian Witjes (VW), Saskia S. Arndt (SSA). AA, CGEG-H, ETG, FJvdS, LF, SR, and VW planned, performed and analysed the original holeboard studies (listed in S2-Supporting Information, Table S2.1). FJvdS and HV conducted the replicability analysis. FJvdS authored the first version of this manuscript. AKD contributed significantly to writing latter versions. All authors (AA, AKD, CGEG-H, EM, ETG, FJvdS, LF, SSA, SR and VW) collaborated to finalize the manuscript and approved the final version.

## Data availability statement

See “S4-Supporting information-Data used for replicability analysis of 12 holeboard studies in pigs”

