## Supplementary material for "On the intra-laboratory replicability of results in animal cognition research, obtained in animals that were not exposed to, or unaffected by, experimental manipulations — Exemplified by spatial learning in pigs *(Sus Scrofa Domesticus)* across 12 holeboard studies": S1 - supporting information

### S1 – Supporting information - Classification of replication studies

Robust scientific findings are generally accepted to be reproducible, replicable, and generalizable. *Reproducibility* refers to a researcher's ability to duplicate the results of a prior study using the same materials and procedures as the original investigator. *Replicability*, on the other hand, refers to a researcher's ability to achieve the same results as a prior study using different data but the same procedures (Bollen *et al.* 2015). There is no consensus yet on how to classify replication study types, and different classifications and taxonomies coexist (e.g., Baldassarre *et al.* 2014; Bettis *et al.* 2016; Greulich and Brendel 2019; Hüffmeier *et al.* 2016; Matarese 2022; Schmidt 2009). We found a division into original study, exact replication and extended replication, the latter subdivided into systematic and differential replication, conceptual replication and quasi replication, useful (van der Staay *et al.* 2010, see also Fig. S1.1.).

Exact, close or direct replications are based on a narrow notion of replication, namely the repetition of an experimental procedure (Schmidt 2009) of the original study (read: experimental results published before, or existing knowledge; Greulich and Brendel 2019) as closely as possible, i.e. exact replications are essentially similar to the study to be replicated (van IJzendoorn 1994). According to Buzbas and Devezer (2023), basically only the random samples differ between replications of this type. All other replications must be considered as non-exact (Buzbas and Devezer 2023).

To determine the external validity or generalizability of the results, the replication may be extended to include additional factors that are either already identified or presumed to be relevant - i.e., by introducing them as independent variables in a multifactorial experimental design that is analysed using an appropriate multifactorial statistical model. *Extended replications* are based on a wider notion of replication, namely the repetition of a test of a hypothesis or a result of earlier work with different methods (Schmidt 2009; Zwaan *et al.* 2018). This approach has also been called systematic replication (Muma 1993) or differentiated replication (Lindsay and Ehrenberg 1993). In extended replication approaches, major aspects of the experimental conditions, such as rearing and housing environments, gender and age of the animal and the exact testing conditions that may have an impact on the generalizability of results of the original study are varied systematically. The number of questions addressed in an extended replication study is increased compared with the original study and its exact replication. Extended replications test hypotheses using altered approaches, i.e. they assess "the robustness of a theoretical claim to alternative research designs, operational definitions, and samples" (Zwaan *et al.* 2018, p. 4).

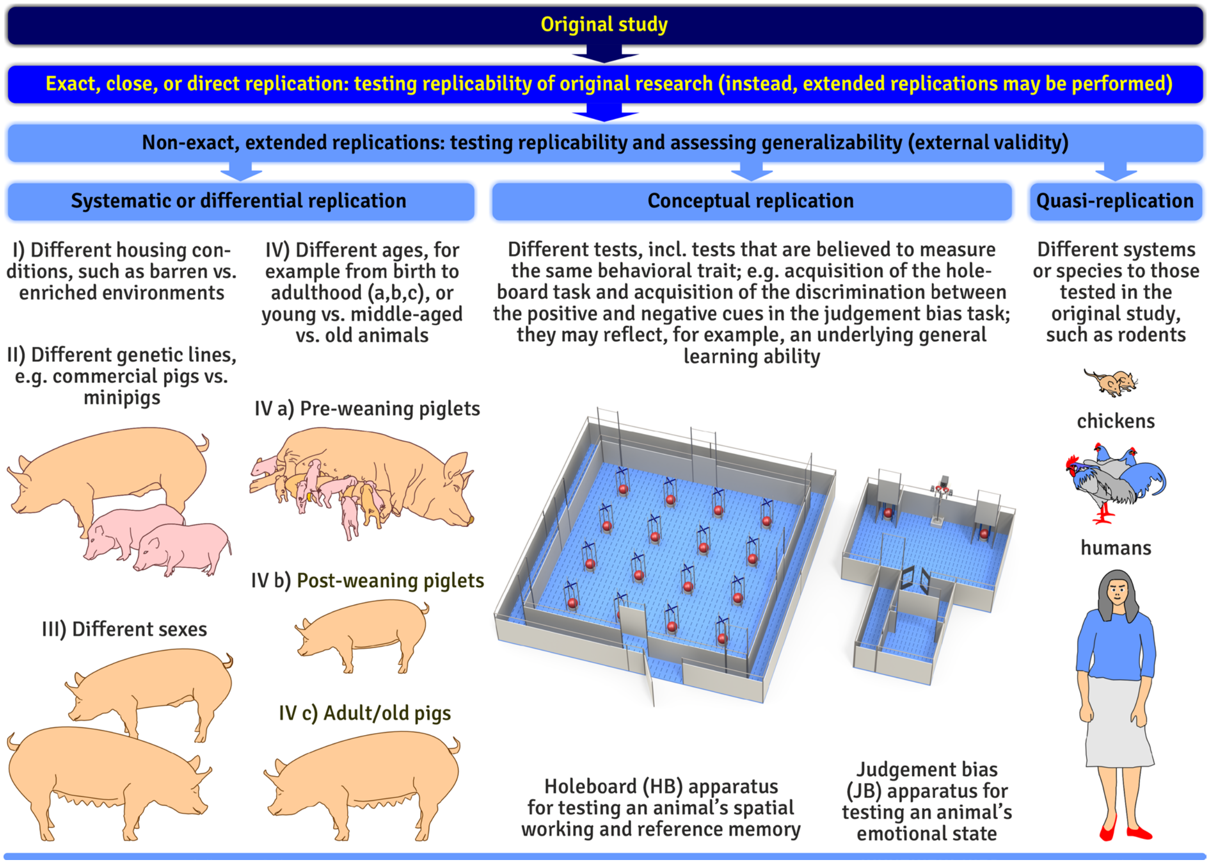


*Figure S1.1. Types of replication studies (modified from van der Staay et al. 2010)*

Other forms of extended replication are *partial replications*, in which (minor) procedural changes are introduced while all other aspects of the original study are closely replicated, and *'conceptual replications'*, in which the same relationships/constructs as in the original study are examined using different procedures.

Exact and extended replications differ only gradually from each other, as an exact replication will always deviate to some extent from the study it aims to replicate (Farrar *et al.* 2020; van IJzendoorn 1994). However, in exact replication approaches the differences from the original study are unintentional, whereas in extended replication approaches the differences are explicit modifications of the study design.

Finally, *quasireplications* test different species from those used in the original study (Palmer 2000). Quasireplications are a first step in assessing the effects of experimental manipulations in a different species, or in developing a new animal model using different species than in the original study. The latter can thus initiate a new process of model building and model evaluation; moreover, quasireplications may have an important role as the quickest route to generalizations (Bettis *et al.* 2016; Palmer 2000). Note that some (e.g., Forstmeier *et al.* 2017; Kelly 2006; Palmer 2000) advise caution in interpreting the results of quasireplications because of the lack of proximity to the study being replicated. Also, it may be a matter of speculation to explain a failure to replicate the original findings due to the larger differences between the original and the quasireplication study.

It should be noted that the cognitive holeboard task was originally developed and validated in rodents, specifically rats (Oades and Isaacson 1978; van der Staay *et al.* 1990, 2012). Since then, it has been adapted for testing other model animals species, including chickens (Nordquist *et al.* 2011), dogs (Smith *et al.* 2021), and pigs. According to the classification system adopted here, these studies—including those employed in our replicability analysis—can be considered quasi-replications of the original rat studies.
