## Supplementary material for "On the intra-laboratory replicability of results in animal cognition research, obtained in animals that were not exposed to, or unaffected by, experimental manipulations — Exemplified by spatial learning in pigs *(Sus Scrofa Domesticus)* across 12 holeboard studies": S2 - supporting information

#### S2 – Supporting information - Additional information about the twelve studies used to analyse the replicability of spatial learning in pigs in a holeboard task

### On the intra-laboratory replicability of results in animal cognition research, obtained in animals that were not exposed to, or unaffected by, experimental manipulations – Exemplified by spatial learning in pigs *(Sus Scrofa Domesticus)* across 12 holeboard studies

**F. Josef van der Staay, Alexandra Antonides, Alexandra K. Dwulit, Lisa Fijn, Elise T. Gieling, Charlotte G. E. Grimberg-Henrici, Ellen Meijer, Sanne Roelofs, J.C.M. (Hans) Vernooij, Vivian Witjes & Saskia S. Arndt**

We have addressed the question of the within-laboratory replicability of learning curves in the complex cognitive spatial holeboard discrimination task in pigs. Additional information about the included studies, the selection of variables used to assess the reproducibility of holeboard (HB) learning in pigs, and the selection of groups of pigs used for analyses is provided in Tables S2.1 – S2.4.

Table S2.1 provides an overview of the questions addressed by the hole-board studies included in the replicability analysis. It also lists the original publications and pig breed(s) tested in each study.

Table S2.2 summarises the post-weaning housing conditions and training and testing procedures in the HB studies.

Table S2.3 describes the groups of pigs tested in each study, and which of these groups were included in the replicability analysis.

Table S2.4 provides additional information about the inter-visit inerval (IVI) and about the ambient temperature during post-weaning housing and during testing (in °C).

##### Ethics statement

All of the studies whose results were used to evaluate the replicability of holeboard learning in pigs have been previously published in full in peer-reviewed scientific journals. They were conducted in strict accordance with the recommendations of the National Institutes of Health's Guide for the Care and Use of Laboratory Animals. Each of these studies was approved by the local ethics committee (DEC, DierExperimenten Commissie) or by the local Animal Welfare Body at Utrecht University, The Netherlands. The corresponding ethics statements can be found in the original publications (see references in Table S2.1.).

#### Selection of variables for assessing the replicability of holeboard (HB) learning in pigs

All 12 studies measured the working memory (WM) and reference memory (RM) components of the spatial holeboard (HB) task. With the exception of H02, each study also measured the inter-visit interval (IVI), which is used to assess the pigs' motivation to search the holes for the hidden food rewards (van der Staay *et al.* 2012).

#### Selection of the groups of pigs used for assessing the replicability of HB learning in pigs

In this study, the reproducibility of HB learning and memory in pigs was investigated in twelve studies conducted in our laboratory. An overview of the pig breeds used in these studies is given in Table S1.1. While seven different breeds were used in the 12 studies, eight studies used [(Terra x Finnish land-race) x Duroc] pigs. Table S1.2 provides an overview of the post-weaning housing conditions and a summary of the main characteristics of the training and testing methods used in the HB task in the 12 studies.

Finally, Table S1.3 provides an overview of the control and untreated groups of pigs included in the analyses per HB study. The included studies looked at differences between males and females, conventional pigs and minipigs, pigs from small and large litters, first- and last-born pigs in a litter or between different breeds. Some studies examined the acquisition of the HB task in untreated pigs that were subjected to various experimental manipulations after learning the task (using within-subject designs). The reference memory component was found to be sensitive to the effects of different experimental conditions, whereas working memory was hardly affected.

In six studies (H01, H03, H06, H09, H10, and H12) there were no differences between groups in how they learned the HB task. All data of pigs tested in these studies were used for further analysis. In the other six studies (H02, H04, H05, H07, H08 and H11) either treatment effects were found or the animals assigned to two groups according to birth weight or breed differed in HB acquisition. In H04, pigs that were deficient in iron acquired the task more slowly. The iron-deficient group was not included in the subsequent analyses. Breeds differed in two studies: Göttingen minipigs took longer to learn the task than conventional pigs in H02. In H07, T40 X Pietrain pigs took longer to learn the HB task compared to Large White x PIC426 pigs. We considered the breeds as valid samples of the population and used their average per study for further analyses, as we did for NBW and LBW pigs. In H08, pigs that were raised in an enriched environment learned the HB task faster than those raised in barren environment. We used the data from all pigs in this study for further analysis.

When comparing the studies that have examined the effects of birth weight on spatial discrimination performance, the results are mixed across studies. In H01 and H03, birth weight did not affect acquisition in the holeboard task. In H11, NBW acquired the task better than LBW pigs, whereas the reverse was found in H05. Therefore, we considered both NBW and LBW pigs as valid samples representing the study population and used the average of the birth weight groups per study for further analyses.

However, the most conservative selection of pigs for the replicability study excludes:

1) pigs with a low birth weight, as this would shift the distribution of birth weights towards lower values;

2) all pigs that have been subjected to pharmacological treatment or fed an iron-deficient diet;

3) Göttingen minipigs, as these differ from commonly used commercial pig breeds in some aspects (e.g. lower body mass, earlier onset of puberty and lower litter size (see, e.g., chapter 32 in Golledge and Richardson 2024)).

*Table S2.1. An overview of the studies used to determine the replicability of HB results in pigs and of the pig breeds trained in the 12 HB studies.*
*Abbreviations: H01–H12: holeboard studies. Note that data from studies H02, H03 and H10 have previously been used to assess the interdependence of judgment bias and cognition* (Roelofs *et al.* 2017)*; LBW, Low Birth-Weight; NBW. Normal Birth weight*

| Questions addressed in the original holeboard studies | Publication | Pig breed(s) tested in the holeboard * | |
|---|---|---|---|
| **H01** Low Birth Weight (LBW) vs. Normal Birth Weight (NBW) pigs | Gieling ET, Park SY, Nordquist RE, van der Staay FJ. Cognitive performance of low- and normal-birth-weight piglets in a spatial hole-board discrimination task. Pediatric research. 2012;71: 71–76. doi:10.1038/pr.2011.5 | (Terra x Finnish landrace) x Duroc | |
| **H02** Conventional pigs vs. Göttingen minipigs | Gieling ET, Wehkamp W, Willigenburg R, Nordquist RE, Ganderup N-C, van der Staay FJ. Performance of conventional pigs and Göttingen miniature pigs in a spatial holeboard task: effects of the putative muscarinic cognition impairer biperiden. Behavioral and brain functions. 2013;9: 4–4. doi:10.1186/1744-9081-9-4 | Duroc x (Yorkshire x Finnish landrace) | Göttingen miniature pig |
| **H03** Effects of prenatal allopurinol in LBW and NBW pigs | Gieling ET, Antonides S, Fink-Gremmels J, ter Haar K, Kuller WI, Meijer E, et al. Chronic allopurinol treatment during the last trimester of pregnancy in sows: effects on low and normal birth weight offspring. PLoS ONE. 2014;9: e86396. doi:10.1371/journal.pone.0086396 | (Terra x Finnish landrace) x Duroc | |
| **H04** Effects of an anaemic iron-deficient diet | Antonides A, Schoonderwoerd AC, Scholz G, Berg BM, Nordquist RE, van der Staay FJ. Pre-weaning dietary iron deficiency impairs spatial learning and memory in the cognitive holeboard task in piglets. Frontiers in behavioral neuroscience. 2015;9: 16. doi:10.3389/fnbeh.2015.00291 | (Terra x Finnish landrace) x Duroc | |
| **H05** very LBW vs. NBW pigs | Antonides A, Schoonderwoerd AC, Nordquist RE, van der Staay FJ. Very low birth weight piglets show improved cognitive performance in the spatial cognitive holeboard task. Frontiers in behavioral neuroscience. 2015;9: 10. doi:10.3389/fnbeh.2015.00043 | (Terra x Finnish landrace) x Duroc | |
| **H06** Effects of a non-anaemic iron deficient diet | Antonides A, van Laarhoven S, van der Staay FJ, Nordquist RE. Non-anemic iron deficiency from birth to weaning does not impair growth or memory in piglets. Frontiers in behavioral neuroscience. 2016;10:112. doi:10.3389/fnbeh.2016.00112 | (Terra x Finnish landrace) x Duroc | |
| **H07** Effects of litter size and supplier/breed | Fijn L, Antonides A, Aalderink D, Nordquist RE, van der Staay FJ. Does litter size affect emotionality, spatial learning and memory in piglets? Applied animal behaviour science. 2016;178: 23–31. doi:10.1016/j.applanim.2016.02.011 | T40 x Pietrain | Large White x PIC426 *(Note that PIC426 has been renamed PIC408M)* |
| **H08** Effects of environmental enrichment | Grimberg-Henrici CG, Vermaak P, Bolhuis JE, Nordquist RE, van der Staay FJ. Effects of environmental enrichment on cognitive performance of pigs in a spatial holeboard discrimination task. Animal cognition. 2016;19: 271–283. doi:10.1007/s10071-015-0932-7 | (Terra x Finnish landrace) x Duroc | |
| **H09** Effects of social isolation (after acquisition of the HB task) | van der Staay FJ, Schoonderwoerd AJ, Stadhouders B, Nordquist RE. Overnight social isolation in pigs decreases salivary cortisol but does not impair spatial learning and memory or performance in a decision-making task. Frontiers in veterinary science - animal behavior and welfare. 2016;2: 13. doi:10.3389/fvets.2015.00081 | Duroc x (Yorkshire x Finnish landrace) | |
| **H10** Males vs. females | Roelofs S, Nordquist RE, van der Staay FJ. Female and male pigs' performance in a spatial holeboard and judgment bias task. Applied animal behaviour science. 2017;191: 5–16. doi:10.1016/j.applanim.2017.01.016 | (Terra x Finnish landrace) x Duroc | |
| **H11** LBW vs. NBW pigs | Roelofs S, van Bommel I, Melis S, van der Staay FJ, Nordquist RE. Low birth weight impairs acquisition of spatial memory task in pigs. Frontiers in veterinary science - animal behavior and welfare. 2018;5:142: 14. doi:10.3389/fvets.2018.00142 | (Terra x Dutch landrace) x Duroc | |
| **H12** First vs. last born piglets in a litter | Witjes VL, Roelofs S, Meijer E, Eicher PHC, Zeinstra EC, Oei CHY, et al. Comparison of first- and last-born pigs revealed no effect of the birth process on acquisition and reversal of the cognitive holeboard task. Applied animal behaviour science. 2025;285, 106585: 10. doi:10.1016/j.applanim.2025.106585 | (Terra x Finnish landrace) x Duroc | |

**: Replicability was analysed using all 12 studies, and in a subset of studies that used (Terra x Finnish landrace) x Duroc pigs.*

*Table S2.2.* ***Post-weaning housing conditions and training and testing procedures in the HB studies.*** *Housing conditions include pen size, pigs per pen, presence of nest boxes and enrichment materials (the standard condition in each study) such as peat or straw bedding (FB), chains (C), chew sticks (CS), balls (B), ropes (R), gunny bags (G), number of feeders and drinkers per pen.*

*For the training and test phase, the test site (F: farm of the Faculty of Veterinary Science, University of Utrecht; C: clinics of the Faculty of Veterinary Science, University of Utrecht), the approximate age at habituation to the HB procedure and apparatus, the approximate age at the start of HB acquisition, the number of acquisition sessions per day and the number of trials per session, the maximum trial duration, and the minimum and maximum number of acquisition trials are listed. The last column indicates whether or not the HB acquisition data were reanalysed by calculating the means of blocks of 4 consecutive trials to achieve comparability across the 12 studies.*

*Note that the size of the holeboard differed between the two test sites (the holeboard arena in the clinics (C) was smaller than at the farm (F). The two holeboard are depicted side by side in Fig. S1.1 (below).*

| Holeboard study | Post-weaning housing conditions | | | | | | Habituation to the HB apparatus and acquisition of the spatial discrimination task | | | | | | | |
|---|---|---|---|---|---|---|---|---|---|---|---|---|---|---|
| | Approximate pen size (m) | Pigs per pen | Nest-box (Y/N) | Enrichment items | Feeders per pen | Drinkers per pen | Test site | Approx. age in weeks at start of habituation | Approx. age in weeks at start of acquisition | Training sessions per day, trials per session[1] | Maximal trial duration (sec.) | Minimum number of acquisition trials[2] | Maximum number of acquisition trials | Re-calculate trial-block means?[4] |
| **H01** – LBW vs. NBW pigs | 4 X 5 | 8, 10 | Y | FB | 1 | 2 | F | 4 | 9 | 2, 1 | 600 | 26 | 26 | yes |
| **H02** – Conventional pigs vs. minipigs | 4 X 5 | 8 | Y | FB, C, B | 1 | 2 | F | 10 | 13 | 2, 2 | 600 | 104 | 104 | no |
| **H03** – Effects of allopurinol in LBW and NBW pigs | 3 X 5 | 9 – 10 | Y | FB, CS, B | 0, on floor | 1 | F | 5 | 7 | 1, 2 | 600 | 40 | 60 | no |
| **H04** – Effects of an anaemic iron-deficient diet | 2.5 X 2.5 | 3 - 5 | N | FB, C, B, R | 1 | 1 | C | 5 | 7 | 2, 2 | 450 | 40 | 60 | no |
| **H05** – very low birth weight (vLBW) vs. NBW pigs | 2.5 X 2.5 | 4 - 5 | N | FB, B | 1 | 1 | C | 4 | 7 | 1, 4[3] | 450 | 46 | 54 | yes |
| **H06** – Effects of a non-anaemic iron deficient diet | 4 X 5 | 11 | Y | FB, CS, B | 1 | 1 | F | 6 | 10 | 1, 2 | 300 | 40 | 60 | no |
| **H07** – Effects of litter size and supplier/breed | 4X 5 | 5 | Y | FB, CS, B, G | 1 | 1 | F | 6 | 9 | 1,2 | 450 | 40 | 60 | no |
| **H08** – Effects of environmental enrichment | 4 X 5 | 10 | Y | FB, CS, B, G | 1 | 1 | F | 7 | 11 | 2, 1 | 300 | 40 | 40 | no |
| **H09** – Effects of social isolation | 4 X 5 | 9, 10 | Y | FB, CS, B, G | 1 | 1 | F | 8 | 10 | 1, 2 | 300 | 40 | 40 | no |
| **H10** – Males vs. females | 4 X 5 | 10 | Y | FB, CS, C, B | 1 | 1 | F | 7 | 10 | 1, 2 | 450 | 44 | 60 | no |
| **H11** – LBW vs. NBW pigs | 4 X 5 | 9, 10 | Y | FB | 1 | 1 | F | 5 | 8 | 1, 2 | 300 | 44 | 76 | no |
| **H12** – First vs. last born piglets in a litter | 4 X 5 | 11, 12 | Y | FB, C, CS, B | 1 | 2 | F | 3 - 4 | 11 | 2, 1 | 450 | 40 | 56 | no |

[1]: *One trial per acquisition session indicates that the pigs were trained with spaced trials, whereas >1 trial per acquisition session indicates that the pigs were trained with massed trials.*

[2]: *In most studies, the pigs were trained until they had reached a criterion of >0.7 on the reference memory measure on two consecutive trial blocks, but training was continued until they had had at least a minimum number of trials. In H01 and H02, the pigs got a fixed number of acquisition trials and no learning criterion was set.*

[3]: *In this study, the pigs were trained with 2 trials per session during the first 4 days of HB acquisition, followed by acquisition sessions with 4 trials each.*

[4]: *Note that in two studies (H01 and H05) block means of 4 successive trials each had to be calculated to enable comparability of the resulting learning curves with those from the other studies.*

*Table S2.3. **The groups of pigs that were included in the analyses per HB study.*** *All comparisons between studies are restricted to working memory (WM) and reference memory (RM) acquisition in the HB task. Note that we do not consider LBW vs. NBW pigs, differences between pig breeds, and males vs. females as representing different treatments. Abbreviations: N: total number of animals in a group, of these ♂: number of males, ♀: number of females; NBW : normal birth weight; LBW: low birth weight.*

| Holeboard study | Group 1 Short description | N | ♂ | ♀ | Group 2 Short description | N | ♂ | ♀ | Group 3 Short description | N | ♂ | ♀ | Group 4 Short description | N | ♂ | ♀ | Diff. between groups? | Which pigs are included in the present analysis? |
|---|---|---|---|---|---|---|---|---|---|---|---|---|---|---|---|---|---|---|
| **H01** Low Birth Weight (LBW) vs. Normal Birth Weight (NBW) pigs | Conventional pigs (NBW) | 9 | – | 9 | Conventional pigs (LBW) | 9 | – | 9 | | | | | | | | | no | All pigs (N=18); Birth weight had no effect on HB acquisition |
| **H02** Conventional pigs vs. Göttingen minipigs | Conventional pigs | 8 | – | 8 | Göttingen miniature pigs | 8 | – | 8 | | | | | | | | | no | All pigs (N=16); Minipigs (n=8) showed slower RM acquisition than conventional pigs (n=8) |
| **H03** Effects of prenatal allopurinol in LBW and NBW pigs | Conventional pigs (NBW, untreated) | 9 | 8 | 1 | Conventional pigs (LBW, untreated) | 8 | 7 | 1 | Conventional pigs (NBW, prenatal Allo treatment) | 10 | 5 | 5 | Conventional pigs (LBW, prenatal Allo treatment) | 10 | 6 | 4 | no | All pigs (N=37); Neither birth weight nor prenatal allopurinol affected HB acquisition |
| **H04** Effects of an anaemic iron-deficient diet | Conventional pigs, fed a balanced diet | 10 | 10 | – | Conventional pigs, fed an iron deficient diet | 8 | 8 | – | | | | | | | | | yes | Pigs fed a balanced diet (N=10); Iron deficiency impaired RM acquisition |
| **H05** very LBW vs. NBW pigs | Conventional pigs (NBW) | 10 | 6 | 4 | Conventional pigs (vLBW) | 9 | 5 | 4 | | | | | | | | | yes | All pigs (N=19); Faster RM acquisition in vLBW pigs (n=9) than in NBW pigs (n=10) |
| **H06** Effects of a non-anaemic iron deficient diet | Conventional pigs, fed a balanced diet | 11 | 6 | 5 | Conventional pigs, fed an iron deficient diet | 11 | 5 | 6 | | | | | | | | | no | All pigs (N=22); No effect of iron-deficient diet on HB acquisition |
| **H07** Effects of litter size and supplier/breed | Conventional pigs (T40 X Pietrain), litter size ≤ 13 | 5 | 5 | – | Conventional pigs (T40 X Pietrain), litter size ≥ 18 | 5 | 5 | – | Conventional pigs (Large White); litter size ≤ 13 | 5 | 5 | – | Conventional pigs (Large White); litter size ≥ 18 | 5 | 5 | – | yes | All pigs (N=20)*; Large White pigs (n=10) acquired RM faster than T40 pigs (n=10) |
| **H08** Effects of environmental enrichment | Conventional pigs, enriched environment | 10 | – | 10 | Conventional pigs, barren environment | 10 | – | 10 | | | | | | | | | yes | All pigs (N=20); Enriched pigs (N=10) acquired RM faster than barren pigs (N=10) |
| **H09** Effects of social isolation (after acquisition of the HB task) | Conventional pigs | 19 | – | 19 | | | | | | | | | | | | | not applicable | All pigs (N=19); No experimental manipulation before or during HB acquisition |
| **H10** Males vs. females | Conventional pigs (males) | 10 | 10 | – | Conventional pigs (females) | 10 | – | 10 | | | | | | | | | no | All pigs (N=20); No differences in HB acquisition between males and females |
| **H11** LBW vs. NBW pigs | Conventional pigs (NBW males) | 9 | 9 | – | Conventional pigs (NBW females) | 10 | 10 | – | Conventional pigs (LBW males) | 10 | – | 10 | Conventional pigs (LBW females) | 9 | – | 9 | yes | All pigs (N=38); The LBW pigs (n-19) acquired RM at a slower rate than the NBW pigs (n=19). |
| **H12** First vs. last born piglets in a litter | Conventional pigs, first born | 11 | 5 | 6 | Conventional pigs, last born | 11 | 7 | 4 | | | | | | | | | no | All pigs (N=22); No difference between first- and last-born pigs for HB acquisition |

**: Note that in H07, pigs from both suppliers were considered as valid controls. However, supplier [1: conventional breeding facility vs. 2: specific pathogen free (SPF) breeding facility] and/or breed differences (breed 1: T40 X Pietrain vs. breed 2: Large White X PIC426) were found for the acquisition of the holeboard task. Supplier effects are indistinguishable from effects due to breed.*

*Table S2.4. In all studies, the IVIs changed across trialblocks, but only in studies H7 and H8, group differences were found for the IVI. Ambient temperature during post-weaning housing and during testing (in °C) are also reported.*
*Abbreviations: n.a., not applicable; n.r., not reported.*

| Study | H1 | H3 | H4 | H5 | H6 | H7 | H8 | H9 | H10 | H11 | H12 |
|---|---|---|---|---|---|---|---|---|---|---|---|
| Changes in IVI across trialblocks? | yes | yes | yes | yes | yes | yes | yes | yes | yes | yes | yes |
| Differences between groups for IVI? | no | no | no | no | no | yes* | yes** | n.a. | no | no | no |
| Range of ambient temperature [C°] | 3 – 31 | n.r. | 20 | n.r. | -1 – 14 | 4 – 24[1]<br>8 – 34[2] | 8 – 20 | -3 – 22 | 8 – 38 | 0 – 27 | 3 – 34 |

*: *Interaction effect between the repeated measures factors Trialblocks and the factor Supplier/Breed.*
**: *Effect of the factor Housing Environment.*
[1]: *Range of ambient temperature [C°] measured from March up to and including May for the first batch of study H7.*
[2]: *Range of ambient temperature [C°] measures from April up to and including July for the second batch of study H7.*
