## Supplementary material for "On the intra-laboratory replicability of results in animal cognition research, obtained in animals that were not exposed to, or unaffected by, experimental manipulations — Exemplified by spatial learning in pigs *(Sus Scrofa Domesticus)* across 12 holeboard studies": S3 - supporting information

#### S3 - Supporting Information - Alternative exclusion and inclusion criteria for selecting pigs per study for the replicability analyses

### On the intra-laboratory replicability of results in animal cognition research, obtained in animals that were not exposed to, or unaffected by, experimental manipulations – Exemplified by spatial learning in pigs *(Sus Scrofa Domesticus)* across 12 holeboard studies

F. Josef van der Staay, Alexandra Antonides, Alexandra K. Dwulit, Lisa Fijn, Elise T. Gieling, Charlotte G. E. Grimberg-Henrici, Ellen Meijer, Sanne Roelofs, J.C.M. (Hans) Vernooij, Vivian Witjes & Saskia S. Arndt

#### Introduction

##### An alternative selection of pigs per study for the replicability analyses

It is not straightforward to determine which pigs should be included in the replicability analyses for each study. Our paper used two sets of selection criteria: first, all pigs that were not exposed to or affected by experimental manipulations were included; second, pigs of the Terra x Finnish Landrace x Duroc breed only. The broadest possible selection was applied to increase the number of pigs used in the replicability analyses. The second, stricter selection of one breed was used to ensure a homogeneous genetic background across studies. To exclude the effects of treatments and the influence of Göttingen minipigs, which differ significantly from commercial pig breeds, from the replicability analysis, an additional analysis was performed using samples selected with stricter exclusion and inclusion criteria. The results of this third analysis are presented and discussed below.

#### Material and Methods

For the present analysis, we applied a set of very strict inclusion/exclusion criteria. Pigs that met any of the exclusion criteria below were excluded from the replicability analyses (see Table S3.1.).

1) All pigs subjected to pharmacological treatment (allopurinol, H03) or fed an iron-deficient diet (H04, H07), even if the treatment had no detectable effects;
2) Groups of pigs with a low birth weight (H01, H03, H05 and H11), as this would have shifted the distribution of birth weights towards lower values;

3) Göttingen minipigs (H02), because they differ from commonly used commercial pig breeds in terms of characteristics such as lower body mass, an earlier onset of puberty, and smaller litters (Golledge and Richardson 2024).

Note that we consider all commercial pig breeds to be representative of pigs kept for food production. Consequently, we subjected the reference and working memory data of the pigs listed in the second column of table S3.1 to replicability analyses and excluded all pigs listed in the third column.

***Table S3.1: The inclusion and exclusion criteria that were used for the strictest selection of groups of pigs for the replicability analysis.*** *The study, the groups of pigs included and excluded from the analysis, and the resulting number of pigs analysed per study are depicted.*

*Abbreviations: LBW, low-birth weight; vLBW, very low-birth weight; NBW, normal-birth weight.*

| Study | Exclude from analysis | Include in analysis * | N |
|---|---|---|---|
| **H01** – LBW vs. NBW pigs | LBW pigs | NBW pigs | 9 |
| **H02** – Conventional pigs vs. minipigs | Göttingen minipigs | Duroc X (X Finnish Landrace) | 8 |
| **H03** – Effects of allopurinol in LBW and NBW pigs | Allopurinol treated and LBW pigs | Untreated NBW pigs | 9 |
| **H04** – Effects of an anaemic iron-deficient diet | Pigs fed an iron-deficient diet | Pigs fed a balanced diet | 10 |
| **H05** – vLBW vs. NBW pigs | vLBW pigs | NBW pigs | 10 |
| **H06** – Effects of a non-anaemic iron deficient diet | Pigs fed an iron-deficient diet | Pigs fed a balanced diet | 11 |
| **H07** – Effects of litter size and supplier/breed | None | All pigs | 20 |
| **H08** – Effects of environmental enrichment | None | All pigs | 20 |
| **H09** – Effects of social isolation | None | All pigs | 19 |
| **H10** – Males vs. females | None | All pigs | 20 |
| **H11** – LBW vs. NBW pigs | LBW pigs | NBW pig | 19 |
| **H12** – First vs. last born piglets in a litter | None | All pigs | 22 |
| | | Total N | **177** |

* *As NBW groups only include pigs near the mean of their litter, they may be more representative than a pooled group consisting of LBW and NBW pigs, since including LBW pigs skew the birth weight distribution.*

##### Statistical analysis

Initial WM and RM levels were measured using performance in the first trialblock. Linear learning progression was measured using the slope calculated across successive trial blocks of the holeboard task during the acquisition phase. These data were subjected to a one-way ANOVA with the factor Study (levels: H01 - H12), supplemented by Bonferroni post hoc pairwise comparisons between studies. All analyses were performed using jamovi, version 2.6.44 (Navarro and Foxcroft 2025; 'The jamovi project' 2025).

#### Results

##### Comparison of WM across all 12 studies

***Initial WM performance (trialblock 1):*** Performance in the first trialblock differed between studies ($F_{11,165}$=2.12, p<0.001). Bonferroni post hoc comparisons did not detect differences between studies for the initial WM performance

***Speed of WM learning (slope):*** The rate of WM learning differed between studies ($F_{11,165}=6.20$, $p<0.001$) Bonferroni post hoc comparisons between studies revealed that learning was faster in H01 than in all other studies, which did not differ from each other.

##### Comparison of RM across all 12 studies

***Initial RM performance (trialblock 1):*** The initial RM performance level differed between studies ($F_{11,165}=4.06$, $p<0.001$). Bonferroni post hoc comparisons revealed that this effect was due to a higher initial performance level in H09, which exceeded that of most other studies (H03, H04, H06- H08, H10-H12).

***Speed of RM learning (slope):*** The studies differed in the speed of RM learning ($F_{11,165}=7.67$ $p<0.001$). This effect was mainly caused by a lower learning rate in H05 than in most other studies (H02, H03, H06-H12), and to slower learning in H04 than in H11 and H12, as confirmed by Bonferroni post hoc comparison.

##### Correlations between WM and RM (see Table S3.2)

The initial WM performance level, i.e. performance in the first trial block, was negatively correlated with the speed of learning the WM component of the holeboard task, confirming the correlation analyses based on different selections of pigs for the repeatability analysis reported in the main paper: the better the initial WM performance, the shallower the increase in WM learning curves across trial blocks.

***Table S3.2: Pearson product-moment correlation coefficients ($r_{PM}$) were calculated for WM and RM performance between the first trial block (as an index of initial performance level) and the slope (as an index of learning).*** *The $r_{PM}$ values across all 12 studies (N = 177) are shown. This analysis only includes data from groups of pigs that meet the inclusion/exclusion criteria set out in Table S3.1.*

| | | WM in 1st trialblock | WM slope | RM in 1st trialblock | RM slope |
|---|---|---|---|---|---|
| **WM in 1st trialblock** (initial performance level) | $r_{PM}$ | 1 | | | |
| | *P* | | | | |
| **WM slope** (working memory slope; linear increase) | $r_{PM}$ | **-0.566** | 1 | | |
| | *P* | ***<.001*** | | | |
| **RM in 1st trialblock** (initial performance level) | $r_{PM}$ | -0.093 | 0.083 | 1 | |
| | *P* | *0.217* | *0.272* | | |
| **RM slope** (reference memory slope; linear increase) | $r_{PM}$ | 0.054 | 0.088 | -0.056 | 1 |
| | *P* | *0.478* | *0.242* | *0.457* | |

#### Discussion

We analysed the replicability of learning in the RM and WM components of pigs across 12 independent studies, determining which groups of pigs to include using two sets of criteria. All pigs that were not affected by or exposed to experimental manipulations qualified for inclusion in the broadest analysis, i.e. with the broadest set of criteria in the first analysis. The second analysis only included pigs of the Terra x Finnish Landrace x Duroc breed to ensure a homogeneous genetic background across studies.

In a third analysis, reported here, we apply an even stricter set of exclusion/inclusion criteria. Low-birth-weight pigs, pigs that had undergone pharmacological treatment (e.g. prenatal allopurinol via the sow) or dietary treatments (e.g. an iron-deficient diet during the nursing period) and Göttingen minipigs (as they differ from the commercial pig breeds used for meat production (Golledge and Richardson 2024)) were excluded from the repeatability analyses.

Irrespective of the exclusion/inclusion criteria used for selection the animals to be included in the replicability analysis, very similar results were obtained in the statistical analyses of the initial performance level of RM and WM, represented by the performance in the first trialblock, and of the speed of learning, represented by the slope across trialblock in the acquisition phase.

The correlation analysis looked at the performance of RM and WM in the first trialblock and the slope of their learning curves. This analysis supports the notion that RM and WM are independent components of spatial memory, as assessed in the holeboard task, corroborating observations that have also been made in rat studies (e.g., van der Staay 1999).

Considering the similarity of results of the replicability analyses using three different sets of exclusion/inclusion criteria, we conclude that the broadest set, namely pigs that were not affected by or exposed to experimental manipulations appears to be valid for assessing the replicability. It has the additional advantage of using the largest possible sample sizes for analysis.
