## Supplementary material for "On the intra-laboratory replicability of results in animal cognition research, obtained in animals that were not exposed to, or unaffected by, experimental manipulations — Exemplified by spatial learning in pigs *(Sus Scrofa Domesticus)* across 12 holeboard studies": S4 - supporting informatio

### S4 - Supporting information - Data used for replicability analysis of 12 holeboard studies in pigs

#### Variable names

**<<Note that variables followed by an "$" are alphanumeric; all other variables are numeric >>**

Study_no $
Breed
Sex $
Birth_W $
Treatment $
Pig_no $
WM_BL01 - WM_BL10
RM_BL01- RM_BL10
IVI_BL01 -IVI_BL10
Overall_WM
Overall_RM
WM_interc
WM_slope
RM_inter
RM_slope;

#### Labels for variables

**<< Blck = Block; acquis. = acquisition; lin. = linear >>**

WM_BL01 = 'Working memory, Blck 01, acquis.'
WM_BL02 = 'Working memory, Blck 02, acquis.'
WM_BL03 = 'Working memory, Blck 03, acquis.'

WM_BL04 = 'Working memory, Blck 04, acquis.'
WM_BL05 = 'Working memory, Blck 05, acquis.'
WM_BL06 = 'Working memory, Blck 06, acquis.'
WM_BL07 = 'Working memory, Blck 07, acquis.'
WM_BL08 = 'Working memory, Blck 08, acquis.'
WM_BL09 = 'Working memory, Blck 09, acquis.'
WM_BL10 = 'Working memory, Blck 10, acquis.'

RM_BL01 = 'Reference memory, Blck 01, acquis.'
RM_BL02 = 'Reference memory, Blck 02, acquis.'
RM_BL03 = 'Reference memory, Blck 03, acquis.'
RM_BL04 = 'Reference memory, Blck 04, acquis.'
RM_BL05 = 'Reference memory, Blck 05, acquis.'
RM_BL06 = 'Reference memory, Blck 06, acquis.'
RM_BL07 = 'Reference memory, Blck 07, acquis.'
RM_BL08 = 'Reference memory, Blck 08, acquis.'
RM_BL09 = 'Reference memory, Blck 09, acquis.'
RM_BL10 = 'Reference memory, Blck 10, acquis.'

IVI_BL01 = 'Log10 Inter-visit-interval, Blck 01, acquis.'
IVI_BL02 = 'Log10 Inter-visit-interval, Blck 02, acquis.'
IVI_BL03 = 'Log10 Inter-visit-interval, Blck 03, acquis.'
IVI_BL04 = 'Log10 Inter-visit-interval, Blck 04, acquis.'
IVI_BL05 = 'Log10 Inter-visit-interval, Blck 05, acquis.'
IVI_BL06 = 'Log10 Inter-visit-interval, Blck 06, acquis.'
IVI_BL07 = 'Log10 Inter-visit-interval, Blck 07, acquis.'
IVI_BL08 = 'Log10 Inter-visit-interval, Blck 08, acquis.'
IVI_BL09 = 'Log10 Inter-visit-interval, Blck 09, acquis.'
IVI_BL10 = 'Log10 Inter-visit-interval, Blck 10, acquis.'

Overall_WM = 'Mean WM performance over 10 trialblocks'
Overall_RM = 'Mean RM performance over 10 trialblocks'
WM_interc = 'Working memory intercept'
WM_slope = 'Working memory slope (lin. increase)'
RM_interc = 'Reference memory intercept'
RM_slope = 'Reference memory slope (lin. increase)';

### Data

**<< Note: A period in a further empty cell indicates "missing information" or "missing data" >>**

**<< Abbreviation: Fleetw., Fleetwood >>**

| | | | | | | | | | | | | | | | |
|---|---|---|---|---|---|---|---|---|---|---|---|---|---|---|---|
| H01 | 1 | F | NBW | . | 1a | 0.563 | 0.767 | 0.817 | 0.817 | 1.000 | 0.900 | . | . | . | . |
| 0.318 | 0.337 | 0.407 | 0.618 | 0.671 | 0.785 | . | . | . | . | | | | | | |
| 1.294 | 0.936 | 0.783 | 0.883 | 0.862 | 0.887 | . | . | . | . | . | . | 0.572 | 0.068 | 0.168 | 0.101 |
| H01 | 1 | F | LBW | . | 1b | 0.579 | 0.692 | 0.900 | 1.000 | 0.834 | 0.900 | . | . | . | . |
| 0.319 | 0.352 | 0.327 | 0.438 | 0.393 | 0.384 | . | . | . | . | | | | | | |
| 0.967 | 0.845 | 0.770 | 0.791 | 0.912 | 0.818 | . | . | . | . | . | . | 0.604 | 0.061 | 0.313 | 0.016 |
| H01 | 1 | F | NBW | . | 2a | 0.711 | 1.000 | 0.950 | 0.917 | 0.900 | 1.000 | . | . | . | . |
| 0.302 | 0.326 | 0.348 | 0.340 | 0.395 | 0.449 | . | . | . | . | | | | | | |
| 1.034 | 0.661 | 0.641 | 0.615 | 0.792 | 0.768 | . | . | . | . | . | . | 0.801 | 0.032 | 0.267 | 0.027 |
| H01 | 1 | F | LBW | . | 2b | 0.822 | 0.718 | 0.793 | 0.950 | 0.950 | 0.867 | . | . | . | . |
| 0.270 | 0.317 | 0.309 | 0.383 | 0.450 | 0.474 | . | . | . | . | | | | | | |
| 0.861 | 0.858 | 0.911 | 0.923 | 0.956 | 0.926 | . | . | . | . | . | . | 0.742 | 0.031 | 0.218 | 0.043 |
| H01 | 1 | F | NBW | . | 3a | 0.843 | 0.768 | 0.817 | 0.810 | 1.000 | 0.950 | . | . | . | . |
| 0.312 | 0.331 | 0.310 | 0.307 | 0.433 | 0.359 | . | . | . | . | | | | | | |
| 0.686 | 0.573 | 0.730 | 0.658 | 0.551 | 0.563 | . | . | . | . | . | . | 0.742 | 0.035 | 0.288 | 0.015 |
| H01 | 1 | F | LBW | . | 3b | 0.875 | 0.717 | 0.736 | 0.792 | 0.867 | 1.000 | . | . | . | . |
| 0.349 | 0.362 | 0.297 | 0.488 | 0.607 | 0.817 | . | . | . | . | | | | | | |

| | | | | | | | | | | | | | | | |
|---|---|---|---|---|---|---|---|---|---|---|---|---|---|---|---|
| 1.658 | 1.239 | 1.061 | 1.088 | 1.148 | 1.147 | . | . | . | . | . | . | 0.718 | 0.032 | 0.160 | 0.093 |
| H01 | 1 | F | NBW | . | 4a | 0.711 | 0.724 | 0.850 | 0.654 | 0.950 | 0.900 | . | . | . | . |
| 0.257 | 0.301 | 0.331 | 0.370 | 0.339 | 0.469 | . | . | . | . | | | | | | |
| 1.331 | 0.811 | 0.764 | 0.862 | 0.778 | 0.784 | . | . | . | . | . | . | 0.655 | 0.041 | 0.223 | 0.035 |
| H01 | 1 | F | LBW | . | 4b | 0.594 | 0.778 | 0.900 | 0.811 | 0.830 | 0.817 | . | . | . | . |
| 0.325 | 0.309 | 0.266 | 0.390 | 0.348 | 0.372 | . | . | . | . | | | | | | |
| 1.367 | 0.738 | 0.651 | 0.794 | 1.180 | 0.858 | . | . | . | . | . | . | 0.670 | 0.034 | 0.287 | 0.014 |
| H01 | 1 | F | NBW | . | 5a | 0.699 | 0.504 | 0.692 | 0.900 | 0.825 | 0.917 | . | . | . | . |
| 0.299 | 0.323 | 0.345 | 0.521 | 0.385 | 0.595 | . | . | . | . | | | | | | |
| 1.028 | 1.225 | 1.193 | 1.237 | 1.165 | 0.938 | . | . | . | . | . | . | 0.530 | 0.065 | 0.227 | 0.053 |
| H01 | 1 | F | LBW | . | 5b | 0.771 | 0.950 | 0.893 | 0.774 | 0.875 | 0.784 | . | . | . | . |
| 0.333 | 0.364 | 0.450 | 0.397 | 0.514 | 0.532 | . | . | . | . | | | | | | |
| 0.892 | 0.683 | 0.705 | 1.036 | 0.949 | 1.031 | . | . | . | . | . | . | 0.869 | -0.008 | 0.292 | 0.040 |
| H01 | 1 | F | NBW | . | 6a | 0.634 | 0.686 | 0.710 | 0.786 | 1.000 | 0.950 | . | . | . | . |
| 0.280 | 0.297 | 0.291 | 0.377 | 0.394 | 0.399 | . | . | . | . | | | | | | |
| 1.236 | 0.840 | 0.707 | 0.662 | 0.624 | 0.638 | . | . | . | . | . | . | 0.534 | 0.074 | 0.242 | 0.028 |
| H01 | 1 | F | LBW | . | 6b | 0.514 | 0.692 | 0.811 | 0.850 | 0.917 | 0.900 | . | . | . | . |
| 0.292 | 0.277 | 0.357 | 0.574 | 0.636 | 0.692 | . | . | . | . | | | | | | |
| 1.012 | 0.817 | 0.906 | 0.936 | 0.936 | 1.032 | . | . | . | . | . | . | 0.516 | 0.076 | 0.142 | 0.094 |
| H01 | 1 | F | NBW | . | 7a | 0.875 | 0.867 | 0.810 | 1.000 | 1.000 | 1.000 | . | . | . | . |
| 0.304 | 0.296 | 0.300 | 0.308 | 0.461 | 0.391 | . | . | . | . | | | | | | |
| 0.935 | 0.628 | 0.551 | 0.561 | 0.676 | 0.684 | . | . | . | . | . | . | 0.804 | 0.035 | 0.249 | 0.027 |
| H01 | 1 | F | LBW | . | 7b | 0.850 | 0.817 | 0.900 | 0.900 | 0.950 | 1.000 | . | . | . | . |
| 0.249 | 0.351 | 0.420 | 0.531 | 0.699 | 0.900 | . | . | . | . | | | | | | |
| 0.865 | 0.764 | 0.740 | 0.757 | 0.868 | 0.904 | . | . | . | . | . | . | 0.788 | 0.033 | 0.084 | 0.126 |
| H01 | 1 | F | NBW | . | 8a | 0.687 | 0.784 | 0.784 | 0.817 | 0.950 | 0.900 | . | . | . | . |
| 0.342 | 0.303 | 0.380 | 0.458 | 0.584 | 0.562 | . | . | . | . | | | | | | |
| 0.970 | 0.679 | 0.647 | 0.739 | 0.809 | 0.731 | . | . | . | . | . | . | 0.660 | 0.046 | 0.236 | 0.058 |
| H01 | 1 | F | LBW | . | 8b | 0.611 | 0.700 | 1.000 | 0.900 | 0.950 | 1.000 | . | . | . | . |
| 0.353 | 0.359 | 0.358 | 0.393 | 0.475 | 0.728 | . | . | . | . | | | | | | |
| 0.965 | 0.904 | 0.668 | 0.721 | 0.749 | 0.892 | . | . | . | . | . | . | 0.601 | 0.074 | 0.219 | 0.064 |
| H01 | 1 | F | NBW | . | 9a | 0.950 | 0.867 | 0.917 | 0.917 | 1.000 | 0.950 | . | . | . | . |
| 0.317 | 0.291 | 0.415 | 0.410 | 0.504 | 0.424 | . | . | . | . | | | | | | |
| 0.832 | 0.798 | 0.693 | 0.737 | 0.720 | 0.692 | . | . | . | . | . | . | 0.893 | 0.011 | 0.277 | 0.033 |
| H01 | 1 | F | LBW | . | 9b | 0.900 | 0.875 | 0.800 | 0.950 | 1.000 | 0.950 | . | . | . | . |
| 0.383 | 0.302 | 0.357 | 0.560 | 0.527 | 0.688 | . | . | . | . | | | | | | |
| 0.951 | 0.785 | 0.747 | 0.841 | 0.822 | 1.000 | . | . | . | . | . | . | 0.835 | 0.022 | 0.229 | 0.069 |
| H02 | 2 | F | . | . | C1 | 0.685 | 0.867 | 0.725 | 0.777 | 1.000 | 0.950 | 0.950 | 0.867 | 1.000 | 0.950 |
| 0.295 | 0.391 | 0.386 | 0.410 | 0.535 | 0.635 | 0.843 | 0.740 | 0.850 | 0.875 | | | | | | |
| . | . | . | . | . | . | . | . | . | . | 0.877 | 0.596 | 0.727 | 0.027 | 0.209 | 0.070 |
| H02 | 2 | F | . | . | C2 | 0.950 | 0.900 | 0.867 | 0.850 | 0.950 | 0.950 | 1.000 | 0.950 | 0.900 | 0.867 |
| 0.247 | 0.325 | 0.392 | 0.477 | 0.717 | 0.712 | 0.833 | 0.825 | 0.754 | 0.846 | | | | | | |
| . | . | . | . | . | . | . | . | . | . | 0.918 | 0.613 | 0.914 | 0.001 | 0.225 | 0.070 |
| H02 | 2 | F | . | . | C3 | 0.694 | 0.917 | 0.817 | 0.833 | 0.917 | 0.800 | 0.817 | 1.000 | 1.000 | 1.000 |
| 0.268 | 0.393 | 0.358 | 0.359 | 0.450 | 0.482 | 0.545 | 0.667 | 0.825 | 0.728 | | | | | | |
| . | . | . | . | . | . | . | . | . | . | 0.879 | 0.507 | 0.743 | 0.025 | 0.197 | 0.056 |
| H02 | 2 | F | . | . | C4 | 0.700 | 0.850 | 0.950 | 0.950 | 0.950 | 0.750 | 0.843 | 0.950 | 0.867 | 0.867 |
| 0.344 | 0.364 | 0.376 | 0.449 | 0.542 | 0.524 | 0.526 | 0.642 | 0.794 | 0.740 | | | | | | |
| . | . | . | . | . | . | . | . | . | . | 0.868 | 0.530 | 0.831 | 0.007 | 0.260 | 0.049 |
| H02 | 2 | F | . | . | C5 | 0.783 | 0.817 | 0.857 | 1.000 | 0.950 | 0.510 | 0.524 | 0.572 | 0.564 | 0.592 |
| 0.380 | 0.295 | 0.292 | 0.350 | 0.412 | 0.510 | 0.524 | 0.572 | 0.564 | 0.592 | | | | | | |
| . | . | . | . | . | . | . | . | . | . | 0.717 | 0.449 | 0.943 | -0.041 | 0.255 | 0.035 |
| H02 | 2 | F | . | . | C6 | 0.833 | 0.867 | 0.900 | 0.950 | 0.917 | 0.750 | 0.867 | 0.950 | 0.950 | 0.889 |
| 0.265 | 0.299 | 0.326 | 0.431 | 0.561 | 0.503 | 0.618 | 0.783 | 0.745 | 0.607 | | | | | | |
| . | . | . | . | . | . | . | . | . | . | 0.887 | 0.514 | 0.857 | 0.006 | 0.214 | 0.054 |
| H02 | 2 | F | . | . | C7 | 0.711 | 0.760 | 0.726 | 0.867 | 0.760 | 0.950 | 0.867 | 0.843 | 0.693 | 0.760 |
| 0.377 | 0.296 | 0.373 | 0.349 | 0.325 | 0.562 | 0.636 | 0.542 | 0.525 | 0.570 | | | | | | |
| . | . | . | . | . | . | . | . | . | . | 0.794 | 0.455 | 0.769 | 0.005 | 0.279 | 0.032 |
| H02 | 2 | F | . | . | C8 | 0.867 | 0.650 | 0.638 | 0.917 | 1.000 | 0.792 | 0.917 | 0.917 | 0.783 | 0.950 |
| 0.317 | 0.306 | 0.332 | 0.343 | 0.388 | 0.434 | 0.503 | 0.547 | 0.659 | 0.657 | | | | | | |
| . | . | . | . | . | . | . | . | . | . | 0.843 | 0.449 | 0.748 | 0.017 | 0.211 | 0.043 |
| H02 | 3 | F | . | . | M1 | 0.933 | 0.717 | 0.811 | 0.622 | 0.733 | 0.728 | 0.817 | 0.867 | 0.861 | 0.950 |
| 0.288 | 0.311 | 0.359 | 0.381 | 0.386 | 0.276 | 0.428 | 0.435 | 0.459 | 0.511 | | | | | | |
| . | . | . | . | . | . | . | . | . | . | 0.804 | 0.383 | 0.737 | 0.012 | 0.269 | 0.021 |
| H02 | 3 | F | . | . | M2 | 0.750 | 0.767 | 0.616 | 0.628 | 0.867 | 0.950 | 0.833 | 0.950 | 0.793 | 0.917 |
| 0.303 | 0.317 | 0.343 | 0.293 | 0.435 | 0.486 | 0.602 | 0.573 | 0.540 | 0.654 | | | | | | |
| . | . | . | . | . | . | . | . | . | . | 0.807 | 0.455 | 0.672 | 0.025 | 0.226 | 0.042 |
| H02 | 3 | F | . | . | M3 | 0.817 | 0.733 | 0.722 | 0.900 | 0.793 | 0.817 | 1.000 | 0.867 | 0.950 | 0.950 |
| 0.283 | 0.269 | 0.339 | 0.319 | 0.415 | 0.426 | 0.459 | 0.422 | 0.466 | 0.486 | | | | | | |
| . | . | . | . | . | . | . | . | . | . | 0.855 | 0.388 | 0.729 | 0.023 | 0.253 | 0.025 |
| H02 | 3 | F | . | . | M4 | 0.817 | 0.900 | 0.950 | 0.950 | 0.950 | 0.917 | 0.761 | 0.893 | 0.950 | 0.867 |
| 0.359 | 0.391 | 0.448 | 0.393 | 0.515 | 0.640 | 0.596 | 0.792 | 0.908 | 0.821 | | | | | | |
| . | . | . | . | . | . | . | . | . | . | 0.895 | 0.586 | 0.898 | -0.001 | 0.245 | 0.062 |
| H02 | 3 | F | . | . | M5 | 0.875 | 0.893 | 0.850 | 0.900 | 1.000 | 0.917 | 0.735 | 0.811 | 0.950 | 0.600 |
| 0.275 | 0.317 | 0.397 | 0.454 | 0.404 | 0.416 | 0.482 | 0.513 | 0.637 | 0.534 | | | | | | |
| . | . | . | . | . | . | . | . | . | . | 0.853 | 0.443 | 0.948 | -0.017 | 0.268 | 0.032 |
| H02 | 3 | F | . | . | M6 | 0.603 | 0.728 | 0.644 | 0.822 | 0.783 | 0.767 | 0.950 | 0.893 | 0.950 | 1.000 |
| 0.305 | 0.339 | 0.373 | 0.336 | 0.425 | 0.493 | 0.591 | 0.594 | 0.740 | 0.833 | | | | | | |
| . | . | . | . | . | . | . | . | . | . | 0.814 | 0.503 | 0.590 | 0.041 | 0.186 | 0.058 |

| | | | | | | | | | | | | | | | |
|---|---|---|---|---|---|---|---|---|---|---|---|---|---|---|---|
| H02 | 3 | F | . | . | M7 | 0.761 | 0.733 | 1.000 | 0.950 | 0.917 | 0.867 | 0.900 | 0.950 | 0.817 | 0.950 |
| 0.275 | 0.346 | 0.348 | 0.375 | 0.381 | 0.421 | 0.429 | 0.535 | 0.520 | 0.775. | | | | | | |
| | . | . | . | . | . | . | . | . | . | 0.884 | 0.440 | 0.823 | 0.011 | 0.212 | 0.042 |
| H02 | 3 | F | . | . | M8 | 0.728 | 0.893 | 0.694 | 0.478 | 0.670 | 0.668 | 0.560 | 0.685 | 0.893 | 0.775 |
| 0.319 | 0.307 | 0.341 | 0.316 | 0.347 | 0.344 | 0.336 | 0.377 | 0.424 | 0.503 | | | | | | |
| . | . | . | . | . | . | . | . | . | . | 0.704 | 0.361 | 0.683 | 0.004 | 0.271 | 0.016 |
| H03 | 1 | . | LBW | Allo | 1 | 0.775 | 0.900 | 0.728 | 0.868 | 0.918 | 0.828 | 0.843 | 0.678 | 0.608 | 0.850 |
| 0.250 | 0.343 | 0.338 | 0.310 | 0.290 | 0.363 | 0.360 | 0.403 | 0.390 | 0.553 | | | | | | |
| . | . | . | . | . | . | . | . | . | . | 0.799 | 0.360 | 0.859 | -0.011 | 0.240 | 0.022 |
| H03 | 1 | . | NBW | Allo | 11 | 0.623 | 0.633 | 0.800 | 0.653 | 0.760 | 0.868 | 0.900 | 0.578 | 0.893 | 1.000 |
| 0.213 | 0.260 | 0.330 | 0.358 | 0.315 | 0.338 | 0.355 | 0.413 | 0.495 | 0.508 | | | | | | |
| . | . | . | . | . | . | . | . | . | . | 0.771 | 0.358 | 0.606 | 0.030 | 0.201 | 0.029 |
| H03 | 1 | . | NBW | Cont | 12 | 0.900 | 0.750 | 0.818 | 0.900 | 0.743 | 0.868 | 0.900 | 0.893 | 0.843 | 0.950 |
| 0.283 | 0.343 | 0.378 | 0.453 | 0.540 | 0.808 | 0.825 | 0.918 | 0.820 | 0.958 | | | | | | |
| . | . | . | . | . | . | . | . | . | . | 0.856 | 0.632 | 0.803 | 0.010 | 0.182 | 0.082 |
| H03 | 1 | . | LBW | Cont | 13 | 0.868 | 0.893 | 0.685 | 0.868 | 0.678 | 0.900 | 0.728 | 0.825 | 0.950 | 1.000 |
| 0.295 | 0.373 | 0.375 | 0.458 | 0.445 | 0.495 | 0.685 | 0.593 | 0.768 | 0.918 | | | | | | |
| . | . | . | . | . | . | . | . | . | . | 0.839 | 0.540 | 0.769 | 0.013 | 0.201 | 0.062 |
| H03 | 1 | . | LBW | Allo | 14 | 0.745 | 0.695 | 0.810 | 0.950 | 0.950 | 0.918 | 0.888 | 0.630 | 1.000 | 0.868 |
| 0.293 | 0.350 | 0.320 | 0.425 | 0.278 | 0.360 | 0.468 | 0.438 | 0.463 | 0.455 | | | | | | |
| . | . | . | . | . | . | . | . | . | . | 0.845 | 0.385 | 0.775 | 0.013 | 0.283 | 0.018 |
| H03 | 1 | . | LBW | Cont | 15 | 0.900 | 0.753 | 0.843 | 0.950 | 0.918 | 1.000 | 0.793 | 0.950 | 0.900 | 0.868 |
| 0.300 | 0.308 | 0.315 | 0.418 | 0.518 | 0.540 | 0.780 | 0.670 | 0.765 | 0.895 | | | | | | |
| . | . | . | . | . | . | . | . | . | . | 0.887 | 0.551 | 0.858 | 0.005 | 0.169 | 0.069 |
| H03 | 1 | . | LBW | Cont | 16 | 0.750 | 0.785 | 0.868 | 1.000 | 0.850 | 0.918 | 0.918 | 0.900 | 0.950 | 0.950 |
| 0.303 | 0.325 | 0.370 | 0.543 | 0.535 | 0.648 | 0.818 | 0.783 | 0.738 | 0.793 | | | | | | |
| . | . | . | . | . | . | . | . | . | . | 0.889 | 0.585 | 0.791 | 0.018 | 0.242 | 0.062 |
| H03 | 1 | . | NBW | Allo | 17 | 0.843 | 0.670 | 0.635 | 0.638 | 0.790 | 0.728 | 0.800 | 0.760 | 0.868 | 0.818 |
| 0.313 | 0.318 | 0.340 | 0.320 | 0.400 | 0.415 | 0.593 | 0.553 | 0.758 | 0.720 | | | | | | |
| . | . | . | . | . | . | . | . | . | . | 0.755 | 0.473 | 0.681 | 0.013 | 0.185 | 0.052 |
| H03 | 1 | . | NBW | Cont | 18 | 0.618 | 0.893 | 0.918 | 0.728 | 0.950 | 1.000 | 1.000 | 1.000 | 0.868 | 0.850 |
| 0.295 | 0.378 | 0.353 | 0.463 | 0.480 | 0.498 | 0.620 | 0.760 | 0.683 | 0.823 | | | | | | |
| . | . | . | . | . | . | . | . | . | . | 0.882 | 0.535 | 0.776 | 0.019 | 0.221 | 0.057 |
| H03 | 1 | . | NBW | Cont | 19 | 0.703 | 0.685 | 0.868 | 0.918 | 0.950 | 0.950 | 0.728 | 0.810 | 0.950 | 0.900 |
| 0.253 | 0.353 | 0.368 | 0.525 | 0.470 | 0.535 | 0.573 | 0.633 | 0.825 | 0.785 | | | | | | |
| . | . | . | . | . | . | . | . | . | . | 0.846 | 0.532 | 0.754 | 0.017 | 0.211 | 0.058 |
| H03 | 1 | . | LBW | Allo | 2 | 0.785 | 0.875 | 0.850 | 0.835 | 0.735 | 0.843 | 0.810 | 0.785 | 0.760 | 1.000 |
| 0.345 | 0.308 | 0.343 | 0.505 | 0.348 | 0.473 | 0.723 | 0.678 | 0.523 | 0.768 | | | | | | |
| . | . | . | . | . | . | . | . | . | . | 0.828 | 0.501 | 0.800 | 0.005 | 0.242 | 0.047 |
| H03 | 1 | . | LBW | Allo | 20 | 0.550 | 0.695 | 0.593 | 0.900 | 0.918 | 0.950 | 0.950 | 1.000 | 0.843 | 0.950 |
| 0.253 | 0.325 | 0.358 | 0.353 | 0.460 | 0.618 | 0.718 | 0.603 | 0.598 | 0.733 | | | | | | |
| . | . | . | . | . | . | . | . | . | . | 0.835 | 0.502 | 0.606 | 0.042 | 0.211 | 0.053 |
| H03 | 1 | . | NBW | Cont | 21 | 0.793 | 0.785 | 0.868 | 0.843 | 0.718 | 0.875 | 0.860 | 0.875 | 1.000 | 1.000 |
| 0.290 | 0.313 | 0.350 | 0.345 | 0.400 | 0.445 | 0.480 | 0.623 | 0.725 | 0.743 | | | | | | |
| . | . | . | . | . | . | . | . | . | . | 0.862 | 0.471 | 0.741 | 0.022 | 0.179 | 0.053 |
| H03 | 1 | . | LBW | Cont | 22 | 0.810 | 1.000 | 0.843 | 0.950 | 0.950 | 0.843 | 0.900 | 0.950 | 0.850 | 0.950 |
| 0.288 | 0.408 | 0.310 | 0.513 | 0.573 | 0.455 | 0.548 | 0.525 | 0.568 | 0.700 | | | | | | |
| . | . | . | . | . | . | . | . | . | . | 0.905 | 0.489 | 0.888 | 0.003 | 0.292 | 0.036 |
| H03 | 1 | . | NBW | Allo | 23 | 0.845 | 0.850 | 0.818 | 0.875 | 0.950 | 0.918 | 0.900 | 0.950 | 1.000 | 0.673 |
| 0.263 | 0.278 | 0.343 | 0.345 | 0.405 | 0.585 | 0.495 | 0.720 | 0.868 | 0.535 | | | | | | |
| . | . | . | . | . | . | . | . | . | . | 0.878 | 0.484 | 0.871 | 0.001 | 0.180 | 0.055 |
| H03 | 1 | . | NBW | Allo | 24 | 0.900 | 0.668 | 0.610 | 0.735 | 0.950 | 0.900 | 1.000 | 0.868 | 1.000 | 0.918 |
| 0.310 | 0.440 | 0.455 | 0.460 | 0.550 | 0.735 | 0.728 | 0.623 | 0.678 | 0.838 | | | | | | |
| . | . | . | . | . | . | . | . | . | . | 0.855 | 0.582 | 0.704 | 0.027 | 0.307 | 0.050 |
| H03 | 1 | . | NBW | Cont | 25 | 0.850 | 0.900 | 0.868 | 0.843 | 0.918 | 0.900 | 0.950 | 0.868 | 0.950 | 0.893 |
| 0.233 | 0.408 | 0.415 | 0.460 | 0.350 | 0.653 | 0.518 | 0.565 | 0.673 | 0.688 | | | | | | |
| . | . | . | . | . | . | . | . | . | . | 0.894 | 0.496 | 0.859 | 0.006 | 0.257 | 0.043 |
| H03 | 1 | . | LBW | Cont | 3 | 0.643 | 0.743 | 0.900 | 0.668 | 0.618 | 0.918 | 0.950 | 0.900 | 0.785 | 0.868 |
| 0.370 | 0.323 | 0.443 | 0.368 | 0.358 | 0.393 | 0.448 | 0.390 | 0.450 | 0.450 | | | | | | |
| . | . | . | . | . | . | . | . | . | . | 0.799 | 0.399 | 0.683 | 0.021 | 0.345 | 0.010 |
| H03 | 1 | . | NBW | Allo | 5 | 0.750 | 0.793 | 0.778 | 0.710 | 0.850 | 0.850 | 0.893 | 0.950 | 0.868 | 0.868 |
| 0.145 | 0.273 | 0.295 | 0.400 | 0.305 | 0.388 | 0.408 | 0.393 | 0.535 | 0.440 | | | | | | |
| . | . | . | . | . | . | . | . | . | . | 0.831 | 0.358 | 0.731 | 0.018 | 0.189 | 0.031 |
| H03 | 1 | . | NBW | Cont | 6 | 0.825 | 0.868 | 0.950 | 1.000 | 0.950 | 0.950 | 0.800 | 1.000 | 0.893 | 0.918 |
| 0.255 | 0.315 | 0.333 | 0.445 | 0.403 | 0.618 | 0.713 | 0.660 | 0.945 | 0.865 | | | | | | |
| . | . | . | . | . | . | . | . | . | . | 0.915 | 0.555 | 0.893 | 0.004 | 0.137 | 0.076 |
| H03 | 1 | . | LBW | Allo | 8 | 0.805 | 1.000 | 0.628 | 0.893 | 0.818 | 0.785 | 0.710 | 0.818 | 0.893 | 0.850 |
| 0.285 | 0.335 | 0.363 | 0.335 | 0.330 | 0.323 | 0.368 | 0.448 | 0.505 | 0.485 | | | | | | |
| . | . | . | . | . | . | . | . | . | . | 0.820 | 0.378 | 0.819 | 0.000 | 0.261 | 0.021 |
| H03 | 1 | . | LBW | Cont | 9 | 0.608 | 0.778 | 0.678 | 0.893 | 1.000 | 1.000 | 0.893 | 0.900 | 0.918 | 1.000 |
| 0.293 | 0.348 | 0.273 | 0.413 | 0.508 | 0.725 | 0.570 | 0.733 | 0.698 | 0.868 | | | | | | |
| . | . | . | . | . | . | . | . | . | . | 0.867 | 0.543 | 0.679 | 0.034 | 0.189 | 0.064 |
| H03 | 1 | . | LBW | Allo | A1 | 0.643 | 0.900 | 0.520 | 0.918 | 0.793 | 0.785 | 0.778 | 0.900 | 0.950 | 1.000 |
| 0.233 | 0.380 | 0.388 | 0.400 | 0.433 | 0.403 | 0.590 | 0.698 | 0.775 | 0.850 | | | | | | |
| . | . | . | . | . | . | . | . | . | . | 0.819 | 0.515 | 0.651 | 0.031 | 0.168 | 0.063 |
| H03 | 1 | . | NBW | Allo | B2 | 0.620 | 0.875 | 0.825 | 0.900 | 0.918 | 1.000 | 0.950 | 0.900 | 0.728 | 1.000 |
| 0.270 | 0.358 | 0.370 | 0.330 | 0.400 | 0.453 | 0.540 | 0.543 | 0.533 | 0.868 | | | | | | |
| . | . | . | . | . | . | . | . | . | . | 0.872 | 0.466 | 0.772 | 0.018 | 0.195 | 0.049 |
| H03 | 1 | . | LBW | Allo | C1 | 0.500 | 0.750 | 0.768 | 0.535 | 0.900 | 0.793 | 0.918 | 0.800 | 0.900 | 0.818 |

| | | | | | | | | | | | | | | | |
|---|---|---|---|---|---|---|---|---|---|---|---|---|---|---|---|
| 0.058 | 0.320 | 0.420 | 0.415 | 0.540 | 0.538 | 0.578 | 0.725 | 0.648 | 0.813 | | | | | | |
| . | . | . | . | . | . | . | . | . | . | 0.768 | 0.505 | 0.598 | 0.031 | 0.135 | 0.067 |
| H03 | 1 | . | LBW | Allo | D2 | 0.625 | 0.843 | 0.760 | 0.918 | 0.840 | 0.810 | 0.793 | 0.950 | 1.000 | 0.918 |
| 0.308 | 0.295 | 0.335 | 0.388 | 0.490 | 0.555 | 0.630 | 0.590 | 0.618 | 0.588 | | | | | | |
| . | . | . | . | . | . | . | . | . | . | 0.846 | 0.480 | 0.703 | 0.026 | 0.251 | 0.041 |
| H03 | 1 | . | NBW | Allo | E1 | 0.793 | 0.715 | 0.713 | 0.735 | 0.810 | 0.868 | 0.893 | 0.918 | 0.918 | 0.868 |
| 0.285 | 0.333 | 0.345 | 0.313 | 0.400 | 0.383 | 0.613 | 0.473 | 0.493 | 0.455 | | | | | | |
| . | . | . | . | . | . | . | . | . | . | 0.823 | 0.409 | 0.701 | 0.022 | 0.270 | 0.025 |
| H03 | 1 | . | NBW | Allo | F2 | 0.710 | 0.635 | 0.563 | 0.800 | 0.825 | 0.950 | 0.950 | 0.950 | 0.918 | 0.950 |
| 0.313 | 0.358 | 0.398 | 0.423 | 0.378 | 0.403 | 0.550 | 0.618 | 0.755 | 0.763 | | | | | | |
| . | . | . | . | . | . | . | . | . | . | 0.825 | 0.496 | 0.603 | 0.040 | 0.218 | 0.051 |
| H03 | 1 | . | LBW | Allo | G1 | 0.785 | 0.610 | 0.638 | 0.728 | 0.950 | 0.893 | 1.000 | 1.000 | 1.000 | 0.950 |
| 0.325 | 0.323 | 0.378 | 0.483 | 0.583 | 0.820 | 0.868 | 0.850 | 1.000 | 0.800 | | | | | | |
| . | . | . | . | . | . | . | . | . | . | 0.855 | 0.643 | 0.629 | 0.041 | 0.217 | 0.077 |
| H03 | 1 | . | LBW | Allo | H2 | 0.775 | 0.503 | 0.548 | 0.843 | 1.000 | 0.900 | 0.618 | 0.630 | 0.775 | 0.950 |
| 0.385 | 0.323 | 0.275 | 0.385 | 0.498 | 0.553 | 0.488 | 0.530 | 0.573 | 0.718 | | | | | | |
| . | . | . | . | . | . | . | . | . | . | 0.754 | 0.473 | 0.650 | 0.019 | 0.260 | 0.039 |
| H03 | 1 | . | NBW | Allo | I1 | 0.760 | 0.710 | 0.760 | 0.868 | 0.918 | 0.950 | 0.785 | 0.950 | 0.900 | 1.000 |
| 0.308 | 0.273 | 0.358 | 0.375 | 0.353 | 0.485 | 0.680 | 0.698 | 0.785 | 0.760 | | | | | | |
| . | . | . | . | . | . | . | . | . | . | 0.860 | 0.507 | 0.719 | 0.026 | 0.160 | 0.063 |
| H03 | 1 | . | NBW | Allo | J2 | 0.525 | 0.810 | 1.000 | 0.810 | 0.950 | 0.843 | 0.950 | 0.900 | 0.918 | 0.743 |
| 0.385 | 0.443 | 0.510 | 0.628 | 0.743 | 0.798 | 0.820 | 0.845 | 0.835 | 0.800 | | | | | | |
| . | . | . | . | . | . | . | . | . | . | 0.845 | 0.681 | 0.761 | 0.015 | 0.388 | 0.053 |
| H03 | 1 | . | LBW | Cont | K1 | 0.868 | 0.793 | 0.768 | 0.660 | 0.950 | 0.950 | 1.000 | 0.875 | 0.893 | 0.695 |
| 0.280 | 0.285 | 0.298 | 0.400 | 0.355 | 0.398 | 0.455 | 0.498 | 0.470 | 0.403 | | | | | | |
| . | . | . | . | . | . | . | . | . | . | 0.845 | 0.384 | 0.822 | 0.004 | 0.264 | 0.022 |
| H03 | 1 | . | NBW | Cont | M1 | 0.918 | 0.950 | 0.743 | 0.825 | 0.950 | 0.950 | 1.000 | 1.000 | 0.793 | 0.868 |
| 0.393 | 0.313 | 0.330 | 0.393 | 0.498 | 0.473 | 0.603 | 0.578 | 0.793 | 0.765 | | | | | | |
| . | . | . | . | . | . | . | . | . | . | 0.900 | 0.514 | 0.891 | 0.002 | 0.228 | 0.052 |
| H03 | 1 | . | NBW | Cont | N2 | 0.613 | 0.650 | 0.810 | 0.843 | 0.918 | 0.835 | 0.950 | 0.675 | 0.950 | 0.950 |
| 0.330 | 0.328 | 0.355 | 0.423 | 0.518 | 0.733 | 0.660 | 0.695 | 0.733 | 0.875 | | | | | | |
| . | . | . | . | . | . | . | . | . | . | 0.819 | 0.565 | 0.663 | 0.029 | 0.219 | 0.063 |
| H03 | 1 | . | LBW | Cont | Q2 | 1.000 | 0.600 | 0.508 | 0.628 | 0.703 | 0.868 | 0.950 | 0.760 | 0.918 | 0.785 |
| 0.225 | 0.358 | 0.540 | 0.383 | 0.410 | 0.495 | 0.593 | 0.740 | 0.850 | 0.855 | | | | | | |
| . | . | . | . | . | . | . | . | . | . | 0.772 | 0.545 | 0.682 | 0.016 | 0.184 | 0.066 |
| H03 | 1 | . | NBW | Cont | R1 | 0.750 | 0.633 | 0.763 | 0.425 | 0.858 | 0.670 | 0.478 | 0.760 | 0.793 | 1.000 |
| 0.190 | 0.380 | 0.485 | 0.500 | 0.423 | 0.498 | 0.528 | 0.480 | 0.605 | 0.868 | | | | | | |
| . | . | . | . | . | . | . | . | . | . | 0.713 | 0.496 | 0.602 | 0.020 | 0.235 | 0.047 |
| H04 | 1 | M | . | Cont | 1 | 0.458 | 0.676 | 0.661 | 0.417 | 0.643 | 0.693 | 0.900 | 0.671 | 0.784 | 0.726 |
| 0.356 | 0.300 | 0.362 | 0.424 | 0.489 | 0.386 | 0.515 | 0.491 | 0.531 | 0.768 | | | | | | |
| 1.388 | 1.122 | 0.918 | 0.955 | 1.040 | 0.805 | 0.885 | 0.824 | 0.831 | 0.774 | 0.663 | 0.462 | 0.506 | 0.029 | 0.258 | 0.037 |
| H04 | 1 | M | . | Cont | 2 | 0.780 | 0.726 | 0.571 | 0.917 | 0.850 | 0.900 | 0.950 | 0.917 | 0.950 | 0.950 |
| 0.309 | 0.363 | 0.322 | 0.498 | 0.465 | 0.403 | 0.560 | 0.577 | 0.599 | 0.688 | | | | | | |
| 1.086 | 1.033 | 1.186 | 0.845 | 0.627 | 0.582 | 0.712 | 0.744 | 0.724 | 0.641 | 0.851 | 0.478 | 0.685 | 0.030 | 0.263 | 0.039 |
| H04 | 1 | M | . | Cont | 3 | 0.694 | 0.867 | 0.900 | 0.792 | 0.917 | 0.867 | 0.727 | 0.950 | 0.817 | 1.000 |
| 0.300 | 0.251 | 0.325 | 0.403 | 0.510 | 0.504 | 0.660 | 0.540 | 0.565 | 0.810 | | | | | | |
| 0.829 | 1.686 | 0.598 | 0.486 | 0.499 | 0.526 | 0.845 | 0.595 | 0.952 | 0.627 | 0.853 | 0.487 | 0.773 | 0.015 | 0.199 | 0.052 |
| H04 | 1 | M | . | Cont | 4 | 1.000 | 0.825 | 0.754 | 0.750 | 0.900 | 0.543 | 0.950 | 0.867 | 0.736 | 0.893 |
| 0.291 | 0.287 | 0.489 | 0.380 | 0.451 | 0.371 | 0.606 | 0.487 | 0.585 | 0.555 | | | | | | |
| 1.654 | 1.145 | 0.877 | 0.972 | 1.064 | 1.051 | 0.717 | 0.716 | 0.751 | 0.714 | 0.822 | 0.450 | 0.848 | -0.005 | 0.282 | 0.031 |
| H04 | 1 | M | . | Cont | 5 | 0.600 | 0.793 | 0.768 | 0.634 | 0.861 | 0.634 | 0.493 | 0.742 | 0.761 | 0.710 |
| 0.227 | 0.287 | 0.230 | 0.385 | 0.396 | 0.381 | 0.388 | 0.506 | 0.559 | 0.628 | | | | | | |
| 1.389 | 1.098 | 1.108 | 0.928 | 1.016 | 1.026 | 1.238 | 0.945 | 1.029 | 1.065 | 0.699 | 0.399 | 0.700 | 0.000 | 0.169 | 0.042 |
| H04 | 1 | M | . | Cont | 6 | 0.778 | 0.900 | 0.771 | 0.708 | 0.475 | 0.667 | 0.641 | 0.750 | 0.654 | 0.950 |
| 0.339 | 0.270 | 0.413 | 0.411 | 0.569 | 0.472 | 0.542 | 0.516 | 0.489 | 0.524 | | | | | | |
| 1.859 | 1.851 | 1.762 | 2.021 | 1.867 | 1.947 | 1.416 | 1.027 | 0.831 | 0.742 | 0.729 | 0.454 | 0.739 | -0.002 | 0.321 | 0.024 |
| H04 | 1 | M | . | Cont | 7 | 0.900 | 0.950 | 0.700 | 0.917 | 0.825 | 0.727 | 0.900 | 1.000 | 0.950 | 0.950 |
| 0.301 | 0.381 | 0.503 | 0.374 | 0.491 | 0.472 | 0.587 | 0.652 | 0.750 | 0.658 | | | | | | |
| 0.556 | 0.474 | 0.622 | 0.553 | 0.484 | 0.562 | 0.581 | 0.590 | 0.587 | 0.672 | 0.882 | 0.517 | 0.822 | 0.011 | 0.278 | 0.043 |
| H04 | 1 | M | . | Cont | 8 | 0.950 | 0.610 | 0.817 | 0.686 | 1.000 | 0.893 | 0.793 | 1.000 | 0.843 | 1.000 |
| 0.268 | 0.370 | 0.411 | 0.379 | 0.438 | 0.600 | 0.539 | 0.686 | 0.575 | 0.635 | | | | | | |
| 0.793 | 0.961 | 0.794 | 0.932 | 0.660 | 0.973 | 0.795 | 0.642 | 0.656 | 0.675 | 0.859 | 0.490 | 0.752 | 0.019 | 0.265 | 0.041 |
| H04 | 1 | M | . | Cont | 9 | 0.833 | 0.667 | 0.825 | 0.579 | 0.674 | 0.610 | 0.713 | 0.569 | 0.508 | 0.615 |
| 0.247 | 0.413 | 0.402 | 0.302 | 0.308 | 0.335 | 0.372 | 0.424 | 0.437 | 0.448 | | | | | | |
| 2.131 | 2.094 | 1.119 | 0.966 | 0.803 | 0.985 | 1.004 | 0.891 | 1.119 | 0.889 | 0.659 | 0.369 | 0.793 | -0.024 | 0.291 | 0.014 |
| H04 | 1 | M | . | Cont | 10 | 0.576 | 0.867 | 0.825 | 0.817 | 0.810 | 0.867 | 0.917 | 0.917 | 0.950 | 0.900 |
| 0.324 | 0.225 | 0.344 | 0.487 | 0.635 | 0.589 | 0.717 | 0.734 | 0.875 | 1.000 | | | | | | |
| 0.646 | 0.600 | 0.530 | 0.580 | 0.553 | 0.556 | 0.585 | 0.601 | 0.630 | 0.700 | 0.844 | 0.593 | 0.701 | 0.026 | 0.152 | 0.080 |
| H05 | 1 | F | NBW | . | 1 | 0.617 | 0.718 | 0.581 | 0.777 | 0.718 | 0.768 | 0.726 | 0.950 | 0.900 | 0.741 |
| 0.364 | 0.359 | 0.331 | 0.317 | 0.453 | 0.396 | 0.482 | 0.666 | 0.555 | 0.609 | | | | | | |
| 1.081 | 1.090 | 0.886 | 0.664 | 0.602 | 0.792 | 0.646 | 0.789 | 0.738 | 0.780 | 0.749 | 0.453 | 0.612 | 0.025 | 0.264 | 0.034 |
| H05 | 1 | M | NBW | . | 2 | 0.843 | 0.679 | 0.641 | 0.643 | 1.000 | 0.810 | 0.693 | 0.700 | 0.893 | 0.950 |
| 0.295 | 0.280 | 0.341 | 0.357 | 0.328 | 0.483 | 0.299 | 0.376 | 0.347 | 0.465 | | | | | | |
| 0.692 | 0.772 | 0.843 | 0.751 | 0.538 | 0.625 | 0.537 | 0.512 | 0.530 | 0.581 | 0.785 | 0.357 | 0.695 | 0.016 | 0.285 | 0.013 |
| H05 | 1 | M | NBW | . | 3 | 0.519 | 0.674 | 0.649 | 0.624 | 0.867 | 0.579 | 0.792 | 0.950 | 0.950 | 0.867 |
| 0.382 | 0.302 | 0.304 | 0.365 | 0.329 | 0.386 | 0.379 | 0.441 | 0.390 | 0.506 | | | | | | |
| 1.164 | 1.193 | 0.845 | 0.741 | 0.678 | 0.830 | 0.627 | 0.571 | 0.533 | 0.588 | 0.747 | 0.378 | 0.521 | 0.041 | 0.294 | 0.015 |
| H05 | 1 | M | NBW | . | 4 | 0.900 | 0.744 | 0.551 | 0.811 | 0.861 | 0.834 | 0.793 | 0.767 | 0.875 | 0.728 |
| 0.361 | 0.300 | 0.248 | 0.334 | 0.364 | 0.386 | 0.319 | 0.424 | 0.412 | 0.448 | | | | | | |

| | | | | | | | | | | | | | | | |
|---|---|---|---|---|---|---|---|---|---|---|---|---|---|---|---|
| 0.475 | 0.700 | 0.688 | 0.499 | 0.446 | 0.414 | 0.456 | 0.478 | 0.524 | 0.677 | 0.786 | 0.360 | 0.774 | 0.002 | 0.279 | 0.015 |
| H05 | 1 | F | NBW | . | 5 | 1.000 | 0.700 | 0.754 | 0.834 | 0.817 | 0.503 | 0.596 | 0.825 | 0.615 | 0.950 |
| 0.259 | 0.329 | 0.295 | 0.301 | 0.336 | 0.349 | 0.327 | 0.329 | 0.506 | 0.462 | | | | | | |
| 0.621 | 0.838 | 0.693 | 0.833 | 0.716 | 0.722 | 0.733 | 0.794 | 0.935 | 0.742 | 0.759 | 0.349 | 0.816 | -0.010 | 0.238 | 0.020 |
| H05 | 1 | F | NBW | . | 6 | 0.893 | 0.526 | 0.761 | 0.742 | 0.786 | 0.685 | 0.784 | 0.950 | 0.950 | 0.700 |
| 0.318 | 0.304 | 0.317 | 0.342 | 0.403 | 0.299 | 0.384 | 0.391 | 0.508 | 0.433 | | | | | | |
| 0.751 | 0.836 | 0.681 | 0.870 | 0.611 | 0.747 | 0.674 | 0.716 | 0.534 | 0.686 | 0.778 | 0.370 | 0.704 | 0.013 | 0.275 | 0.017 |
| H05 | 1 | M | NBW | . | 7 | 0.621 | 0.817 | 0.692 | 0.760 | 0.768 | 0.900 | 1.000 | 0.817 | 0.810 | 0.950 |
| 0.293 | 0.338 | 0.347 | 0.329 | 0.381 | 0.333 | 0.405 | 0.397 | 0.427 | 0.563 | | | | | | |
| 0.621 | 0.532 | 0.477 | 0.536 | 0.445 | 0.518 | 0.581 | 0.695 | 0.736 | 0.758 | 0.813 | 0.381 | 0.667 | 0.027 | 0.265 | 0.021 |
| H05 | 1 | M | NBW | . | 8 | 0.800 | 0.661 | 0.743 | 0.843 | 1.000 | 0.822 | 0.917 | 1.000 | 0.867 | 0.917 |
| 0.275 | 0.238 | 0.316 | 0.305 | 0.418 | 0.406 | 0.410 | 0.458 | 0.349 | 0.439 | | | | | | |
| 0.585 | 0.531 | 0.430 | 0.441 | 0.387 | 0.454 | 0.359 | 0.469 | 0.390 | 0.387 | 0.857 | 0.361 | 0.729 | 0.023 | 0.252 | 0.020 |
| H05 | 1 | M | NBW | . | 9 | 0.783 | 0.783 | 0.950 | 0.695 | 0.750 | 0.917 | 0.843 | 0.742 | 0.784 | 0.750 |
| 0.250 | 0.317 | 0.313 | 0.288 | 0.328 | 0.374 | 0.425 | 0.463 | 0.528 | 0.578 | | | | | | |
| 0.596 | 0.509 | 0.385 | 0.542 | 0.431 | 0.366 | 0.433 | 0.514 | 0.606 | 0.517 | 0.800 | 0.386 | 0.824 | -0.004 | 0.198 | 0.034 |
| H05 | 1 | F | NBW | . | 10 | 0.741 | 0.725 | 0.793 | 0.843 | 0.861 | 1.000 | 0.867 | 0.900 | 0.742 | 0.811 |
| 0.271 | 0.270 | 0.279 | 0.337 | 0.266 | 0.409 | 0.402 | 0.414 | 0.356 | 0.438 | | | | | | |
| 0.638 | 0.552 | 0.424 | 0.433 | 0.356 | 0.433 | 0.450 | 0.493 | 0.821 | 0.665 | 0.828 | 0.344 | 0.778 | 0.009 | 0.241 | 0.019 |
| H05 | 1 | F | vLBW | . | 11 | 0.650 | 0.760 | 0.867 | 0.492 | 0.754 | 0.861 | 0.950 | 0.917 | 0.742 | 0.917 |
| 0.337 | 0.324 | 0.377 | 0.383 | 0.428 | 0.497 | 0.539 | 0.536 | 0.611 | 0.693 | | | | | | |
| 0l.793 | 0.657 | 0.466 | 0.999 | 0.653 | 0.575 | 0.500 | 0.507 | 0.654 | 0.642 | 0.791 | 0.472 | 0.657 | 0.024 | 0.254 | 0.040 |
| H05 | 1 | M | vLBW | . | 12 | 0.800 | 0.734 | 0.552 | 0.792 | 0.632 | 0.736 | 0.810 | 0.917 | 1.000 | 0.750 |
| 0.292 | 0.324 | 0.376 | 0.445 | 0.395 | 0.496 | 0.525 | 0.592 | 0.760 | 0.703 | | | | | | |
| 0.652 | 0.631 | 0.591 | 0.581 | 0.688 | 0.558 | 0.668 | 0.624 | 0.666 | 0.668 | 0.772 | 0.491 | 0.659 | 0.021 | 0.218 | 0.050 |
| H05 | 1 | M | vLBW | . | 13 | 0.784 | 0.833 | 0.700 | 0.850 | 0.792 | 0.609 | 0.768 | 0.843 | 0.867 | 0.917 |
| 0.311 | 0.208 | 0.297 | 0.392 | 0.457 | 0.504 | 0.486 | 0.527 | 0.682 | 0.643 | | | | | | |
| 0.943 | 1.933 | 0.863 | 0.997 | 0.668 | 1.221 | 1.126 | 0.856 | 0.808 | 0.736 | 0.796 | 0.451 | 0.739 | 0.010 | 0.191 | 0.047 |
| H05 | 1 | F | vLBW | . | 14 | 0.844 | 0.750 | 0.621 | 0.636 | 0.760 | 0.675 | 0.867 | 0.811 | 0.709 | 0.917 |
| 0.278 | 0.334 | 0.355 | 0.325 | 0.461 | 0.419 | 0.621 | 0.698 | 0.636 | 0.577 | | | | | | |
| 1.176 | 1.440 | 0.790 | 0.730 | 0.632 | 0.654 | 0.677 | 0.661 | 0.807 | 0.609 | 0.759 | 0.470 | 0.695 | 0.012 | 0.225 | 0.045 |
| H05 | 1 | M | vLBW | . | 15 | 0.633 | 0.619 | 0.800 | 0.850 | 0.726 | 0.843 | 0.784 | 0.700 | 0.725 | 0.668 |
| 0.245 | 0.294 | 0.371 | 0.376 | 0.330 | 0.351 | 0.399 | 0.401 | 0.487 | 0.418 | | | | | | |
| 1.855 | 1.008 | 0.759 | 0.706 | 0.528 | 0.554 | 0.500 | 0.527 | 0.638 | 0.546 | 0.735 | 0.367 | 0.719 | 0.003 | 0.262 | 0.019 |
| H05 | 1 | F | vLBW | . | 16 | 0.834 | 0.810 | 0.577 | 0.817 | 0.900 | 0.867 | 0.792 | 0.867 | 0.843 | 0.950 |
| 0.254 | 0.291 | 0.324 | 0.296 | 0.295 | 0.468 | 0.431 | 0.505 | 0.376 | 0.517 | | | | | | |
| 0.611 | 0.552 | 0.761 | 0.465 | 0.357 | 0.500 | 0.596 | 0.549 | 0.528 | 0.619 | 0.825 | 0.376 | 0.738 | 0.016 | 0.228 | 0.027 |
| H05 | 1 | M | vLBW | . | 17 | 0.685 | 0.613 | 0.655 | 0.710 | 0.560 | 0.763 | 0.950 | 0.917 | 0.900 | 0.950 |
| 0.233 | 0.305 | 0.327 | 0.374 | 0.322 | 0.413 | 0.484 | 0.566 | 0.539 | 0.721 | | | | | | |
| 0.643 | 0.767 | 0.496 | 0.466 | 0.456 | 0.492 | 0.526 | 0.504 | 0.506 | 0.519 | 0.770 | 0.428 | 0.549 | 0.040 | 0.173 | 0.046 |
| H05 | 1 | F | vLBW | . | 18 | 0.643 | 0.736 | 0.700 | 0.778 | 0.666 | 0.718 | 0.917 | 0.810 | 1.000 | 0.917 |
| 0.222 | 0.316 | 0.355 | 0.333 | 0.371 | 0.417 | 0.395 | 0.495 | 0.460 | 0.421 | | | | | | |
| 1.477 | 0.877 | 0.802 | 0.628 | 0.624 | 0.507 | 0.514 | 0.523 | 0.502 | 0.519 | 0.788 | 0.378 | 0.610 | 0.032 | 0.254 | 0.023 |
| H05 | 1 | M | vLBW | . | 20 | 0.604 | 0.742 | 0.695 | 0.784 | 0.817 | 0.760 | 0.718 | 0.601 | 0.834 | 0.917 |
| 0.305 | 0.309 | 0.309 | 0.338 | 0.468 | 0.623 | 0.437 | 0.591 | 0.544 | 0.633 | | | | | | |
| 1.031 | 0.852 | 0.702 | 0.522 | 0.532 | 0.616 | 0.672 | 0.822 | 0.637 | 0.621 | 0.747 | 0.456 | 0.656 | 0.017 | 0.240 | 0.039 |
| H06 | 1 | M | . | 0 | 1 | 0.838 | 0.938 | 0.834 | 0.732 | 0.817 | 0.817 | 0.793 | 0.900 | 0.817 | 0.817 |
| 0.216 | 0.297 | 0.254 | 0.264 | 0.346 | 0.386 | 0.475 | 0.536 | 0.667 | 0.647 | | | | | | |
| 1.351 | 1.352 | 1.787 | 1.191 | 0.760 | 0.751 | 1.067 | 0.845 | 0.828 | 0.877 | 0.830 | 0.409 | 0.847 | -0.003 | 0.124 | 0.052 |
| H06 | 1 | F | . | 0 | 2 | 0.843 | 0.718 | 0.717 | 0.743 | 0.786 | 0.743 | 0.917 | 0.950 | 0.843 | 0.850 |
| 0.337 | 0.344 | 0.454 | 0.405 | 0.435 | 0.552 | 0.559 | 0.548 | 0.789 | 0.762 | | | | | | |
| 0.665 | 0.763 | 0.722 | 0.755 | 0.658 | 1.116 | 0.815 | 0.784 | 1.034 | 0.972 | 0.811 | 0.518 | 0.725 | 0.016 | 0.252 | 0.048 |
| H06 | 1 | M | . | 0 | 3 | 0.618 | 0.700 | 0.834 | 0.754 | 0.754 | 0.900 | 0.917 | 0.817 | 0.867 | 1.000 |
| 0.287 | 0.371 | 0.354 | 0.399 | 0.328 | 0.391 | 0.483 | 0.422 | 0.762 | 0.726 | | | | | | |
| 0.927 | 1.365 | 0.655 | 0.656 | 0.675 | 0.662 | 0.759 | 0.882 | 0.913 | 0.859 | 0.816 | 0.452 | 0.644 | 0.031 | 0.207 | 0.045 |
| H06 | 1 | F | . | 1 | 4 | 0.900 | 0.917 | 0.725 | 0.710 | 0.917 | 0.950 | 0.938 | 0.917 | 1.000 | 0.917 |
| 0.311 | 0.310 | 0.375 | 0.565 | 0.618 | 0.742 | 0.559 | 0.726 | 0.900 | 0.850 | | | | | | |
| 0.845 | 1.065 | 0.995 | 0.888 | 0.654 | 0.752 | 1.106 | 0.788 | 0.813 | 0.940 | 0.889 | 0.595 | 0.809 | 0.015 | 0.234 | 0.066 |
| H06 | 1 | F | . | 1 | 5 | 1.000 | 0.875 | 0.917 | 0.691 | 0.900 | 0.841 | 0.900 | 0.867 | 0.900 | 0.950 |
| 0.212 | 0.270 | 0.381 | 0.380 | 0.412 | 0.500 | 0.648 | 0.635 | 0.763 | 0.801 | | | | | | |
| 1.003 | 1.132 | 0.650 | 1.028 | 0.629 | 0.730 | 0.684 | 0.732 | 0.836 | 0.619 | 0.884 | 0.500 | 0.883 | 0.000 | 0.136 | 0.066 |
| H06 | 1 | M | . | 0 | 6 | 0.950 | 0.893 | 0.778 | 0.793 | 0.917 | 0.867 | 0.784 | 0.850 | 1.000 | 0.950 |
| 0.276 | 0.341 | 0.312 | 0.368 | 0.376 | 0.506 | 0.389 | 0.443 | 0.416 | 0.457 | | | | | | |
| 0.516 | 0.460 | 0.486 | 0.477 | 0.464 | 0.488 | 0.489 | 0.504 | 0.510 | 0.497 | 0.878 | 0.388 | 0.844 | 0.006 | 0.288 | 0.018 |
| H06 | 1 | M | . | 0 | 7 | 0.761 | 0.850 | 0.850 | 0.917 | 0.843 | 0.900 | 0.893 | 0.775 | 0.900 | 0.917 |
| 0.287 | 0.334 | 0.385 | 0.458 | 0.402 | 0.571 | 0.596 | 0.616 | 0.835 | 0.814 | | | | | | |
| 0.869 | 0.715 | 0.649 | 0.702 | 0.875 | 0.726 | 0.998 | 0.826 | 0.792 | 0.744 | 0.860 | 0.530 | 0.815 | 0.008 | 0.197 | 0.061 |
| H06 | 1 | M | . | 1 | 8 | 0.850 | 0.843 | 0.621 | 0.861 | 0.817 | 0.950 | 0.950 | 0.917 | 1.000 | 0.917 |
| 0.300 | 0.299 | 0.329 | 0.368 | 0.449 | 0.592 | 0.704 | 0.512 | 0.867 | 0.756 | | | | | | |
| 1.257 | 0.689 | 0.673 | 0.652 | 0.586 | 0.662 | 0.852 | 0.661 | 0.678 | 0.688 | 0.872 | 0.517 | 0.753 | 0.022 | 0.179 | 0.061 |
| H06 | 1 | F | . | 0 | 9 | 0.774 | 0.867 | 0.633 | 1.000 | 0.817 | 0.900 | 0.893 | 0.893 | 1.000 | 0.861 |
| 0.254 | 0.355 | 0.392 | 0.443 | 0.492 | 0.703 | 0.561 | 0.681 | 0.726 | 0.782 | | | | | | |
| 0.761 | 0.652 | 0.797 | 0.616 | 0.585 | 0.657 | 0.710 | 0.733 | 0.657 | 0.789 | 0.864 | 0.539 | 0.771 | 0.017 | 0.227 | 0.057 |
| H06 | 1 | F | . | 1 | 10 | 0.725 | 0.705 | 0.633 | 0.692 | 0.867 | 0.811 | 0.825 | 0.917 | 1.000 | 1.000 |
| 0.293 | 0.310 | 0.376 | 0.443 | 0.384 | 0.383 | 0.511 | 0.592 | 0.671 | 0.576 | | | | | | |
| 1.117 | 1.156 | 1.453 | 0.821 | 0.585 | 0.714 | 0.706 | 0.643 | 0.692 | 0.639 | 0.817 | 0.454 | 0.608 | 0.038 | 0.242 | 0.038 |
| H06 | 1 | F | . | 1 | 11 | 0.625 | 0.521 | 0.736 | 0.778 | 0.784 | 0.867 | 0.867 | 1.000 | 1.000 | 0.736 |
| 0.288 | 0.343 | 0.370 | 0.383 | 0.366 | 0.487 | 0.556 | 0.601 | 0.784 | 0.761 | | | | | | |
| 0.967 | 0.815 | 0.663 | 0.799 | 0.737 | 0.865 | 0.998 | 0.683 | 0.828 | 0.857 | 0.791 | 0.494 | 0.590 | 0.037 | 0.189 | 0.055 |

| | | | | | | | | | | | | | | | |
|---|---|---|---|---|---|---|---|---|---|---|---|---|---|---|---|
| H06 | 1 | F | . | 1 | 12 | 0.767 | 0.652 | 0.625 | 0.792 | 0.893 | 0.917 | 0.900 | 0.917 | 0.843 | 0.950 |
| 0.331 | 0.285 | 0.361 | 0.348 | 0.388 | 0.420 | 0.489 | 0.532 | 0.581 | 0.633 | | | | | | |
| 1.102 | 0.783 | 0.742 | 0.675 | 0.636 | 0.651 | 0.680 | 0.697 | 0.777 | 0.629 | 0.825 | 0.437 | 0.666 | 0.029 | 0.234 | 0.037 |
| H06 | 1 | F | . | 1 | 13 | 1.000 | 0.750 | 0.700 | 0.692 | 0.813 | 0.875 | 0.838 | 0.843 | 0.834 | 0.850 |
| 0.342 | 0.250 | 0.345 | 0.363 | 0.408 | 0.291 | 0.451 | 0.507 | 0.675 | 0.739 | | | | | | |
| 1.015 | 1.772 | 1.957 | 1.733 | 1.855 | 1.547 | 1.357 | 1.176 | 1.152 | 1.055 | 0.819 | 0.437 | 0.804 | 0.003 | 0.187 | 0.045 |
| H06 | 1 | M | . | 1 | 14 | 0.669 | 0.552 | 0.700 | 0.767 | 0.950 | 0.850 | 0.811 | 0.810 | 0.867 | 0.793 |
| 0.300 | 0.381 | 0.360 | 0.399 | 0.461 | 0.441 | 0.491 | 0.567 | 0.593 | 0.588 | | | | | | |
| 0.999 | 0.939 | 0.852 | 0.701 | 0.542 | 0.630 | 0.618 | 0.602 | 0.612 | 0.578 | 0.777 | 0.458 | 0.647 | 0.024 | 0.279 | 0.033 |
| H06 | 1 | F | . | 0 | 15 | 0.661 | 0.900 | 0.767 | 0.917 | 0.850 | 1.000 | 0.917 | 0.843 | 0.867 | 0.893 |
| 0.291 | 0.329 | 0.374 | 0.602 | 0.754 | 0.760 | 0.754 | 0.867 | 1.000 | 0.794 | | | | | | |
| 1.078 | 0.954 | 1.122 | 1.263 | 1.238 | 1.063 | 1.205 | 1.122 | 1.322 | 1.340 | 0.861 | 0.652 | 0.782 | 0.014 | 0.247 | 0.074 |
| H06 | 1 | F | . | 0 | 16 | 0.900 | 0.700 | 0.742 | 0.702 | 0.678 | 1.000 | 1.000 | 1.000 | 0.768 | 1.000 |
| 0.371 | 0.340 | 0.329 | 0.438 | 0.470 | 0.610 | 0.659 | 0.817 | 0.719 | 0.850 | | | | | | |
| 0.502 | 0.782 | 1.187 | 0.813 | 0.757 | 0.671 | 0.744 | 0.734 | 0.748 | 0.641 | 0.849 | 0.560 | 0.720 | 0.024 | 0.220 | 0.062 |
| H06 | 1 | M | . | 1 | 17 | 0.854 | 0.867 | 0.861 | 1.000 | 0.917 | 0.768 | 0.763 | 0.843 | 1.000 | 0.950 |
| 0.288 | 0.381 | 0.366 | 0.396 | 0.450 | 0.560 | 0.421 | 0.650 | 0.603 | 0.591 | | | | | | |
| 0.740 | 0.671 | 0.579 | 0.639 | 0.726 | 0.971 | 1.181 | 0.719 | 0.763 | 0.782 | 0.882 | 0.470 | 0.854 | 0.005 | 0.274 | 0.036 |
| H06 | 1 | M | . | 1 | 18 | 0.850 | 0.742 | 0.900 | 0.710 | 0.760 | 0.900 | 0.760 | 0.875 | 0.900 | 0.817 |
| 0.282 | 0.352 | 0.403 | 0.416 | 0.638 | 0.528 | 0.631 | 0.784 | 0.756 | 0.700 | | | | | | |
| 0.848 | 0.929 | 0.755 | 0.900 | 0.900 | 0.749 | 0.777 | 0.822 | 0.757 | 0.724 | 0.821 | 0.549 | 0.789 | 0.006 | 0.248 | 0.055 |
| H06 | 1 | M | . | 0 | 19 | 0.718 | 0.742 | 0.917 | 0.726 | 0.950 | 0.767 | 0.950 | 1.000 | 0.950 | 1.000 |
| 0.302 | 0.379 | 0.323 | 0.503 | 0.518 | 0.634 | 0.666 | 0.645 | 0.875 | 0.917 | | | | | | |
| 0.764 | 0.834 | 0.718 | 0.788 | 0.806 | 0.900 | 0.813 | 0.841 | 0.908 | 0.934 | 0.872 | 0.576 | 0.709 | 0.030 | 0.202 | 0.068 |
| H06 | 1 | F | . | 1 | 20 | 0.552 | 1.000 | 0.750 | 0.950 | 0.685 | 0.950 | 0.760 | 0.900 | 0.850 | 0.950 |
| 0.245 | 0.357 | 0.400 | 0.429 | 0.466 | 0.500 | 0.482 | 0.532 | 0.711 | 0.723 | | | | | | |
| 0.756 | 0.583 | 0.596 | 0.536 | 0.605 | 0.579 | 0.606 | 0.671 | 0.663 | 0.690 | 0.835 | 0.484 | 0.736 | 0.018 | 0.230 | 0.046 |
| H06 | 1 | F | . | 0 | 21 | 1.000 | 0.701 | 0.867 | 0.553 | 0.950 | 0.850 | 0.595 | 0.867 | 1.000 | 1.000 |
| 0.352 | 0.291 | 0.380 | 0.378 | 0.466 | 0.534 | 0.602 | 0.739 | 0.793 | 0.654 | | | | | | |
| 0.677 | 0.873 | 1.241 | 1.016 | 0.740 | 0.821 | 1.042 | 0.852 | 0.799 | 0.747 | 0.838 | 0.519 | 0.768 | 0.013 | 0.227 | 0.053 |
| H06 | 1 | M | . | 0 | 22 | 0.750 | 0.671 | 0.685 | 0.736 | 0.718 | 0.950 | 0.811 | 0.843 | 0.843 | 0.950 |
| 0.276 | 0.350 | 0.352 | 0.380 | 0.485 | 0.419 | 0.414 | 0.425 | 0.538 | 0.573 | | | | | | |
| 1.072 | 0.753 | 0.747 | 1.224 | 0.806 | 0.569 | 0.855 | 0.654 | 0.740 | 0.836 | 0.796 | 0.421 | 0.654 | 0.026 | 0.275 | 0.027 |
| H07 | 5 | M | . | Large | Aap | 0.900 | 0.900 | 0.918 | 0.793 | 0.868 | 0.710 | 0.950 | 0.635 | 1.000 | 0.918 |
| 0.275 | 0.383 | 0.345 | 0.323 | 0.308 | 0.370 | 0.393 | 0.375 | 0.448 | 0.525 | | | | | | |
| 0.805 | 0.691 | 0.636 | 0.910 | 0.842 | 0.667 | 0.646 | 0.802 | 0.702 | 0.589 | 0.859 | 0.374 | 0.867 | -0.001 | 0.270 | 0.019 |
| H07 | 5 | M | . | Large | Beer | 0.725 | 0.700 | 0.775 | 0.768 | 0.868 | 0.818 | 1.000 | 0.860 | 1.000 | 1.000 |
| 0.293 | 0.330 | 0.388 | 0.433 | 0.465 | 0.395 | 0.453 | 0.425 | 0.620 | 0.785 | | | | | | |
| 0.927 | 0.882 | 0.721 | 0.860 | 0.645 | 0.607 | 0.576 | 0.684 | 0.643 | 0.636 | 0.851 | 0.459 | 0.663 | 0.034 | 0.237 | 0.040 |
| H07 | 5 | M | . | Large | Co | 0.693 | 0.675 | 0.743 | 0.553 | 0.845 | 0.710 | 0.900 | 0.893 | 1.000 | 0.900 |
| 0.343 | 0.395 | 0.330 | 0.423 | 0.343 | 0.403 | 0.608 | 0.720 | 0.900 | 0.785 | | | | | | |
| 1.195 | 1.602 | 1.185 | 1.163 | 1.400 | 0.737 | 0.780 | 1.031 | 1.185 | 0.902 | 0.791 | 0.525 | 0.598 | 0.035 | 0.189 | 0.061 |
| H07 | 5 | M | . | Large | Don | 0.950 | 0.918 | 0.950 | 0.950 | 1.000 | 1.000 | 1.000 | 1.000 | 0.918 | 1.000 |
| 0.355 | 0.323 | 0.340 | 0.415 | 0.468 | 0.588 | 0.753 | 0.693 | 0.643 | 0.818 | | | | | | |
| 0.640 | 0.597 | 0.680 | 0.648 | 0.613 | 0.607 | 0.786 | 0.714 | 0.818 | 0.698 | 0.969 | 0.539 | 0.940 | 0.005 | 0.229 | 0.056 |
| H07 | 5 | M | . | Large | Ek | 0.850 | 0.950 | 0.900 | 0.843 | 0.900 | 0.793 | 0.893 | 1.000 | 1.000 | 0.900 |
| 0.273 | 0.400 | 0.333 | 0.508 | 0.450 | 0.465 | 0.485 | 0.703 | 0.678 | 0.875 | | | | | | |
| 1.042 | 0.811 | 0.691 | 0.750 | 0.695 | 0.691 | 0.705 | 0.786 | 0.707 | 0.790 | 0.903 | 0.517 | 0.858 | 0.008 | 0.211 | 0.056 |
| H07 | 5 | M | . | Small | Fetso | 0.735 | 0.843 | 0.793 | 0.918 | 0.743 | 0.668 | 0.693 | 0.868 | 0.950 | 0.950 |
| 0.258 | 0.313 | 0.338 | 0.335 | 0.403 | 0.438 | 0.583 | 0.650 | 0.693 | 0.845 | | | | | | |
| 0.655 | 0.630 | 0.687 | 0.614 | 0.647 | 0.620 | 0.630 | 0.673 | 0.650 | 0.780 | 0.816 | 0.485 | 0.739 | 0.014 | 0.142 | 0.062 |
| H07 | 5 | M | . | Small | Gap | 0.875 | 0.875 | 0.725 | 0.775 | 0.850 | 0.950 | 0.868 | 0.735 | 0.835 | 0.900 |
| 0.270 | 0.295 | 0.300 | 0.343 | 0.363 | 0.415 | 0.500 | 0.638 | 0.725 | 0.728 | | | | | | |
| 1.470 | 0.953 | 1.231 | 0.994 | 0.818 | 0.686 | 0.725 | 0.972 | 1.041 | 0.931 | 0.839 | 0.458 | 0.826 | 0.002 | 0.146 | 0.057 |
| H07 | 5 | M | . | Small | Hop | 0.633 | 0.735 | 0.825 | 0.800 | 0.750 | 0.723 | 0.628 | 0.875 | 0.628 | 0.733 |
| 0.310 | 0.353 | 0.350 | 0.307 | 0.460 | 0.337 | 0.365 | 0.353 | 0.413 | 0.505 | | | | | | |
| 1.356 | 1.396 | 1.430 | 1.880 | 1.962 | 1.419 | 1.270 | 1.214 | 1.360 | 1.171 | 0.733 | 0.375 | 0.738 | -0.001 | 0.300 | 0.014 |
| H07 | 5 | M | . | Small | Ips | 1.000 | 0.780 | 1.000 | 0.917 | 0.730 | 0.720 | 0.868 | 0.775 | 0.718 | 0.950 |
| 0.270 | 0.417 | 0.263 | 0.260 | 0.385 | 0.485 | 0.445 | 0.500 | 0.580 | 0.900 | | | | | | |
| 2.076 | 1.875 | 1.425 | 1.759 | 1.817 | 1.990 | 1.044 | 1.157 | 1.391 | 1.156 | 0.846 | 0.450 | 0.918 | -0.013 | 0.162 | 0.052 |
| H07 | 5 | M | . | Small | Jes | 0.785 | 0.710 | 1.000 | 0.868 | 1.000 | 0.710 | 0.900 | 0.950 | 0.918 | 0.950 |
| 0.318 | 0.315 | 0.440 | 0.408 | 0.403 | 0.470 | 0.495 | 0.663 | 0.678 | 0.713 | | | | | | |
| 1.223 | 0.806 | 0.587 | 0.616 | 0.554 | 0.613 | 0.629 | 0.662 | 0.740 | 0.770 | 0.879 | 0.490 | 0.796 | 0.015 | 0.239 | 0.046 |
| H07 | 6 | M | . | Large | Krok | 0.868 | 0.900 | 0.768 | 0.810 | 0.843 | 0.818 | 0.950 | 0.950 | 0.900 | 0.950 |
| 0.305 | 0.405 | 0.408 | 0.488 | 0.450 | 0.583 | 0.718 | 0.693 | 0.855 | 0.793 | | | | | | |
| 0.844 | 0.651 | 0.733 | 0.690 | 0.715 | 0.843 | 0.728 | 0.661 | 0.716 | 0.632 | 0.876 | 0.570 | 0.807 | 0.012 | 0.243 | 0.059 |
| H07 | 6 | M | . | Large | Lip | 0.893 | 0.668 | 0.950 | 0.775 | 0.918 | 0.703 | 0.653 | 0.768 | 0.875 | 0.950 |
| 0.253 | 0.308 | 0.353 | 0.395 | 0.450 | 0.578 | 0.535 | 0.633 | 0.680 | 0.768 | | | | | | |
| 0.713 | 0.741 | 0.615 | 0.613 | 0.631 | 0.797 | 0.684 | 0.744 | 0.736 | 0.731 | 0.815 | 0.495 | 0.799 | 0.003 | 0.189 | 0.056 |
| H07 | 6 | M | . | Large | Max | 0.938 | 0.738 | 0.868 | 0.675 | 0.793 | 0.950 | 0.645 | 1.000 | 0.843 | 0.950 |
| 0.180 | 0.313 | 0.435 | 0.415 | 0.558 | 0.775 | 0.638 | 0.760 | 0.818 | 0.875 | | | | | | |
| 1.578 | 1.569 | 1.274 | 1.192 | 1.227 | 1.554 | 1.468 | 1.365 | 1.290 | 1.219 | 0.840 | 0.577 | 0.787 | 0.010 | 0.167 | 0.075 |
| H07 | 6 | M | . | Large | Nano | 0.933 | 0.525 | 0.835 | 0.868 | 1.000 | 0.900 | 1.000 | 0.818 | 0.950 | 0.868 |
| 0.287 | 0.380 | 0.403 | 0.443 | 0.570 | 0.598 | 0.685 | 0.625 | 0.738 | 0.828 | | | | | | |
| 1.261 | 1.463 | 0.691 | 0.695 | 0.645 | 1.459 | 0.785 | 0.846 | 0.995 | 0.890 | 0.870 | 0.555 | 0.783 | 0.016 | 0.248 | 0.056 |
| H07 | 6 | M | . | Large | Otter | 0.893 | 0.700 | 0.875 | 1.000 | 1.000 | 0.743 | 0.900 | 0.843 | 0.950 | 1.000 |
| 0.293 | 0.240 | 0.308 | 0.413 | 0.425 | 0.360 | 0.480 | 0.653 | 0.690 | 0.818 | | | | | | |
| 0.824 | 1.042 | 0.667 | 0.683 | 0.723 | 0.972 | 0.917 | 1.030 | 0.932 | 0.870 | 0.890 | 0.468 | 0.824 | 0.012 | 0.143 | 0.059 |
| H07 | 6 | M | . | Small | Peer | 1.000 | 1.000 | 1.000 | 0.893 | 1.000 | 0.950 | 0.893 | 1.000 | 0.950 | 0.868 |

| | | | | | | | | | | | | | | | |
|---|---|---|---|---|---|---|---|---|---|---|---|---|---|---|---|
| 0.308 | 0.413 | 0.413 | 0.443 | 0.653 | 0.808 | 0.743 | 0.760 | 0.858 | 0.958 | | | | | | |
| 0.970 | 0.758 | 0.655 | 0.691 | 0.712 | 0.790 | 0.815 | 0.712 | 0.901 | 0.978 | 0.955 | 0.635 | 1.008 | -0.010 | 0.243 | 0.071 |
| H07 | 6 | M | . | Small | Quento | 0.950 | 0.900 | 0.950 | 0.900 | 0.950 | 1.000 | 1.000 | 0.950 | 1.000 | 0.950 |
| 0.288 | 0.290 | 0.330 | 0.348 | 0.360 | 0.468 | 0.545 | 0.603 | 0.613 | 0.775 | | | | | | |
| 0.558 | 0.558 | 0.542 | 0.521 | 0.452 | 0.518 | 0.558 | 0.640 | 0.622 | 0.753 | 0.955 | 0.462 | 0.920 | 0.006 | 0.172 | 0.053 |
| H07 | 6 | M | . | Small | Raaf | 0.868 | 0.710 | 0.668 | 0.760 | 0.825 | 0.900 | 0.918 | 1.000 | 0.713 | 1.000 |
| 0.250 | 0.350 | 0.338 | 0.405 | 0.358 | 0.495 | 0.575 | 0.618 | 0.655 | 0.793 | | | | | | |
| 0.765 | 0.710 | 0.886 | 0.921 | 0.579 | 0.542 | 0.925 | 0.628 | 1.195 | 0.987 | 0.836 | 0.484 | 0.722 | 0.021 | 0.181 | 0.055 |
| H07 | 6 | M | . | Small | Sil | 0.868 | 0.653 | 0.613 | 0.818 | 0.918 | 0.900 | 0.918 | 0.950 | 1.000 | 0.918 |
| 0.320 | 0.323 | 0.525 | 0.530 | 0.728 | 0.573 | 0.615 | 0.775 | 0.710 | 0.868 | | | | | | |
| 0.946 | 0.975 | 0.894 | 0.855 | 0.832 | 0.816 | 0.942 | 0.782 | 0.839 | 0.861 | 0.855 | 0.597 | 0.694 | 0.029 | 0.297 | 0.054 |
| H07 | 6 | M | . | Small | Tox | 0.838 | 0.950 | 0.768 | 0.810 | 1.000 | 0.718 | 0.620 | 0.950 | 0.868 | 0.900 |
| 0.310 | 0.490 | 0.738 | 0.613 | 0.850 | 0.670 | 0.420 | 0.723 | 0.908 | 0.908 | | | | | | |
| 1.339 | 0.862 | 1.437 | 1.096 | 0.820 | 1.032 | 1.082 | 0.820 | 0.816 | 0.871 | 0.842 | 0.663 | 0.841 | 0.000 | 0.414 | 0.045 |
| H08 | 1 | F | . | barren | 1 | 0.900 | 0.900 | 0.624 | 0.867 | 0.784 | 0.817 | 1.000 | 0.950 | 0.950 | 0.784 |
| 0.288 | 0.222 | 0.376 | 0.452 | 0.514 | 0.587 | 0.760 | 0.742 | 0.742 | 0.881 | | | | | | |
| 0.758 | 1.096 | 1.038 | 0.744 | 1.071 | 0.934 | 0.984 | 0.855 | 0.812 | 0.934 | 0.857 | 0.556 | 0.812 | 0.008 | 0.163 | 0.072 |
| H08 | 1 | F | . | barren | 10 | 0.950 | 0.900 | 0.850 | 0.850 | 0.900 | 0.850 | 1.000 | 1.000 | 0.834 | 0.825 |
| 0.251 | 0.389 | 0.330 | 0.362 | 0.471 | 0.545 | 0.577 | 0.659 | 0.486 | 0.664 | | | | | | |
| 0.755 | 0.671 | 0.632 | 0.665 | 0.723 | 0.665 | 0.690 | 0.776 | 0.817 | 1.076 | 0.896 | 0.473 | 0.911 | -0.003 | 0.248 | 0.041 |
| H08 | 1 | F | . | barren | 2 | 0.646 | 0.775 | 0.804 | 0.825 | 0.900 | 1.000 | 0.854 | 1.000 | 0.893 | 0.950 |
| 0.267 | 0.282 | 0.345 | 0.376 | 0.486 | 0.600 | 0.685 | 0.652 | 0.685 | 0.647 | | | | | | |
| 1.440 | 0.876 | 0.585 | 0.567 | 0.520 | 0.629 | 0.722 | 0.889 | 0.737 | 0.661 | 0.865 | 0.502 | 0.707 | 0.029 | 0.208 | 0.053 |
| H08 | 1 | F | . | barren | 3 | 1.000 | 0.900 | 1.000 | 0.867 | 1.000 | 0.950 | 0.867 | 0.950 | 0.950 | 0.950 |
| 0.247 | 0.260 | 0.287 | 0.411 | 0.470 | 0.566 | 0.526 | 0.736 | 0.738 | 0.556 | | | | | | |
| 0.582 | 0.695 | 0.631 | 0.590 | 0.625 | 0.595 | 0.724 | 0.679 | 0.709 | 0.684 | 0.943 | 0.480 | 0.957 | -0.002 | 0.186 | 0.053 |
| H08 | 1 | F | . | barren | 4 | 0.917 | 0.950 | 0.900 | 0.900 | 0.867 | 0.950 | 0.811 | 0.793 | 1.000 | 0.950 |
| 0.302 | 0.416 | 0.416 | 0.538 | 0.487 | 0.577 | 0.589 | 0.603 | 0.843 | 0.768 | | | | | | |
| 0.805 | 0.778 | 0.820 | 0.755 | 0.809 | 0.868 | 0.693 | 0.884 | 0.834 | 0.910 | 0.904 | 0.554 | 0.906 | 0.000 | 0.275 | 0.051 |
| H08 | 1 | F | . | barren | 5 | 0.600 | 0.588 | 0.817 | 0.867 | 1.000 | 1.000 | 0.950 | 0.810 | 0.811 | 0.893 |
| 0.261 | 0.336 | 0.298 | 0.468 | 0.440 | 0.493 | 0.427 | 0.584 | 0.480 | 0.530 | | | | | | |
| 0.959 | 0.830 | 0.642 | 0.482 | 0.690 | 0.660 | 0.530 | 0.868 | 0.699 | 0.803 | 0.833 | 0.432 | 0.686 | 0.027 | 0.272 | 0.029 |
| H08 | 1 | F | . | barren | 6 | 0.834 | 0.875 | 0.825 | 0.950 | 0.893 | 0.843 | 1.000 | 0.950 | 0.950 | 0.888 |
| 0.253 | 0.354 | 0.438 | 0.601 | 0.485 | 0.560 | 0.636 | 0.775 | 0.858 | 0.764 | | | | | | |
| 0.662 | 0.570 | 0.604 | 0.697 | 0.613 | 0.850 | 0.710 | 0.803 | 0.960 | 0.851 | 0.901 | 0.572 | 0.843 | 0.011 | 0.239 | 0.061 |
| H08 | 1 | F | . | barren | 7 | 0.950 | 0.843 | 0.867 | 0.867 | 0.834 | 0.686 | 0.843 | 0.950 | 0.900 | 0.950 |
| 0.293 | 0.334 | 0.375 | 0.393 | 0.483 | 0.350 | 0.456 | 0.742 | 0.582 | 0.740 | | | | | | |
| 0.632 | 0.639 | 0.522 | 0.516 | 0.587 | 0.759 | 0.608 | 0.647 | 0.597 | 0.725 | 0.869 | 0.475 | 0.849 | 0.004 | 0.220 | 0.046 |
| H08 | 1 | F | . | barren | 8 | 1.000 | 0.867 | 0.950 | 0.950 | 0.900 | 1.000 | 0.817 | 1.000 | 0.900 | 0.867 |
| 0.316 | 0.384 | 0.378 | 0.411 | 0.396 | 0.427 | 0.587 | 0.634 | 0.797 | 0.521 | | | | | | |
| 0.567 | 0.618 | 0.444 | 0.501 | 0.652 | 0.536 | 0.586 | 0.663 | 0.719 | 0.950 | 0.925 | 0.485 | 0.959 | -0.006 | 0.266 | 0.040 |
| H08 | 1 | F | . | barren | 9 | 0.604 | 0.754 | 0.692 | 0.767 | 0.780 | 0.825 | 0.830 | 0.817 | 1.000 | 1.000 |
| 0.317 | 0.277 | 0.413 | 0.472 | 0.594 | 0.857 | 0.763 | 0.825 | 0.800 | 0.850 | | | | | | |
| 1.385 | 1.191 | 1.130 | 1.384 | 1.235 | 1.406 | 0.990 | 0.896 | 0.887 | 0.824 | 0.807 | 0.617 | 0.602 | 0.037 | 0.228 | 0.071 |
| H08 | 1 | F | . | enr. | 11 | 0.760 | 0.917 | 1.000 | 1.000 | 0.950 | 0.950 | 0.792 | 1.000 | 0.875 | 0.917 |
| 0.292 | 0.309 | 0.392 | 0.579 | 0.625 | 0.908 | 0.709 | 0.917 | 0.882 | 0.886 | | | | | | |
| 0.693 | 0.616 | 0.630 | 0.608 | 0.672 | 0.721 | 0.736 | 0.767 | 0.779 | 0.794 | 0.916 | 0.650 | 0.899 | 0.003 | 0.228 | 0.077 |
| H08 | 1 | F | . | enr. | 12 | 0.742 | 0.692 | 0.900 | 0.800 | 0.834 | 0.811 | 0.950 | 1.000 | 0.900 | 1.000 |
| 0.305 | 0.283 | 0.332 | 0.357 | 0.350 | 0.408 | 0.444 | 0.542 | 0.590 | 0.578 | | | | | | |
| 0.743 | 0.815 | 0.633 | 0.678 | 0.677 | 0.695 | 0.656 | 0.600 | 0.600 | 0.684 | 0.863 | 0.419 | 0.706 | 0.029 | 0.220 | 0.036 |
| H08 | 1 | F | . | enr. | 13 | 0.575 | 1.000 | 0.843 | 0.867 | 0.791 | 0.850 | 1.000 | 1.000 | 1.000 | 0.867 |
| 0.286 | 0.396 | 0.426 | 0.544 | 0.520 | 0.599 | 0.784 | 0.950 | 0.734 | 0.888 | | | | | | |
| 1.110 | 0.643 | 0.631 | 0.615 | 0.504 | 0.692 | 0.612 | 0.614 | 0.664 | 0.716 | 0.879 | 0.613 | 0.750 | 0.023 | 0.239 | 0.068 |
| H08 | 1 | F | . | enr. | 14 | 0.950 | 0.774 | 0.950 | 0.917 | 0.900 | 1.000 | 0.893 | 0.950 | 0.950 | 1.000 |
| 0.337 | 0.312 | 0.402 | 0.562 | 0.515 | 0.850 | 0.825 | 0.823 | 0.958 | 1.000 | | | | | | |
| 0.582 | 0.820 | 0.563 | 0.685 | 0.853 | 0.735 | 0.784 | 0.796 | 0.919 | 0.804 | 0.928 | 0.658 | 0.871 | 0.010 | 0.201 | 0.083 |
| H08 | 1 | F | . | enr. | 15 | 0.686 | 1.000 | 0.825 | 1.000 | 0.950 | 1.000 | 0.888 | 1.000 | 1.000 | 1.000 |
| 0.331 | 0.351 | 0.472 | 0.676 | 0.656 | 0.643 | 0.806 | 0.950 | 0.900 | 0.950 | | | | | | |
| 0.666 | 0.604 | 0.617 | 0.490 | 0.522 | 0.607 | 0.672 | 0.736 | 0.559 | 0.656 | 0.935 | 0.674 | 0.821 | 0.021 | 0.268 | 0.074 |
| H08 | 1 | F | . | enr. | 16 | 0.671 | 0.834 | 0.917 | 0.950 | 0.730 | 0.875 | 1.000 | 0.843 | 0.810 | 0.950 |
| 0.350 | 0.354 | 0.388 | 0.431 | 0.468 | 0.431 | 0.556 | 0.692 | 0.629 | 0.768 | | | | | | |
| 0.789 | 0.588 | 0.611 | 0.591 | 0.632 | 0.639 | 0.563 | 0.680 | 0.742 | 0.676 | 0.858 | 0.507 | 0.782 | 0.014 | 0.255 | 0.046 |
| H08 | 1 | F | . | enr. | 17 | 0.900 | 0.850 | 0.950 | 1.000 | 1.000 | 0.893 | 1.000 | 1.000 | 0.950 | 0.900 |
| 0.285 | 0.335 | 0.432 | 0.545 | 0.601 | 0.784 | 0.775 | 0.767 | 0.845 | 0.750 | | | | | | |
| 0.687 | 0.716 | 0.606 | 0.546 | 0.580 | 0.674 | 0.693 | 0.761 | 0.809 | 0.994 | 0.944 | 0.612 | 0.916 | 0.005 | 0.268 | 0.062 |
| H08 | 1 | F | . | enr. | 18 | 0.825 | 0.825 | 0.817 | 0.917 | 0.950 | 0.900 | 0.917 | 0.825 | 0.850 | 0.950 |
| 0.307 | 0.349 | 0.336 | 0.374 | 0.425 | 0.622 | 0.571 | 0.599 | 0.879 | 0.609 | | | | | | |
| 0.596 | 0.787 | 0.649 | 0.634 | 0.609 | 0.753 | 0.639 | 0.599 | 0.734 | 0.668 | 0.878 | 0.507 | 0.834 | 0.008 | 0.223 | 0.052 |
| H08 | 1 | F | . | enr. | 19 | 0.811 | 0.680 | 0.817 | 0.875 | 0.867 | 0.834 | 0.834 | 1.000 | 0.888 | 1.000 |
| 0.447 | 0.264 | 0.389 | 0.486 | 0.516 | 0.479 | 0.523 | 0.760 | 0.624 | 0.700 | | | | | | |
| 0.608 | 0.766 | 0.566 | 0.607 | 0.614 | 0.706 | 0.641 | 0.738 | 0.734 | 0.842 | 0.860 | 0.519 | 0.730 | 0.024 | 0.294 | 0.041 |
| H08 | 1 | F | . | enr. | 20 | 0.867 | 0.867 | 0.843 | 0.867 | 0.843 | 0.950 | 0.817 | 1.000 | 1.000 | 1.000 |
| 0.297 | 0.370 | 0.326 | 0.430 | 0.422 | 0.455 | 0.519 | 0.793 | 0.760 | 0.810 | | | | | | |
| 0.622 | 0.498 | 0.645 | 0.552 | 0.576 | 0.597 | 0.600 | 0.662 | 0.687 | 0.684 | 0.905 | 0.518 | 0.809 | 0.017 | 0.185 | 0.060 |
| H09 | 2 | F | . | . | Asis | 0.810 | 0.810 | 0.900 | 0.917 | 0.933 | 0.843 | 0.893 | 0.950 | 0.950 | 1.000 |
| 0.337 | 0.423 | 0.486 | 0.556 | 0.794 | 0.738 | 0.719 | 0.789 | 0.718 | 0.784 | | | | | | |
| 0.649 | 0.651 | 0.643 | 0.590 | 0.734 | 0.809 | 0.759 | 0.741 | 0.983 | 0.732 | 0.900 | 0.634 | 0.808 | 0.017 | 0.366 | 0.049 |
| H09 | 2 | F | . | . | Babe | 0.834 | 0.750 | 0.676 | 0.725 | 0.950 | 0.950 | 0.861 | 0.843 | 0.867 | 0.793 |
| 0.384 | 0.341 | 0.420 | 0.545 | 0.536 | 0.476 | 0.537 | 0.624 | 0.602 | 0.612 | | | | | | |

| | | | | | | | | | | | | | | | |
|---|---|---|---|---|---|---|---|---|---|---|---|---|---|---|---|
| 0.504 | 0.547 | 0.646 | 0.596 | 0.745 | 0.742 | 0.734 | 0.696 | 0.641 | 0.673 | 0.825 | 0.507 | 0.769 | 0.010 | 0.347 | 0.029 |
| H09 | 2 | F | . | . | Chipo | 0.917 | 0.710 | 0.900 | 0.900 | 0.843 | 1.000 | 0.817 | 0.867 | 0.850 | 1.000 |
| 0.484 | 0.349 | 0.407 | 0.670 | 0.536 | 0.465 | 0.585 | 0.672 | 0.724 | 0.950 | | | | | | |
| 0.620 | 0.752 | 0.673 | 0.621 | 0.657 | 0.625 | 0.630 | 0.886 | 0.842 | 0.636 | 0.880 | 0.584 | 0.831 | 0.009 | 0.323 | 0.047 |
| H09 | 2 | F | . | . | Dumbo | 0.718 | 0.867 | 0.784 | 0.867 | 0.678 | 0.950 | 0.817 | 0.950 | 0.675 | 0.900 |
| 0.322 | 0.411 | 0.538 | 0.574 | 0.498 | 0.546 | 0.617 | 0.810 | 0.645 | 0.691 | | | | | | |
| 0.881 | 0.661 | 0.712 | 0.643 | 0.773 | 0.719 | 0.736 | 0.710 | 0.878 | 0.690 | 0.820 | 0.565 | 0.779 | 0.008 | 0.349 | 0.039 |
| H09 | 2 | F | . | . | Fayola | 0.843 | 0.825 | 1.000 | 0.736 | 1.000 | 0.893 | 0.867 | 0.900 | 0.950 | 0.850 |
| 0.333 | 0.549 | 0.504 | 0.561 | 0.556 | 0.635 | 0.682 | 0.729 | 0.716 | 0.815 | | | | | | |
| 0.794 | 0.801 | 0.748 | 0.880 | 0.833 | 0.870 | 0.830 | 0.765 | 0.800 | 0.897 | 0.886 | 0.608 | 0.862 | 0.004 | 0.372 | 0.043 |
| H09 | 2 | F | . | . | Gwenn | 0.601 | 0.728 | 0.710 | 0.728 | 0.825 | 0.933 | 1.000 | 0.917 | 0.917 | 0.950 |
| 0.335 | 0.439 | 0.438 | 0.484 | 0.559 | 0.778 | 0.710 | 0.695 | 0.854 | 0.879 | | | | | | |
| 0.730 | 0.688 | 0.656 | 0.753 | 0.732 | 0.777 | 0.788 | 0.764 | 0.715 | 0.685 | 0.831 | 0.617 | 0.617 | 0.039 | 0.284 | 0.061 |
| H09 | 2 | F | . | . | Hakuna | 0.889 | 0.857 | 0.792 | 0.742 | 0.893 | 1.000 | 0.893 | 1.000 | 0.950 | 1.000 |
| 0.343 | 0.506 | 0.487 | 0.549 | 0.598 | 0.652 | 0.613 | 0.734 | 0.685 | 0.817 | | | | | | |
| 0.830 | 0.821 | 0.656 | 0.704 | 0.713 | 0.697 | 0.683 | 0.611 | 0.754 | 0.744 | 0.902 | 0.598 | 0.793 | 0.020 | 0.365 | 0.042 |
| H09 | 2 | F | . | . | Ilori | 0.725 | 0.917 | 0.726 | 0.843 | 0.843 | 0.850 | 1.000 | 1.000 | 0.950 | 1.000 |
| 0.295 | 0.465 | 0.532 | 0.601 | 0.754 | 0.646 | 0.635 | 0.667 | 0.845 | 0.867 | | | | | | |
| 0.496 | 0.479 | 0.517 | 0.549 | 0.649 | 0.618 | 0.617 | 0.579 | 0.607 | 0.549 | 0.885 | 0.630 | 0.734 | 0.028 | 0.348 | 0.051 |
| H09 | 2 | F | . | . | Juju | 0.850 | 0.704 | 0.702 | 1.000 | 1.000 | 0.950 | 1.000 | 0.933 | 1.000 | 1.000 |
| 0.307 | 0.376 | 0.478 | 0.576 | 0.636 | 0.771 | 0.800 | 0.742 | 0.793 | 0.867 | | | | | | |
| 0.779 | 0.969 | 0.922 | 0.642 | 0.752 | 0.678 | 0.706 | 0.778 | 0.679 | 0.625 | 0.914 | 0.635 | 0.763 | 0.027 | 0.298 | 0.061 |
| H09 | 2 | F | . | . | Knorr | 0.867 | 0.744 | 0.775 | 0.867 | 0.867 | 0.950 | 0.810 | 0.950 | 0.843 | 0.950 |
| 0.378 | 0.362 | 0.519 | 0.524 | 0.573 | 0.795 | 0.772 | 0.782 | 0.823 | 0.762 | | | | | | |
| 0.436 | 0.642 | 0.651 | 0.587 | 0.644 | 0.624 | 0.670 | 0.705 | 0.598 | 0.748 | 0.862 | 0.629 | 0.788 | 0.014 | 0.330 | 0.054 |
| H09 | 2 | F | . | . | Lewa | 0.767 | 0.867 | 0.811 | 0.734 | 0.817 | 0.917 | 0.950 | 0.950 | 1.000 | 0.950 |
| 0.327 | 0.466 | 0.481 | 0.586 | 0.779 | 0.734 | 0.908 | 0.958 | 0.734 | 0.900 | | | | | | |
| 0.694 | 0.615 | 0.808 | 0.898 | 0.740 | 0.775 | 0.676 | 0.702 | 0.664 | 0.668 | 0.876 | 0.687 | 0.742 | 0.024 | 0.343 | 0.063 |
| H09 | 2 | F | . | . | Matata | 0.768 | 0.743 | 0.950 | 0.786 | 0.811 | 0.950 | 0.742 | 0.843 | 0.950 | 0.900 |
| 0.439 | 0.407 | 0.455 | 0.437 | 0.438 | 0.503 | 0.648 | 0.616 | 0.754 | 0.797 | | | | | | |
| 0.574 | 0.590 | 0.476 | 0.651 | 0.648 | 0.530 | 0.656 | 0.695 | 0.757 | 0.571 | 0.844 | 0.549 | 0.774 | 0.013 | 0.311 | 0.043 |
| H09 | 2 | F | . | . | Nala | 0.685 | 0.867 | 1.000 | 0.950 | 0.817 | 1.000 | 1.000 | 0.893 | 0.950 | 1.000 |
| 0.461 | 0.581 | 0.917 | 0.858 | 0.679 | 0.933 | 0.806 | 0.752 | 1.000 | 1.000 | | | | | | |
| 0.692 | 0.565 | 0.696 | 0.690 | 0.694 | 0.717 | 0.670 | 0.651 | 0.711 | 0.643 | 0.916 | 0.799 | 0.809 | 0.020 | 0.563 | 0.043 |
| H09 | 2 | F | . | . | Ollie | 0.596 | 0.702 | 0.843 | 0.893 | 0.850 | 0.889 | 0.950 | 0.850 | 0.843 | 1.000 |
| 0.448 | 0.433 | 0.536 | 0.577 | 0.535 | 0.583 | 0.698 | 0.814 | 0.634 | 0.867 | | | | | | |
| 0.606 | 0.684 | 0.564 | 0.631 | 0.626 | 0.768 | 0.678 | 0.704 | 0.655 | 0.664 | 0.841 | 0.612 | 0.679 | 0.030 | 0.380 | 0.042 |
| H09 | 2 | F | . | . | Pumba | 0.585 | 0.725 | 0.784 | 0.900 | 1.000 | 0.900 | 1.000 | 0.950 | 0.917 | 0.950 |
| 0.338 | 0.367 | 0.489 | 0.473 | 0.686 | 0.720 | 0.374 | 0.670 | 0.784 | 0.733 | | | | | | |
| 0.975 | 0.923 | 0.771 | 0.749 | 0.750 | 0.798 | 0.748 | 0.776 | 0.796 | 0.825 | 0.871 | 0.563 | 0.682 | 0.034 | 0.326 | 0.043 |
| H09 | 2 | F | . | . | Qwanda | 0.917 | 0.861 | 0.917 | 0.843 | 0.834 | 0.933 | 0.950 | 0.950 | 0.917 | 1.000 |
| 0.332 | 0.380 | 0.405 | 0.550 | 0.595 | 0.767 | 0.789 | 0.845 | 0.721 | 0.767 | | | | | | |
| 0.487 | 0.583 | 0.460 | 0.520 | 0.556 | 0.627 | 0.762 | 0.582 | 0.562 | 0.619 | 0.912 | 0.615 | 0.855 | 0.010 | 0.302 | 0.057 |
| H09 | 2 | F | . | . | Rudi | 0.784 | 0.728 | 0.950 | 0.793 | 0.950 | 0.889 | 0.917 | 0.950 | 0.867 | 0.917 |
| 0.336 | 0.505 | 0.528 | 0.640 | 0.740 | 0.604 | 0.679 | 0.867 | 0.875 | 0.964 | | | | | | |
| 0.858 | 0.934 | 0.631 | 0.826 | 0.698 | 0.863 | 0.730 | 0.751 | 0.828 | 0.838 | 0.874 | 0.674 | 0.792 | 0.015 | 0.343 | 0.060 |
| H09 | 2 | F | . | . | Siti | 0.783 | 0.685 | 0.750 | 0.810 | 0.917 | 0.841 | 0.950 | 0.810 | 0.861 | 0.817 |
| 0.346 | 0.420 | 0.460 | 0.482 | 0.570 | 0.754 | 0.629 | 0.700 | 0.785 | 0.906 | | | | | | |
| 0.662 | 0.464 | 0.543 | 0.572 | 0.617 | 0.668 | 0.612 | 0.710 | 0.660 | 0.653 | 0.822 | 0.605 | 0.750 | 0.013 | 0.291 | 0.057 |
| H09 | 2 | F | . | . | Thandi | 0.811 | 1.000 | 0.736 | 0.950 | 1.000 | 0.950 | 1.000 | 0.950 | 0.917 | 0.917 |
| 0.335 | 0.333 | 0.400 | 0.718 | 0.728 | 0.808 | 0.635 | 0.494 | 0.771 | 0.743 | | | | | | |
| 0.539 | 0.501 | 0.590 | 0.546 | 0.571 | 0.627 | 0.699 | 0.632 | 0.628 | 0.651 | 0.923 | 0.596 | 0.872 | 0.009 | 0.362 | 0.043 |
| H10 | 1 | F | . | . | 1 | 0.742 | 0.761 | 0.678 | 0.628 | 0.685 | 0.950 | 0.950 | 0.950 | 0.917 | 1.000 |
| 0.238 | 0.350 | 0.384 | 0.380 | 0.471 | 0.875 | 0.843 | 0.560 | 0.888 | 0.784 | | | | | | |
| 0.941 | 0.691 | 0.716 | 0.741 | 0.790 | 1.911 | 1.068 | 0.805 | 0.863 | 0.844 | 0.826 | 0.577 | 0.626 | 0.036 | 0.199 | 0.069 |
| H10 | 1 | F | . | . | 2 | 0.838 | 0.717 | 0.750 | 0.709 | 0.692 | 0.684 | 0.843 | 0.900 | 0.843 | 0.950 |
| 0.250 | 0.369 | 0.440 | 0.590 | 0.489 | 0.585 | 0.531 | 0.724 | 0.836 | 0.851 | | | | | | |
| 1.341 | 1.422 | 1.734 | 1.223 | 1.452 | 1.103 | 0.901 | 0.905 | 1.029 | 0.951 | 0.792 | 0.566 | 0.691 | 0.018 | 0.232 | 0.061 |
| H10 | 1 | F | . | . | 3 | 0.700 | 0.615 | 0.625 | 0.667 | 0.775 | 1.000 | 0.708 | 0.843 | 0.917 | 0.700 |
| 0.264 | 0.349 | 0.350 | 0.373 | 0.326 | 0.363 | 0.512 | 0.442 | 0.578 | 0.480 | | | | | | |
| 1.699 | 1.065 | 0.908 | 0.858 | 0.808 | 0.686 | 1.073 | 0.846 | 0.941 | 0.924 | 0.755 | 0.403 | 0.636 | 0.022 | 0.255 | 0.027 |
| H10 | 1 | F | . | . | 4 | 0.817 | 0.774 | 0.793 | 0.843 | 0.950 | 0.817 | 0.743 | 0.900 | 0.950 | 1.000 |
| 0.261 | 0.327 | 0.389 | 0.358 | 0.430 | 0.536 | 0.458 | 0.446 | 0.599 | 0.685 | | | | | | |
| 1.109 | 0.957 | 0.740 | 0.800 | 0.622 | 0.767 | 0.978 | 0.866 | 0.823 | 0.823 | 0.859 | 0.449 | 0.759 | 0.018 | 0.235 | 0.039 |
| H10 | 1 | F | . | . | 5 | 0.817 | 0.900 | 0.917 | 0.716 | 0.867 | 0.825 | 1.000 | 0.843 | 0.917 | 0.950 |
| 0.275 | 0.253 | 0.356 | 0.372 | 0.495 | 0.526 | 0.676 | 0.595 | 0.771 | 0.676 | | | | | | |
| 0.887 | 0.696 | 0.712 | 0.890 | 0.743 | 0.787 | 0.793 | 0.871 | 0.859 | 0.697 | 0.875 | 0.500 | 0.817 | 0.011 | 0.187 | 0.057 |
| H10 | 1 | F | . | . | 6 | 0.857 | 0.517 | 0.636 | 0.850 | 0.725 | 0.900 | 1.000 | 0.650 | 0.756 | 1.000 |
| 0.285 | 0.493 | 0.671 | 0.465 | 0.503 | 0.607 | 0.825 | 0.958 | 0.766 | 0.685 | | | | | | |
| 0.824 | 1.231 | 1.889 | 0.942 | 0.768 | 0.689 | 0.851 | 1.482 | 1.623 | 0.765 | 0.789 | 0.626 | 0.667 | 0.022 | 0.355 | 0.049 |
| H10 | 1 | F | . | . | 7 | 0.811 | 0.535 | 0.833 | 0.778 | 0.950 | 0.950 | 1.000 | 0.767 | 0.662 | 0.800 |
| 0.309 | 0.341 | 0.396 | 0.397 | 0.424 | 0.490 | 0.611 | 0.781 | 0.569 | 0.446 | | | | | | |
| 0.893 | 1.131 | 1.208 | 0.826 | 0.667 | 0.730 | 0.793 | 1.097 | 1.495 | 1.160 | 0.809 | 0.476 | 0.771 | 0.007 | 0.294 | 0.033 |
| H10 | 1 | F | . | . | 8 | 0.792 | 0.760 | 0.728 | 0.875 | 0.867 | 0.900 | 0.950 | 0.950 | 0.760 | 0.900 |
| 0.301 | 0.469 | 0.504 | 0.504 | 0.663 | 0.593 | 0.834 | 0.738 | 0.751 | 0.734 | | | | | | |
| 1.121 | 1.003 | 0.917 | 0.818 | 0.832 | 0.836 | 0.872 | 0.857 | 0.960 | 0.817 | 0.848 | 0.609 | 0.770 | 0.014 | 0.344 | 0.048 |
| H10 | 1 | F | . | . | 9 | 0.775 | 0.817 | 0.843 | 0.676 | 1.000 | 0.950 | 0.950 | 1.000 | 0.867 | 0.917 |
| 0.320 | 0.366 | 0.336 | 0.459 | 0.711 | 0.739 | 0.731 | 0.784 | 0.938 | 0.750 | | | | | | |
| 0.656 | 0.591 | 0.709 | 0.622 | 0.639 | 0.701 | 0.738 | 0.716 | 0.865 | 0.745 | 0.879 | 0.613 | 0.773 | 0.019 | 0.248 | 0.066 |

| | | | | | | | | | | | | | | | |
|---|---|---|---|---|---|---|---|---|---|---|---|---|---|---|---|
| H10 | 1 | F | . | . | 10 | 0.743 | 1.000 | 0.917 | 0.754 | 0.834 | 0.867 | 1.000 | 1.000 | 0.784 | 0.867 |
| 0.301 | 0.316 | 0.454 | 0.464 | 0.543 | 0.615 | 0.867 | 0.900 | 0.810 | 0.770 | | | | | | |
| 1.190 | 0.694 | 0.670 | 0.723 | 0.692 | 0.924 | 0.809 | 0.921 | 1.133 | 1.121 | 0.876 | 0.604 | 0.850 | 0.005 | 0.231 | 0.068 |
| H10 | 1 | M | . | . | 11 | 0.950 | 0.728 | 0.784 | 0.825 | 0.950 | 0.743 | 1.000 | 0.917 | 1.000 | 0.950 |
| 0.291 | 0.323 | 0.490 | 0.573 | 0.721 | 0.823 | 0.867 | 0.864 | 0.817 | 0.950 | | | | | | |
| 1.381 | 1.107 | 0.962 | 1.101 | 1.121 | 1.120 | 0.994 | 0.955 | 0.812 | 0.920 | 0.885 | 0.672 | 0.788 | 0.018 | 0.264 | 0.074 |
| H10 | 1 | M | . | . | 12 | 0.625 | 0.817 | 0.674 | 0.734 | 0.817 | 0.726 | 0.900 | 0.900 | 0.646 | 0.709 |
| 0.301 | 0.334 | 0.325 | 0.406 | 0.445 | 0.611 | 0.606 | 0.655 | 0.792 | 0.680 | | | | | | |
| 1.118 | 0.848 | 1.433 | 1.052 | 0.938 | 1.081 | 1.131 | 1.032 | 1.143 | 1.144 | 0.755 | 0.515 | 0.718 | 0.007 | 0.214 | 0.055 |
| H10 | 1 | M | . | . | 13 | 0.900 | 0.867 | 0.768 | 0.710 | 0.850 | 0.917 | 0.917 | 1.000 | 0.893 | 1.000 |
| 0.231 | 0.348 | 0.377 | 0.446 | 0.394 | 0.477 | 0.636 | 0.610 | 0.557 | 0.617 | | | | | | |
| 0.738 | 0.727 | 0.644 | 0.627 | 0.648 | 0.613 | 0.662 | 0.663 | 0.851 | 0.722 | 0.882 | 0.469 | 0.784 | 0.018 | 0.244 | 0.041 |
| H10 | 1 | M | . | . | 14 | 0.900 | 0.750 | 0.833 | 0.670 | 0.565 | 0.603 | 0.625 | 0.900 | 0.768 | 0.950 |
| 0.292 | 0.496 | 0.345 | 0.684 | 0.625 | 0.552 | 0.716 | 0.644 | 0.779 | 0.675 | | | | | | |
| 1.594 | 1.387 | 1.843 | 1.835 | 1.639 | 1.383 | 1.513 | 1.156 | 1.310 | 1.112 | 0.756 | 0.581 | 0.729 | 0.005 | 0.349 | 0.042 |
| H10 | 1 | M | . | . | 15 | 0.900 | 0.710 | 0.710 | 0.760 | 0.659 | 0.900 | 1.000 | 0.767 | 1.000 | 1.000 |
| 0.291 | 0.344 | 0.385 | 0.453 | 0.534 | 0.740 | 0.603 | 0.668 | 0.793 | 0.767 | | | | | | |
| 0.921 | 0.817 | 0.812 | 0.830 | 0.834 | 0.893 | 0.827 | 0.887 | 0.861 | 0.851 | 0.840 | 0.558 | 0.701 | 0.025 | 0.241 | 0.058 |
| H10 | 1 | M | . | . | 16 | 0.761 | 0.476 | 0.613 | 0.567 | 0.479 | 0.674 | 0.875 | 0.889 | 0.843 | 0.676 |
| 0.304 | 0.501 | 0.442 | 0.423 | 0.526 | 0.517 | 0.650 | 0.495 | 0.408 | 0.468 | | | | | | |
| 1.406 | 1.557 | 1.641 | 1.589 | 1.459 | 1.810 | 2.005 | 1.772 | 1.091 | 0.976 | 0.685 | 0.473 | 0.542 | 0.026 | 0.415 | 0.011 |
| H10 | 1 | M | . | . | 17 | 0.817 | 0.631 | 0.843 | 0.950 | 0.775 | 0.893 | 0.760 | 0.708 | 0.850 | 0.843 |
| 0.295 | 0.383 | 0.395 | 0.391 | 0.465 | 0.414 | 0.545 | 0.491 | 0.493 | 0.646 | | | | | | |
| 0.922 | 1.275 | 0.833 | 0.755 | 1.022 | 0.881 | 1.131 | 0.989 | 0.964 | 0.862 | 0.807 | 0.452 | 0.786 | 0.004 | 0.291 | 0.029 |
| H10 | 1 | M | . | . | 18 | 0.850 | 0.900 | 0.850 | 0.893 | 0.750 | 0.867 | 0.917 | 0.867 | 0.893 | 0.900 |
| 0.296 | 0.332 | 0.447 | 0.434 | 0.526 | 0.654 | 0.767 | 0.816 | 0.811 | 0.764 | | | | | | |
| 1.109 | 1.019 | 0.852 | 0.860 | 1.146 | 1.060 | 0.872 | 0.896 | 1.073 | 0.938 | 0.869 | 0.585 | 0.846 | 0.004 | 0.233 | 0.064 |
| H10 | 1 | M | . | . | 19 | 0.709 | 0.593 | 0.792 | 0.843 | 0.800 | 1.000 | 0.775 | 0.950 | 0.810 | 0.917 |
| 0.268 | 0.335 | 0.376 | 0.422 | 0.499 | 0.476 | 0.588 | 0.690 | 0.845 | 0.734 | | | | | | |
| 1.427 | 1.057 | 0.848 | 0.855 | 0.912 | 0.638 | 0.815 | 0.722 | 0.810 | 0.787 | 0.819 | 0.523 | 0.679 | 0.025 | 0.196 | 0.059 |
| H10 | 1 | M | . | . | 20 | 0.750 | 0.750 | 0.768 | 0.761 | 0.850 | 0.742 | 0.810 | 0.750 | 0.893 | 0.793 |
| 0.319 | 0.300 | 0.414 | 0.480 | 0.458 | 0.587 | 0.755 | 0.769 | 0.775 | 0.873 | | | | | | |
| 1.686 | 1.778 | 1.291 | 1.793 | 1.268 | 1.121 | 1.382 | 1.179 | 1.085 | 1.172 | 0.787 | 0.573 | 0.742 | 0.008 | 0.205 | 0.067 |
| H11 | 4 | F | LBW | . | Ariel | 0.650 | 0.557 | 0.651 | 0.861 | 0.900 | 0.917 | 1.000 | 0.825 | 0.843 | 1.000 |
| 0.257 | 0.278 | 0.363 | 0.483 | 0.562 | 0.626 | 0.702 | 0.623 | 0.714 | 0.867 | | | | | | |
| 0.949 | 1.031 | 0.730 | 0.720 | 0.730 | 0.920 | 0.754 | 0.952 | 0.869 | 0.834 | 0.820 | 0.547 | 0.605 | 0.039 | 0.195 | 0.064 |
| H11 | 4 | F | LBW | . | Babe | 0.768 | 0.784 | 0.700 | 0.726 | 1.000 | 0.900 | 0.950 | 0.950 | 0.950 | 0.950 |
| 0.313 | 0.346 | 0.381 | 0.404 | 0.456 | 0.580 | 0.612 | 0.639 | 0.716 | 0.795 | | | | | | |
| 0.609 | 0.600 | 0.659 | 0.647 | 0.768 | 0.651 | 0.629 | 0.714 | 0.669 | 0.722 | 0.868 | 0.524 | 0.714 | 0.028 | 0.225 | 0.054 |
| H11 | 4 | F | LBW | . | Bentley | 0.950 | 0.719 | 0.768 | 0.760 | 0.754 | 1.000 | 0.685 | 0.843 | 0.861 | 0.900 |
| 0.356 | 0.288 | 0.357 | 0.370 | 0.586 | 0.767 | 0.527 | 0.854 | 0.813 | 0.787 | | | | | | |
| 0.524 | 0.805 | 0.544 | 0.688 | 0.835 | 0.746 | 1.012 | 0.996 | 0.892 | 1.107 | 0.824 | 0.570 | 0.792 | 0.006 | 0.214 | 0.065 |
| H11 | 4 | F | LBW | . | Cookie | 0.710 | 0.760 | 0.686 | 0.900 | 0.861 | 1.000 | 0.917 | 1.000 | 0.900 | 0.950 |
| 0.312 | 0.298 | 0.343 | 0.299 | 0.390 | 0.382 | 0.541 | 0.528 | 0.646 | 0.593 | | | | | | |
| 0.623 | 0.640 | 0.565 | 0.519 | 0.577 | 0.601 | 0.618 | 0.588 | 0.640 | 0.630 | 0.868 | 0.433 | 0.705 | 0.030 | 0.213 | 0.040 |
| H11 | 4 | M | LBW | . | Cooper | 0.624 | 0.586 | 0.742 | 0.867 | 0.867 | 0.843 | 0.950 | 1.000 | 0.950 | 1.000 |
| 0.304 | 0.347 | 0.296 | 0.402 | 0.496 | 0.516 | 0.592 | 0.611 | 0.825 | 0.950 | | | | | | |
| 1.406 | 1.075 | 0.684 | 0.562 | 0.640 | 0.731 | 0.711 | 0.692 | 0.804 | 0.882 | 0.843 | 0.534 | 0.594 | 0.045 | 0.156 | 0.069 |
| H11 | 4 | M | LBW | . | Delta | 0.817 | 0.817 | 0.535 | 0.726 | 1.000 | 0.917 | 0.825 | 0.867 | 0.867 | 0.950 |
| 0.340 | 0.326 | 0.269 | 0.363 | 0.419 | 0.584 | 0.534 | 0.723 | 0.550 | 0.742 | | | | | | |
| 0.634 | 0.646 | 0.702 | 0.621 | 0.619 | 0.654 | 0.664 | 0.699 | 0.767 | 0.866 | 0.832 | 0.485 | 0.718 | 0.021 | 0.214 | 0.049 |
| H11 | 4 | M | LBW | . | Echo | 0.825 | 0.893 | 0.633 | 0.917 | 0.817 | 0.850 | 0.843 | 0.893 | 1.000 | 1.000 |
| 0.308 | 0.302 | 0.401 | 0.584 | 0.539 | 0.839 | 0.831 | 0.875 | 0.900 | 0.950 | | | | | | |
| 1.038 | 0.765 | 0.879 | 0.700 | 0.731 | 1.062 | 0.924 | 0.926 | 0.914 | 0.901 | 0.867 | 0.653 | 0.752 | 0.021 | 0.207 | 0.081 |
| H11 | 4 | M | LBW | . | Elliott | 0.850 | 0.811 | 0.900 | 0.950 | 0.950 | 1.000 | 0.917 | 0.850 | 0.834 | 0.893 |
| 0.341 | 0.339 | 0.479 | 0.557 | 0.574 | 0.587 | 0.622 | 0.657 | 0.676 | 0.734 | | | | | | |
| 0.704 | 0.798 | 0.626 | 0.720 | 0.717 | 0.806 | 0.732 | 0.799 | 0.718 | 0.751 | 0.895 | 0.556 | 0.887 | 0.001 | 0.323 | 0.042 |
| H11 | 4 | F | NBW | . | Ernie | 0.786 | 0.850 | 0.827 | 0.817 | 0.867 | 0.950 | 0.917 | 1.000 | 1.000 | 1.000 |
| 0.300 | 0.433 | 0.441 | 0.637 | 0.950 | 0.851 | 0.864 | 0.850 | 0.917 | 0.950 | | | | | | |
| 0.718 | 0.635 | 0.780 | 0.817 | 0.851 | 0.741 | 0.766 | 0.789 | 0.765 | 0.757 | 0.901 | 0.719 | 0.760 | 0.026 | 0.324 | 0.072 |
| H11 | 4 | F | LBW | . | Fien | 0.893 | 1.000 | 0.792 | 0.950 | 0.950 | 0.950 | 1.000 | 1.000 | 0.917 | 1.000 |
| 0.336 | 0.298 | 0.452 | 0.442 | 0.490 | 0.507 | 0.579 | 0.668 | 0.635 | 0.843 | | | | | | |
| 0.602 | 0.621 | 0.692 | 0.681 | 0.682 | 0.643 | 0.701 | 0.736 | 0.771 | 0.832 | 0.945 | 0.525 | 0.893 | 0.010 | 0.244 | 0.051 |
| H11 | 4 | M | LBW | . | Forrest | 0.850 | 0.691 | 0.917 | 0.850 | 0.917 | 1.000 | 0.843 | 0.893 | 1.000 | 1.000 |
| 0.339 | 0.363 | 0.310 | 0.439 | 0.416 | 0.476 | 0.540 | 0.474 | 0.661 | 0.494 | | | | | | |
| 0.605 | 1.031 | 0.826 | 0.668 | 0.564 | 0.576 | 0.637 | 0.601 | 0.605 | 0.485 | 0.896 | 0.451 | 0.781 | 0.021 | 0.296 | 0.028 |
| H11 | 4 | M | NBW | . | Frodo | 0.867 | 0.861 | 0.793 | 0.817 | 0.950 | 1.000 | 0.950 | 0.950 | 1.000 | 1.000 |
| 0.254 | 0.336 | 0.389 | 0.401 | 0.393 | 0.466 | 0.573 | 0.700 | 0.850 | 0.704 | | | | | | |
| 0.480 | 0.519 | 0.451 | 0.444 | 0.493 | 0.585 | 0.649 | 0.618 | 0.712 | 0.647 | 0.919 | 0.506 | 0.805 | 0.021 | 0.180 | 0.059 |
| H11 | 4 | M | LBW | . | Glenn | 0.424 | 0.479 | 0.775 | 0.695 | 0.950 | 0.793 | 1.000 | 0.792 | 0.778 | 1.000 |
| 0.357 | 0.338 | 0.402 | 0.484 | 0.707 | 0.614 | 0.750 | 0.701 | 0.649 | 0.760 | | | | | | |
| 1.724 | 0.954 | 0.687 | 0.848 | 0.689 | 0.657 | 0.773 | 0.820 | 0.795 | 0.819 | 0.769 | 0.576 | 0.498 | 0.049 | 0.309 | 0.049 |
| H11 | 4 | M | LBW | . | Hudson | 0.817 | 0.867 | 0.867 | 0.950 | 0.850 | 0.950 | 0.950 | 0.950 | 0.950 | 0.917 |
| 0.328 | 0.293 | 0.454 | 0.773 | 0.729 | 0.829 | 0.775 | 0.908 | 0.875 | 0.730 | | | | | | |
| 0.895 | 0.820 | 0.763 | 0.815 | 0.782 | 0.821 | 0.905 | 0.909 | 0.854 | 0.847 | 0.907 | 0.669 | 0.840 | 0.012 | 0.334 | 0.061 |
| H11 | 4 | F | LBW | . | Indy | 0.800 | 0.619 | 0.736 | 0.893 | 0.900 | 0.850 | 0.917 | 0.738 | 0.917 | 1.000 |
| 0.288 | 0.363 | 0.338 | 0.341 | 0.497 | 0.493 | 0.775 | 0.646 | 0.601 | 0.867 | | | | | | |
| 1.110 | 0.716 | 0.623 | 0.640 | 0.595 | 0.683 | 0.749 | 0.842 | 0.706 | 0.843 | 0.837 | 0.521 | 0.706 | 0.024 | 0.197 | 0.059 |
| H11 | 4 | M | LBW | . | James | 0.736 | 0.734 | 0.654 | 0.817 | 0.559 | 0.825 | 0.700 | 0.676 | 0.867 | 0.700 |

| | | | | | | | | | | | | | | | |
|---|---|---|---|---|---|---|---|---|---|---|---|---|---|---|---|
| 0.355 | 0.326 | 0.365 | 0.422 | 0.381 | 0.495 | 0.470 | 0.652 | 0.615 | 0.638 | | | | | | |
| 0.769 | 0.659 | 0.574 | 0.608 | 0.564 | 0.623 | 0.681 | 0.741 | 0.884 | 0.759 | 0.727 | 0.472 | 0.705 | 0.004 | 0.263 | 0.038 |
| H11 | 4 | M | NBW | . | Kaz | 0.694 | 0.900 | 0.784 | 0.900 | 0.950 | 1.000 | 0.893 | 0.825 | 0.817 | 0.950 |
| 0.328 | 0.331 | 0.411 | 0.549 | 0.675 | 0.867 | 0.842 | 0.663 | 0.839 | 1.000 | | | | | | |
| 0.821 | 0.726 | 0.666 | 0.679 | 0.798 | 0.888 | 0.873 | 1.079 | 0.963 | 0.913 | 0.871 | 0.650 | 0.806 | 0.012 | 0.252 | 0.072 |
| H11 | 4 | F | NBW | . | Lara | 0.687 | 0.718 | 0.767 | 0.793 | 0.783 | 0.867 | 0.950 | 0.950 | 0.700 | 0.900 |
| 0.290 | 0.379 | 0.386 | 0.509 | 0.481 | 0.556 | 0.575 | 0.821 | 0.771 | 0.781 | | | | | | |
| 1.006 | 0.841 | 0.599 | 0.661 | 0.742 | 0.705 | 0.759 | 0.741 | 0.808 | 0.949 | 0.811 | 0.555 | 0.702 | 0.020 | 0.234 | 0.058 |
| H11 | 4 | M | NBW | . | Lux | 0.950 | 0.867 | 0.778 | 1.000 | 1.000 | 0.850 | 0.867 | 0.900 | 0.950 | 1.000 |
| 0.319 | 0.343 | 0.361 | 0.389 | 0.470 | 0.452 | 0.592 | 0.497 | 0.601 | 0.512 | | | | | | |
| 0.535 | 0.553 | 0.675 | 0.535 | 0.620 | 0.719 | 0.619 | 0.578 | 0.661 | 0.589 | 0.916 | 0.453 | 0.880 | 0.007 | 0.293 | 0.029 |
| H11 | 4 | F | NBW | . | May | 0.678 | 0.726 | 0.810 | 0.843 | 0.841 | 0.843 | 0.850 | 0.917 | 0.917 | 1.000 |
| 0.304 | 0.304 | 0.335 | 0.333 | 0.449 | 0.680 | 0.714 | 0.867 | 0.767 | 0.917 | | | | | | |
| 0.681 | 0.694 | 0.602 | 0.638 | 0.689 | 0.797 | 0.827 | 0.838 | 0.746 | 0.790 | 0.842 | 0.567 | 0.683 | 0.029 | 0.140 | 0.078 |
| H11 | 4 | F | NBW | . | Molly | 0.742 | 0.678 | 0.917 | 1.000 | 0.775 | 0.778 | 1.000 | 0.950 | 0.917 | 1.000 |
| 0.253 | 0.363 | 0.414 | 0.413 | 0.539 | 0.592 | 0.643 | 0.718 | 0.673 | 0.659 | | | | | | |
| 0.908 | 0.839 | 0.666 | 0.652 | 0.713 | 0.680 | 0.720 | 0.738 | 0.808 | 0.697 | 0.876 | 0.527 | 0.737 | 0.025 | 0.257 | 0.049 |
| H11 | 4 | F | LBW | . | Muis | 0.867 | 0.950 | 0.817 | 0.811 | 0.867 | 0.867 | 1.000 | 0.950 | 0.917 | 0.800 |
| 0.299 | 0.343 | 0.440 | 0.473 | 0.760 | 0.733 | 1.000 | 0.908 | 0.807 | 0.734 | | | | | | |
| 0.672 | 0.753 | 0.748 | 0.725 | 0.835 | 0.775 | 0.829 | 0.784 | 0.803 | 0.708 | 0.884 | 0.650 | 0.871 | 0.002 | 0.281 | 0.067 |
| H11 | 4 | F | NBW | . | Nola | 0.811 | 0.676 | 0.843 | 0.875 | 1.000 | 0.850 | 0.900 | 1.000 | 1.000 | 0.875 |
| 0.240 | 0.376 | 0.437 | 0.401 | 0.468 | 0.599 | 0.637 | 0.900 | 1.000 | 0.882 | | | | | | |
| 0.510 | 0.551 | 0.579 | 0.555 | 0.553 | 0.637 | 0.690 | 0.738 | 0.769 | 0.774 | 0.883 | 0.594 | 0.765 | 0.022 | 0.151 | 0.081 |
| H11 | 4 | F | NBW | . | Nukka | 0.784 | 0.793 | 0.702 | 0.717 | 0.950 | 0.900 | 0.810 | 0.950 | 1.000 | 0.900 |
| 0.331 | 0.329 | 0.457 | 0.478 | 0.685 | 0.698 | 0.676 | 0.950 | 0.659 | 0.809 | | | | | | |
| 0.746 | 0.599 | 0.531 | 0.621 | 0.672 | 0.733 | 0.744 | 0.795 | 0.791 | 0.854 | 0.850 | 0.607 | 0.718 | 0.024 | 0.284 | 0.059 |
| H11 | 4 | F | NBW | . | Odette | 0.950 | 0.726 | 0.767 | 0.850 | 0.917 | 0.867 | 0.817 | 0.950 | 1.000 | 1.000 |
| 0.280 | 0.324 | 0.407 | 0.474 | 0.602 | 0.711 | 0.663 | 0.834 | 0.850 | 0.950 | | | | | | |
| 0.757 | 0.800 | 0.714 | 0.783 | 0.756 | 1.078 | 0.823 | 0.884 | 0.911 | 0.872 | 0.884 | 0.609 | 0.780 | 0.019 | 0.192 | 0.076 |
| H11 | 4 | M | LBW | . | Oliver | 1.000 | 0.784 | 0.604 | 0.810 | 0.843 | 0.875 | 1.000 | 0.760 | 0.950 | 0.917 |
| 0.317 | 0.317 | 0.456 | 0.489 | 0.696 | 0.554 | 0.652 | 0.569 | 0.664 | 0.786 | | | | | | |
| 0.682 | 0.739 | 0.732 | 0.694 | 0.784 | 0.756 | 0.722 | 0.657 | 0.680 | 0.747 | 0.854 | 0.550 | 0.794 | 0.011 | 0.298 | 0.046 |
| H11 | 4 | F | LBW | . | Peach | 0.917 | 0.817 | 0.784 | 0.817 | 0.658 | 0.709 | 0.768 | 0.867 | 0.817 | 1.000 |
| 0.261 | 0.322 | 0.332 | 0.405 | 0.506 | 0.595 | 0.546 | 0.727 | 0.749 | 0.793 | | | | | | |
| 0.644 | 0.619 | 0.616 | 0.582 | 0.725 | 0.638 | 0.686 | 0.662 | 0.663 | 0.746 | 0.815 | 0.523 | 0.780 | 0.006 | 0.181 | 0.062 |
| H11 | 4 | M | NBW | . | Pino | 0.893 | 0.825 | 0.767 | 1.000 | 1.000 | 0.917 | 1.000 | 0.917 | 0.917 | 0.843 |
| 0.344 | 0.324 | 0.414 | 0.521 | 0.726 | 0.710 | 0.950 | 0.743 | 0.834 | 0.867 | | | | | | |
| 0.610 | 0.589 | 0.639 | 0.649 | 0.779 | 0.814 | 0.908 | 0.865 | 0.894 | 0.934 | 0.908 | 0.643 | 0.879 | 0.005 | 0.270 | 0.068 |
| H11 | 4 | M | NBW | . | Pumbaa | 0.900 | 0.709 | 0.600 | 0.950 | 0.950 | 0.813 | 0.950 | 0.652 | 0.784 | 0.725 |
| 0.308 | 0.369 | 0.341 | 0.373 | 0.387 | 0.558 | 0.533 | 0.503 | 0.514 | 0.569 | | | | | | |
| 0.676 | 0.690 | 0.929 | 0.749 | 0.688 | 0.895 | 0.713 | 1.034 | 0.903 | 0.875 | 0.803 | 0.445 | 0.834 | -0.006 | 0.284 | 0.029 |
| H11 | 4 | M | NBW | . | Quat | 0.760 | 0.760 | 0.825 | 0.900 | 0.817 | 0.810 | 0.950 | 0.950 | 1.000 | 1.000 |
| 0.314 | 0.331 | 0.483 | 0.538 | 0.560 | 0.640 | 0.674 | 0.893 | 1.000 | 1.000 | | | | | | |
| 0.689 | 0.692 | 0.643 | 0.612 | 0.697 | 0.750 | 0.691 | 0.912 | 0.899 | 0.847 | 0.877 | 0.643 | 0.723 | 0.028 | 0.197 | 0.081 |
| H11 | 4 | M | NBW | . | Rudolph | 0.810 | 0.817 | 0.784 | 0.950 | 0.893 | 0.950 | 0.900 | 0.950 | 0.867 | 0.950 |
| 0.299 | 0.316 | 0.460 | 0.461 | 0.594 | 0.657 | 0.724 | 0.649 | 0.767 | 0.825 | | | | | | |
| 0.654 | 0.624 | 0.714 | 0.660 | 0.762 | 0.775 | 0.752 | 0.791 | 0.923 | 0.868 | 0.887 | 0.575 | 0.808 | 0.014 | 0.252 | 0.059 |
| H11 | 4 | F | NBW | . | Sem | 0.767 | 0.900 | 0.833 | 0.900 | 0.867 | 0.817 | 1.000 | 0.900 | 0.900 | 0.825 |
| 0.296 | 0.343 | 0.382 | 0.487 | 0.402 | 0.613 | 0.603 | 0.713 | 0.718 | 0.656 | | | | | | |
| 0.694 | 0.738 | 0.822 | 0.781 | 0.730 | 0.813 | 0.829 | 1.040 | 0.803 | 0.949 | 0.871 | 0.521 | 0.834 | 0.007 | 0.252 | 0.049 |
| H11 | 4 | F | NBW | . | Smeagel | 0.768 | 0.611 | 0.760 | 0.818 | 0.843 | 0.834 | 0.817 | 0.792 | 0.768 | 0.950 |
| 0.303 | 0.317 | 0.436 | 0.481 | 0.520 | 0.534 | 0.793 | 0.657 | 0.786 | 0.879 | | | | | | |
| 0.663 | 0.622 | 0.551 | 0.835 | 0.609 | 0.607 | 0.626 | 0.668 | 0.749 | 0.675 | 0.796 | 0.570 | 0.700 | 0.017 | 0.220 | 0.064 |
| H11 | 4 | M | LBW | . | Snorro | 0.867 | 0.768 | 0.900 | 0.950 | 0.810 | 1.000 | 0.686 | 0.917 | 1.000 | 0.778 |
| 0.247 | 0.308 | 0.358 | 0.372 | 0.520 | 0.609 | 0.509 | 0.888 | 0.893 | 0.864 | | | | | | |
| 0.733 | 0.690 | 0.564 | 0.594 | 0.594 | 0.595 | 0.814 | 0.769 | 0.735 | 0.931 | 0.867 | 0.557 | 0.857 | 0.002 | 0.130 | 0.078 |
| H11 | 4 | F | NBW | . | Spot | 0.611 | 0.760 | 0.861 | 0.900 | 0.893 | 0.658 | 1.000 | 0.950 | 0.900 | 1.000 |
| 0.286 | 0.343 | 0.441 | 0.416 | 0.590 | 0.540 | 0.601 | 0.694 | 0.764 | 0.834 | | | | | | |
| 0.985 | 0.767 | 0.608 | 0.605 | 0.622 | 0.860 | 0.718 | 0.759 | 0.769 | 0.869 | 0.853 | 0.551 | 0.687 | 0.030 | 0.229 | 0.059 |
| H11 | 4 | M | NBW | . | Tank | 0.843 | 0.545 | 0.655 | 0.843 | 0.950 | 0.850 | 1.000 | 0.950 | 0.800 | 0.950 |
| 0.419 | 0.294 | 0.329 | 0.409 | 0.421 | 0.684 | 0.867 | 0.784 | 0.602 | 0.771 | | | | | | |
| 0.697 | 0.808 | 0.666 | 0.581 | 0.580 | 0.663 | 0.734 | 0.745 | 0.879 | 0.799 | 0.839 | 0.558 | 0.685 | 0.028 | 0.250 | 0.056 |
| H11 | 4 | M | LBW | . | Whiskey | 0.643 | 1.000 | 0.784 | 0.843 | 0.950 | 0.893 | 0.810 | 0.867 | 0.867 | 0.867 |
| 0.268 | 0.390 | 0.305 | 0.344 | 0.404 | 0.419 | 0.425 | 0.633 | 0.546 | 0.756 | | | | | | |
| 1.047 | 0.605 | 0.614 | 0.549 | 0.645 | 0.609 | 0.603 | 0.620 | 0.630 | 0.663 | 0.852 | 0.449 | 0.807 | 0.008 | 0.203 | 0.045 |
| H11 | 4 | M | NBW | . | Wyatt | 0.817 | 0.810 | 0.793 | 0.917 | 0.950 | 0.750 | 0.867 | 1.000 | 0.950 | 0.950 |
| 0.361 | 0.330 | 0.397 | 0.434 | 0.480 | 0.554 | 0.521 | 0.593 | 0.779 | 0.771 | | | | | | |
| 0.445 | 0.491 | 0.511 | 0.546 | 0.524 | 0.617 | 0.576 | 0.541 | 0.638 | 0.558 | 0.880 | 0.522 | 0.785 | 0.017 | 0.251 | 0.049 |
| H12 | 1 | F | . | First | Appa | 0.834 | 0.917 | 0.893 | 0.760 | 1.000 | 0.834 | 0.950 | 1.000 | 1.000 | 0.950 |
| 0.321 | 0.376 | 0.366 | 0.459 | 0.486 | 0.486 | 0.808 | 0.672 | 0.800 | 0.845 | | | | | | |
| 1.175 | 1.456 | 0.828 | 1.167 | 0.837 | 0.878 | 0.995 | 0.823 | 0.904 | 0.778 | 0.914 | 0.562 | 0.828 | 0.016 | 0.220 | 0.062 |
| H12 | 1 | M | . | Last | Babe | 0.900 | 0.768 | 1.000 | 0.850 | 0.724 | 0.900 | 0.843 | 1.000 | 0.900 | 1.000 |
| 0.298 | 0.345 | 0.338 | 0.492 | 0.509 | 0.538 | 0.664 | 0.517 | 0.695 | 0.843 | | | | | | |
| 0.853 | 0.954 | 0.703 | 0.824 | 1.165 | 0.816 | 0.967 | 1.004 | 1.000 | 0.842 | 0.888 | 0.524 | 0.822 | 0.012 | 0.231 | 0.053 |
| H12 | 1 | M | . | First | Calvin | 0.950 | 0.950 | 1.000 | 0.917 | 1.000 | 1.000 | 1.000 | 0.950 | 1.000 | 1.000 |
| 0.389 | 0.364 | 0.397 | 0.528 | 0.692 | 0.767 | 0.867 | 1.000 | 0.950 | 0.850 | | | | | | |
| 0.737 | 1.047 | 0.838 | 0.834 | 0.848 | 0.769 | 0.854 | 0.798 | 0.809 | 0.678 | 0.977 | 0.680 | 0.950 | 0.005 | 0.268 | 0.075 |
| H12 | 1 | M | . | Last | Donny | 0.825 | 0.917 | 0.950 | 0.817 | 1.000 | 1.000 | 0.950 | 1.000 | 1.000 | 0.893 |
| 0.320 | 0.302 | 0.376 | 0.335 | 0.383 | 0.484 | 0.673 | 0.559 | 0.810 | 0.785 | | | | | | |

| | | | | | | | | | | | | | | | |
|---|---|---|---|---|---|---|---|---|---|---|---|---|---|---|---|
| 1.040 | 0.884 | 0.655 | 0.862 | 1.034 | 1.009 | 0.963 | 0.758 | 0.801 | 0.806 | 0.935 | 0.502 | 0.874 | 0.011 | 0.177 | 0.059 |
| H12 | 1 | M | . | First | Fleetw. | 1.000 | 1.000 | 1.000 | 1.000 | 0.900 | 1.000 | 0.825 | 1.000 | 1.000 | 0.917 |
| 0.254 | 0.279 | 0.413 | 0.493 | 0.621 | 0.546 | 0.665 | 0.750 | 0.917 | 0.867 | | | | | | |
| 1.245 | 0.958 | 0.623 | 0.753 | 0.913 | 0.678 | 1.101 | 0.946 | 0.856 | 0.775 | 0.964 | 0.580 | 1.003 | -0.007 | 0.177 | 0.073 |
| H12 | 1 | M | . | Last | Hobbes | 0.850 | 0.888 | 0.917 | 0.900 | 0.950 | 1.000 | 0.950 | 1.000 | 1.000 | 0.950 |
| 0.290 | 0.307 | 0.466 | 0.647 | 0.726 | 0.692 | 0.829 | 0.867 | 0.834 | 1.000 | | | | | | |
| 1.071 | 1.164 | 1.060 | 0.984 | 1.058 | 0.952 | 0.995 | 0.784 | 0.878 | 0.855 | 0.940 | 0.666 | 0.864 | 0.014 | 0.246 | 0.076 |
| H12 | 1 | M | . | Last | JD | 1.000 | 0.900 | 0.950 | 0.950 | 0.867 | 1.000 | 1.000 | 1.000 | 1.000 | 0.893 |
| 0.346 | 0.338 | 0.455 | 0.397 | 0.673 | 0.716 | 0.718 | 0.661 | 0.767 | 0.909 | | | | | | |
| 0.736 | 0.875 | 0.621 | 0.719 | 0.760 | 0.725 | 0.750 | 0.669 | 0.736 | 0.939 | 0.956 | 0.598 | 0.947 | 0.002 | 0.261 | 0.061 |
| H12 | 1 | F | . | Last | Kokkie | 1.000 | 1.000 | 0.784 | 0.850 | 0.950 | 1.000 | 0.950 | 1.000 | 0.950 | 0.950 |
| 0.436 | 0.366 | 0.431 | 0.538 | 0.637 | 0.635 | 0.900 | 0.579 | 0.850 | 0.900 | | | | | | |
| 1.107 | 1.031 | 1.027 | 1.241 | 1.333 | 1.228 | 1.290 | 1.052 | 1.427 | 1.172 | 0.943 | 0.627 | 0.922 | 0.004 | 0.314 | 0.057 |
| H12 | 1 | F | . | Last | Loki | 0.917 | 1.000 | 0.950 | 0.917 | 0.900 | 1.000 | 0.950 | 1.000 | 1.000 | 0.950 |
| 0.260 | 0.376 | 0.388 | 0.364 | 0.464 | 0.547 | 0.623 | 0.676 | 0.661 | 0.839 | | | | | | |
| 1.036 | 0.687 | 0.710 | 0.633 | 0.715 | 0.725 | 0.824 | 0.660 | 0.658 | 0.632 | 0.958 | 0.520 | 0.933 | 0.005 | 0.203 | 0.058 |
| H12 | 1 | F | . | Last | Mac | 0.780 | 0.875 | 0.830 | 0.917 | 0.674 | 0.817 | 0.867 | 0.950 | 1.000 | 1.000 |
| 0.303 | 0.381 | 0.309 | 0.358 | 0.361 | 0.478 | 0.553 | 0.557 | 0.784 | 0.685 | | | | | | |
| 1.154 | 0.959 | 1.065 | 1.067 | 1.179 | 0.879 | 0.778 | 0.758 | 0.764 | 0.744 | 0.871 | 0.477 | 0.756 | 0.021 | 0.204 | 0.050 |
| H12 | 1 | F | . | First | Mary | 0.850 | 0.838 | 0.867 | 0.888 | 1.000 | 1.000 | 1.000 | 0.950 | 1.000 | 1.000 |
| 0.378 | 0.254 | 0.441 | 0.612 | 0.592 | 0.676 | 0.867 | 0.823 | 1.000 | 0.867 | | | | | | |
| 0.933 | 1.495 | 1.240 | 1.137 | 1.734 | 0.841 | 1.000 | 0.801 | 0.890 | 0.760 | 0.939 | 0.651 | 0.831 | 0.020 | 0.239 | 0.075 |
| H12 | 1 | F | . | Last | Momo | 0.917 | 0.893 | 0.950 | 0.950 | 1.000 | 0.950 | 0.917 | 1.000 | 1.000 | 1.000 |
| 0.362 | 0.355 | 0.392 | 0.522 | 0.700 | 0.806 | 0.679 | 0.843 | 0.900 | 0.950 | | | | | | |
| 1.107 | 0.890 | 0.634 | 0.856 | 0.870 | 0.901 | 0.944 | 0.770 | 0.942 | 0.795 | 0.958 | 0.651 | 0.904 | 0.010 | 0.253 | 0.072 |
| H12 | 1 | F | . | First | Peppie | 0.950 | 0.950 | 0.867 | 0.900 | 1.000 | 1.000 | 1.000 | 0.950 | 1.000 | 0.950 |
| 0.255 | 0.389 | 0.487 | 0.464 | 0.700 | 0.692 | 0.867 | 0.718 | 0.834 | 0.790 | | | | | | |
| 1.231 | 0.761 | 1.053 | 0.839 | 0.914 | 0.859 | 1.639 | 0.793 | 0.932 | 1.236 | 0.957 | 0.619 | 0.921 | 0.006 | 0.277 | 0.062 |
| H12 | 1 | M | . | Last | Pippin | 0.950 | 0.950 | 0.867 | 0.950 | 1.000 | 0.917 | 0.800 | 0.950 | 0.850 | 1.000 |
| 0.296 | 0.279 | 0.421 | 0.431 | 0.601 | 0.602 | 0.773 | 0.749 | 0.815 | 0.900 | | | | | | |
| 0.890 | 1.026 | 0.799 | 0.978 | 1.081 | 0.947 | 1.404 | 0.977 | 1.088 | 0.851 | 0.923 | 0.587 | 0.936 | -0.002 | 0.192 | 0.072 |
| H12 | 1 | M | . | First | Prima | 0.700 | 0.850 | 0.817 | 0.950 | 0.867 | 0.867 | 1.000 | 1.000 | 1.000 | 1.000 |
| 0.357 | 0.357 | 0.367 | 0.510 | 0.636 | 0.616 | 0.619 | 0.668 | 0.817 | 0.900 | | | | | | |
| 1.172 | 1.059 | 0.834 | 0.830 | 0.862 | 0.814 | 0.758 | 0.762 | 0.844 | 0.831 | 0.905 | 0.585 | 0.744 | 0.029 | 0.254 | 0.060 |
| H12 | 1 | M | . | Last | Pumba | 0.825 | 1.000 | 0.744 | 0.950 | 0.917 | 0.900 | 0.950 | 1.000 | 1.000 | 1.000 |
| 0.303 | 0.343 | 0.433 | 0.486 | 0.580 | 0.637 | 0.681 | 0.784 | 0.850 | 0.867 | | | | | | |
| 1.190 | 0.853 | 1.030 | 0.727 | 0.974 | 0.965 | 0.992 | 0.788 | 1.067 | 0.798 | 0.929 | 0.596 | 0.834 | 0.017 | 0.229 | 0.067 |
| H12 | 1 | F | . | First | Spikkel | 1.000 | 0.950 | 1.000 | 0.900 | 0.875 | 1.000 | 0.854 | 0.917 | 0.888 | 1.000 |
| 0.294 | 0.389 | 0.340 | 0.403 | 0.461 | 0.817 | 0.601 | 0.556 | 0.867 | 0.950 | | | | | | |
| 0.996 | 1.285 | 0.836 | 0.770 | 1.232 | 0.873 | 1.390 | 0.795 | 1.428 | 0.786 | 0.938 | 0.568 | 0.967 | -0.005 | 0.192 | 0.068 |
| H12 | 1 | F | . | First | Stevie | 0.667 | 0.813 | 0.825 | 0.750 | 0.950 | 0.950 | 1.000 | 0.893 | 0.867 | 0.950 |
| 0.335 | 0.281 | 0.388 | 0.577 | 0.712 | 0.615 | 0.867 | 0.639 | 0.858 | 0.823 | | | | | | |
| 1.402 | 1.421 | 1.310 | 1.464 | 1.348 | 0.992 | 1.068 | 1.016 | 1.142 | 0.898 | 0.866 | 0.609 | 0.732 | 0.024 | 0.261 | 0.063 |
| H12 | 1 | F | . | First | Thor | 0.917 | 0.900 | 0.850 | 0.850 | 0.938 | 0.917 | 0.950 | 1.000 | 1.000 | 0.950 |
| 0.291 | 0.314 | 0.437 | 0.505 | 0.479 | 0.696 | 0.685 | 0.768 | 0.736 | 0.950 | | | | | | |
| 1.037 | 0.862 | 1.204 | 0.987 | 1.268 | 0.860 | 0.902 | 0.833 | 0.829 | 0.758 | 0.927 | 0.586 | 0.859 | 0.012 | 0.209 | 0.068 |
| H12 | 1 | M | . | First | Timon | 0.900 | 0.950 | 0.917 | 1.000 | 1.000 | 0.950 | 0.867 | 0.950 | 1.000 | 1.000 |
| 0.281 | 0.265 | 0.407 | 0.470 | 0.504 | 0.564 | 0.600 | 0.542 | 0.734 | 0.867 | | | | | | |
| 0.853 | 0.887 | 0.793 | 0.703 | 0.656 | 0.685 | 0.886 | 0.733 | 0.889 | 0.763 | 0.953 | 0.523 | 0.921 | 0.006 | 0.201 | 0.059 |
| H12 | 1 | M | . | First | Turk | 0.817 | 0.950 | 1.000 | 1.000 | 0.867 | 0.917 | 0.950 | 0.950 | 0.950 | 1.000 |
| 0.255 | 0.328 | 0.393 | 0.479 | 0.544 | 0.610 | 0.698 | 0.740 | 0.845 | 0.775 | | | | | | |
| 0.999 | 0.805 | 0.747 | 0.675 | 0.876 | 0.778 | 0.751 | 0.665 | 0.874 | 0.695 | 0.940 | 0.567 | 0.897 | 0.008 | 0.208 | 0.065 |
| H12 | 1 | M | . | Last | Wonder | 0.888 | 0.950 | 0.900 | 0.843 | 0.893 | 1.000 | 0.950 | 0.950 | 0.950 | 0.950 |
| 0.275 | 0.384 | 0.328 | 0.407 | 0.478 | 0.546 | 0.768 | 0.560 | 0.929 | 0.756 | | | | | | |
| 1.197 | 1.017 | 0.865 | 1.115 | 1.030 | 0.980 | 1.167 | 0.943 | 0.978 | 0.899 | 0.927 | 0.543 | 0.886 | 0.008 | 0.195 | 0.063 |

### Information on the publications of the studies


**H01:** Gieling, E.T., Park, S.Y., Nordquist, R.E., van der Staay, F.J., 2012.
Cognitive performance of low- and normal-birth-weight piglets in a spatial hole-board discrimination task. Pediatric research 71, 71–76. https://doi.org/10.1038/pr.2011.5

**H02:** Gieling, E.T., Wehkamp, W., Willigenburg, R., Nordquist, R.E., Ganderup, N.-C., van der Staay, F.J., 2013.
Performance of conventional pigs and Göttingen miniature pigs in a spatial holeboard task: effects of the putative muscarinic cognition impairer biperiden. Behavioral and brain functions 9, 4–4. https://doi.org/10.1186/1744-9081-9-4

**H03:** Gieling, E.T., Antonides, S., Fink-Gremmels, J., ter Haar, K., Kuller, W.I., Meijer, E., Nordquist, R.E., Stouten, J.M., Zeinstra, E., van der Staay, F.J., 2014.
Chronic allopurinol treatment during the last trimester of pregnancy in sows: effects on low and normal birth weight offspring. PLoS ONE 9, e86396. https://doi.org/10.1371/journal.pone.0086396

**H04:** Antonides, A., Schoonderwoerd, A.C., Scholz, G., Berg, B.M., Nordquist, R.E., van der Staay, F.J., 2015b.
Pre-weaning dietary iron deficiency impairs spatial learning and memory in the cognitive holeboard task in piglets. Frontiers in behavioral neuroscience 9, 16. https://doi.org/10.3389/fnbeh.2015.00291

**H05:** Antonides, A., Schoonderwoerd, A.C., Nordquist, R.E., van der Staay, F.J., 2015a.
Very low birth weight piglets show improved cognitive performance in the spatial cognitive holeboard task. Frontiers in behavioral neuroscience 9, 10. https://doi.org/10.3389/fnbeh.2015.00043

**H06:** Antonides, A., van Laarhoven, S., van der Staay, F.J., Nordquist, R.E., 2016.
Non-anemic iron deficiency from birth to weaning does not impair growth or memory in piglets. Frontiers in behavioral neuroscience 10:112. https://doi.org/10.3389/fnbeh.2016.00112

**H07:** Fijn, L., Antonides, A., Aalderink, D., Nordquist, R.E., van der Staay, F.J., 2016.
Does litter size affect emotionality, spatial learning and memory in piglets? Applied animal behaviour science 178, 23–31. https://doi.org/10.1016/j.applanim.2016.02.011

**H08:** Grimberg-Henrici, C.G., Vermaak, P., Bolhuis, J.E., Nordquist, R.E., van der Staay, F.J., 2016.
Effects of environmental enrichment on cognitive performance of pigs in a spatial holeboard discrimination task. Animal cognition 19, 271–283. https://doi.org/10.1007/s10071-015-0932-7

**H09:** van der Staay, F.J., Schoonderwoerd, A.J., Stadhouders, B., Nordquist, R.E., 2016.
Overnight social isolation in pigs decreases salivary cortisol but does not impair spatial learning and memory or performance in a decision-making task. Frontiers in veterinary science - animal behavior and welfare 2, 13. https://doi.org/10.3389/fvets.2015.00081

**H10:** Roelofs, S., Nordquist, R.E., van der Staay, F.J., 2017.
Female and male pigs' performance in a spatial holeboard and judgment bias task. Applied animal behaviour science 191, 5–16. https://doi.org/10.1016/j.applanim.2017.01.016

**H11:** Roelofs, S., van Bommel, I., Melis, S., van der Staay, F.J., Nordquist, R.E., 2018.
Low birth weight impairs acquisition of spatial memory task in pigs. Frontiers in veterinary science - animal behavior and welfare 5:142, 14. https://doi.org/10.3389/fvets.2018.00142

**H12:** Witjes VL, Roelofs S, Meijer E, Eicher PHC, Zeinstra EC, Oei CHY, et al.
Comparison of first- and last-born pigs revealed no effect of the birth process on acquisition and reversal of the cognitive holeboard task. Applied animal behaviour science. 2025;285, 106585:10.

#### Codes "Studies"

'H01'= '01: (Gieling et al., 2012)'
'H02'= '02: (Gieling et al., 2013)'
'H03'= '03: (Gieling et al., 2014)'
'H04'= '04: (Antonides et al., 2015b)'
'H05'= '05: (Antonides et al., 2015a)'
'H06'= '06: (Antonides et al., 2016)'
'H07'= '07: (Fijn et al., 2016)'
'H08'= '08: (Grimberg-Henrici et al., 2016)'
'H09'= '09: (van der Staay et al., 2016)'
'H10'= '10: (Roelofs et al., 2017)'
'H11'= '11: (Roelofs et al., 2018)'
'H12'= '12: (Witjes et al., 2018)'

#### Codes "Birthweight"

'vLBW'= 'very Low Birth Weight'
'LBW'= 'Low Birth Weight'
'NBW'= 'normal birth weight'

#### Codes "Sex"

'M' = 'Male'

'F' =	'Female'

### Codes "Breeds"

1=	'(Terra X Finn. Landr.) X Duroc'
2=	'Duroc X (Yorkshire X Finn. Landr.)'
3=	'Göttingen miniature pig'
4=	'(Terra X Dutch Landrace) X Duroc'
5=	'T40 X Pietrain'
6=	'Large White X PIC426'

### Codes "Treatment"

'barren'=  'Barren environment'
'enr.'= 'Enriched environment'
